# A network analysis for the identification of gene modules in the transcriptome during *Nicotiana benthamiana* interfamily grafting

**DOI:** 10.64898/2026.01.26.701652

**Authors:** Frank Opoku-Agyemang, Ken-ichi Kurotani, Michitaka Notaguchi

**Affiliations:** Graduate School of Bioagricultural Sciences, Nagoya University, Furo-cho, Chikusa-ku, Nagoya 464-8601, Japan; Bioscience and Biotechnology Center, Nagoya University, Furo-cho, Chikusa-ku, Nagoya 464-8601, Japan; Department of Botany, Graduate School of Science, Kyoto University, Kitashirakawa Oiwake-cho, Sakyo-ku, Kyoto 606-8502, Japan; National Key Laboratory for Germplasm Innovation and Utilization for Fruit and Vegetable Horticultural Crops, College of Horticulture and Forestry Sciences, Huazhong Agricultural University, Wuhan 430070, Hubei Province, China

**Keywords:** Bayesian network analysis, Gene modules, Hub genes, Interfamily grafting, Plant grafting, Transcriptome

## Abstract

**Background:** *Nicotiana benthamiana* has been found to exhibit grafting capability with phylogenetically distant plant species by accomplishing cell-cell adhesion as the first step in graft establishment. Morphological and physiological studies combined with time-course transcriptome analysis revealed that this grafting triggered various biological processes. Thus, further elucidation of accumulated datasets is required to describe the processes during grafting.

**Results:** In this study, we performed a Bayesian network analysis to identify crucial gene modules in the transcriptome of *Nicotiana benthamiana* interfamily grafting. Our bioinformatics analyses of the transcriptome included threshold-based clustering, functional annotation, Bayesian network analysis, module analysis, and hub gene identification. We defined six distinct temporal gene expression patterns in the transcriptome data. Gene ontology enrichment was performed for each expression pattern, and results were summarized as Gene ontology supercluster treemaps. Bayesian gene networks were constructed using the SiGN-BN HC + BS program along with 120 *N. benthamiana* transcriptome data covering the whole life cycle. Gene modules were identified using two module detection algorithms: Molecular Complex Detection (MCODE) and GLay. Gene modules were characterized by identifying Gene Ontologies and hub genes as the most interconnected nodes in the gene network using the Cytohubba plugin.

**Conclusion:** This study provides further knowledge and enhances our understanding of the molecular mechanisms underlying interfamily grafting.

## Introduction

Grafting is an ancient propagation technique that is used to join cut stems to establish a single plant following successful vascular adhesion and graft union formation [1–3]. It has been used extensively in horticulture to propagate vegetables, ornamentals, and fruit trees with desirable traits such as disease resistance, stress resilience, and improved yield and quality [2,4]. Graft success is dependent on a series of union development processes, including wound response, cell-cell adhesion, callus proliferation, and vascular formation at the graft junction. Successful accomplishment of these processes ensures strong adhesion between the rootstock and scion with subsequent scion outgrowth, flowering, and/or fruiting [5–7]. The practice of intrafamily and phylogenetically closely related plant grafting has been the most successful, thereby limiting the scope of graft combinations available [4,8–11], even though several interfamily graft combinations have been reported [12–15]. For example, Solanaceae plants are usually grafted with tomato (*Solanum lycopersicum*) for improved yield, disease resistance, and fruit quality [7].

However, recent studies on interfamily partner accepting grafts (iPAG) reported the successful graft establishment of *N. benthamiana* and *Petunia hybrida* scion (Solanaceae) with phylogenetically distant plant species, including *Arabidopsis thaliana* stock (Brassicaceae) [16] and *Chrysanthemum morifolium* (Asteraceae) [17,18] *N. benthamiana* has been reported to establish interfamily grafting with a wide range of evolutionary plant species and families from 73 species and 38 families, including magnoliids, monocots, eudicots, vegetable, flower, and fruit tree crops. These findings have significantly expanded successful graft combinations across diverse plant families. Morphological and physiological studies during *N. benthamiana*/*A. thaliana* interfamily grafting described biological processes during tissue reunion at graft wound sites [17]. Although cell-cell adhesion is a characteristic of interspecies grafting, other biological processes are less compatible. Therefore, the prolonged tissue reunion in interfamily grafts enables the capture of related biological events.

To evaluate the cellular mechanisms and elucidate the molecular events during interfamily grafting, transcriptome analyses have been conducted on interfamily graft combinations, including *N. benthamiana*/*A. thaliana* [17] and *P hybrida*/*A. thaliana* [18]. Particularly, in [17], graft-associated genes, including *IAA1* [19], *WOX4* [20], *VND7* [21], *ANAC071* [22], *TMO6* [23], and *OPS* [24] involved in auxin action, wound repair, and cambium and vascular development were identified as upregulated genes in response to *N. benthamiana*/*A. thaliana* interfamily grafting. Furthermore, GO terms for the 189 early upregulated genes identified in the study revealed that the extracellular region, cell wall, and apoplast were the top terms for these genes. Transcriptome analysis of *P. hybrida*/*A. thaliana* interfamily graft also revealed the expression of conventional graft-related genes. Specifically, graft-related genes, including *PIN1*, *ANAC071*, *WOX4*, *PLL1*, *VND7*, and *OPS*, which are involved in cell proliferation, wound repair, cambium activity, and xylem development, were identified [18]. Moreover, a study on xylem formation at the graft junction following transcriptome analyses of *N. benthamiana*/*A. thaliana* interfamily grafting [25] revealed upregulated expression patterns of known xylem-associated genes, four homologs of *VND7,* which have been reported in *A. thaliana* to be major transcription factors for xylem formation [26–28].

Consistent with the hypothesis that gene network analysis has been a powerful approach employed to elucidate the molecular mechanisms underlying plant grafting, recent studies on the efficient curation of co-regulated genes in the graft transcriptome based on gene regulatory network analysis have been reported [25,29]. Weighted gene co-expression network analysis (WGCNA) [30], on the transcriptome of internode and stem sections from auto-grafted tomato, revealed gene co-expression networks related to graft healing and union formation [29]. Regulatory gene network analysis [31–33] of compatible and incompatible grafts revealed *SlWOX4* as a potential regulator of graft compatibility [7]. Moreover, transcriptome and Bayesian gene network analyses using the SiGN-BN HC + BS program, implemented on a supercomputer system [34,35] of *N. benthamiana*/*A. thaliana* interfamily grafting revealed gene modules for tracheary element formation, including *XYLEM CYSTEINE PROTEASE* (*NbXCP*) genes. The module was centered around the *NbVND7* transcription factors, and co-regulated genes involved in xylem cell differentiation and immune response were identified [25]. In our recent study on the roles of four *NbVND7* homolog genes in grafting [36], the SiGN-BN HC + BS program was applied to estimate their gene regulatory networks using 36 RNA-seq data of *N. benthamiana* grafting. The program found the different modes of gene regulation among *VND7* homolog genes, and these regulatory modes were confirmed by observing the distinct gene expression patterns of *VND7* homologs in the intact and grafted stem tissues. These studies demonstrated that the SiGN-BN HC + BS program is very efficient in identifying previously known and unknown gene networks for each biological process. Therefore, we applied this Bayesian gene network analysis.

In this study, we investigated the molecular mechanisms and provided new insights underlying interfamily grafting through gene regulatory network estimations and analysis. RNA-seq data collected by [17,37] were used in this study. Data processing and gene expression calculation were performed for the dataset. Six distinct temporal gene expression patterns of the *N. benthamiana* scion in the *N. benthamiana*/*A. thaliana* interfamily graft transcriptome was identified following distance threshold-based clustering. GO enrichment analysis was performed for each expression pattern and summarized into treemaps of GO superclusters. Next, we estimated and constructed Bayesian networks to reveal the causal relationship between gene expression in each of the patterns using the SiGN-BN HC + BS program along with 120 *N. benthamiana* transcriptome data presented in [37]. We then identified gene modules using network module algorithms and hub genes, which were highly connected genes in each gene network. In reference to the hypothesis of our study, we have identified crucial gene modules and hub genes following gene network analysis during *N. benthamiana*/*A. thaliana* interfamily grafting. Finally, we performed a GO enrichment analysis to highlight the major biological processes during grafting as impacted by each gene module. The identification of hub genes that play key roles in plant grafting and their associated gene modules will provide a new basis and direction for grafting. Thus, two aims of the present study are as follows: (1) to investigate the molecular events and provide new insights into various biological processes involved in interfamily grafting through Bayesian gene regulatory network estimation and analysis, and (2) to identify gene modules, key functional annotations, and hub genes as targets for novel grafting strategies. The findings of this study may provide new insight into interfamily grafting at the transcriptome level and enhance our understanding of the molecular, cellular, and biological processes for novel grafting strategies.

## Results

### Setting of grafting states based on transcriptome data

The overall bioinformatics analysis workflow is shown in Fig. 1A. The RNA-seq data set from 21 *N. benthamiana* samples, comprising 3 *N. benthamiana* intact plants and 18 *N. benthamiana*/*A. thaliana* interfamily grafts from six time points, 1, 3, 5, 7, 14, and 28 days after grafting (DAG), obtained previously in [17], were analyzed using the *N. benthamiana* genome dataset, Nbe.v1.1 [37]. The gene expression levels and transcripts per million (TPM) count values were obtained. Then principal component analysis (PCA) was performed on the gene expression data set to assess the relationships of the biological replicates and to determine the extent of distant thresholds of intact compared to grafted samples, and according to the PCA, PC1 and PC2 accounted for 48% and 20% of gene expression variation, respectively. Based on the PCA, we divided the samples into three state changes according to their sampling timepoints post grafting, and based on the clustering of the biological replicates, they were named, ground state (intact samples), second state (1, 3, 5 and 7 DAG samples) and third state (14 and 28 DAG samples) (Fig. 1B). The concise clustering of the biological replicates at specific time points revealed that the sequential changes in gene expression appeared as time passed after interfamily grafting.

**Figure 1.**
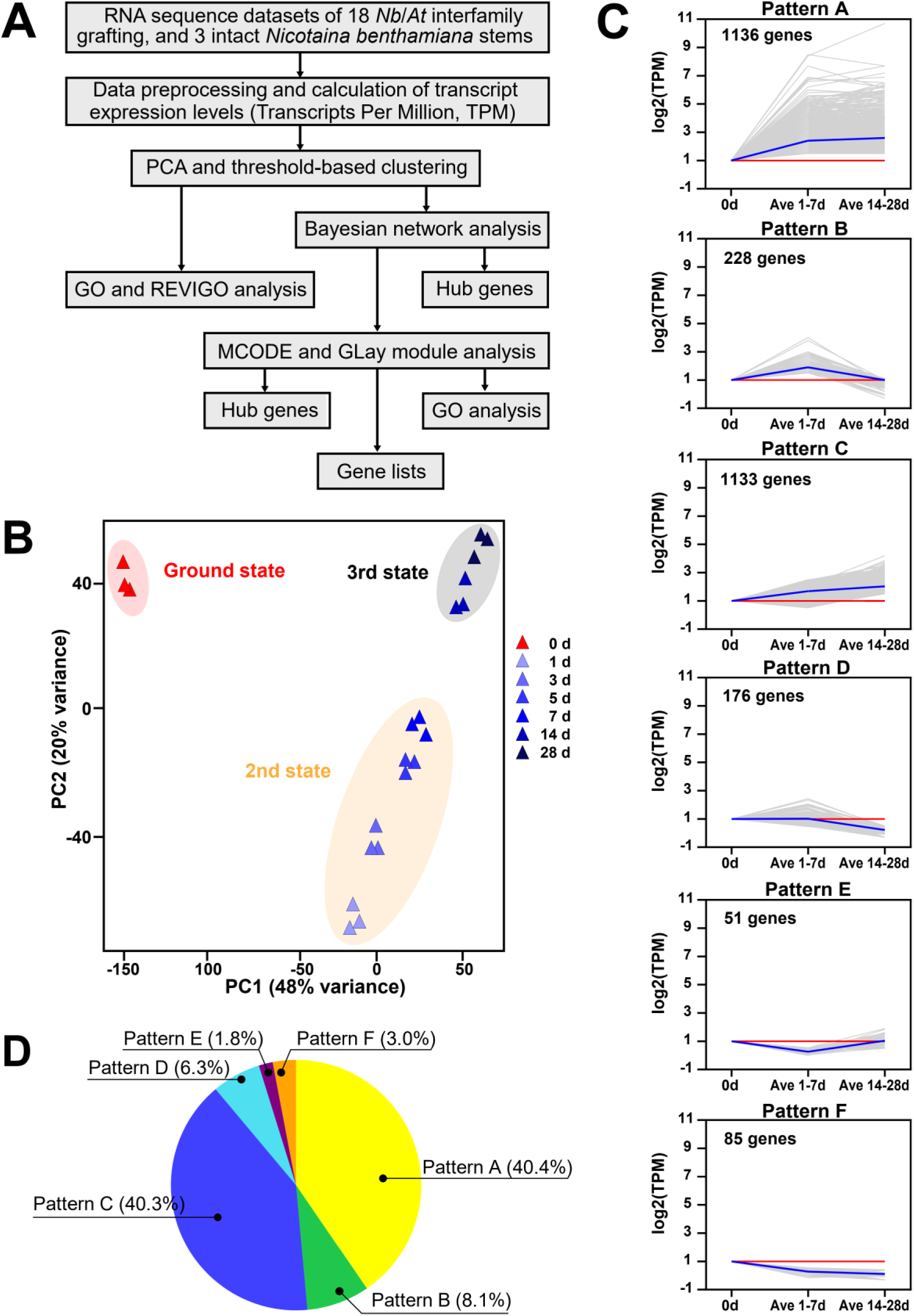
Classification of distinct temporal gene expression patterns in the transcriptome data of *N. benthamiana*/*A. thaliana* interfamily grafting. **(A)** Overview of the bioinformatics analysis pathway. **(B)** Principal component analysis (PCA) of the expression profiles of the *N. benthamiana* intact and the *N. benthamiana*/*A. thaliana* interfamily grafts at seven time points (three biological replicates per time point) and classification of the distinct gene expression patterns in the RNA-seq data set. Note: The groups are distinguished by different colors and shapes. The PCA revealed that PC1 and PC2 contributed 48% and 20% of the total variance, respectively. PCA plot showed the tight clustering of biological replicates, indicating a good correlation between the biological repeats and distinct clustering between groups, revealing three state changes, including ground state (intact samples), second state (1, 3, 5, and 7 DAG samples), and third state (14 and 28 DAG samples). **(C)** The six gene expression patterns in the *N. benthamiana* scions across different time points after interfamily grafting. The six patterns were identified from the distance threshold-based clustering analysis based on the gene expression data set. Gray lines indicate the relative expression levels of each gene in the pattern. Blue line indicates the mean relative expression in each pattern. The red line indicates a baseline value of 1 log_2_TPM. The x-axis is samples, and the y-axis is the transformed relative expression level. The gene number is marked on each pattern in the upper left corner. **(D)** Number of genes in the distribution of each gene expression pattern type.

### Classification of the distinct temporal gene expression patterns

We classified the expression patterns of the *N. benthamiana*/*A. thaliana* interfamily gene expression dataset into six distinct temporal gene expression categories by calculating their transcript abundance relative to the intact samples and according to the state changes revealed by the PCA (Fig. 1B). Using the log_2_ transformed TPM values, 3 data points classification was set including, ground state (intact samples with a baseline value of 1 log_2_TPM), an average of the log_2_TPM values of the second state including 1, 3, 5 and 7 DAG samples represented by “Ave 1–7 DAG” in the data presentation, and an average of the log_2_TPM values of the third state including 14 and 28 DAG samples represented by “Ave 14–28 DAG” in the data presentation. Setting the above classification with a baseline ensured an interpretable visualization of gene expression across the different time points and also focused on the genes with biological relevance during the *N. benthamiana*/*A. thaliana* interfamily grafting. Six distinct gene expression patterns were defined following the distance threshold-based clustering and the changing expression profiles of genes across the different time points post-grafting relative to the expression levels in intact samples (Fig. 1C). Expression pattern A genes were predominantly highly expressed in the second (Ave 1–7 DAG) and third states (Ave 14–28 DAG), pattern B genes were predominantly highly expressed in the second state (Ave 1–7 DAG) only, and pattern C genes were predominantly highly expressed in the third state (Ave 14–28 DAG) only (Fig. 1D). Expression patterns A and C were the most abundant in the gene expression data, accounting for approximately 40% each among differentially expressed genes (Fig. 1D). Patterns D–E were downregulated in the opposite manners of patterns A–C as follows; patterns D genes showed lower expression levels in the third state (Ave 14–28 DAG) only, pattern E genes had lower expression levels in the second state (Ave 1–7 DAG) only, and pattern F genes showed lower expression levels in the second (Ave 1–7 DAG) and third states (Ave 14–28 DAG) (Fig. 1D). Among upregulated genes in patterns A–C, genes involved in various cellular/biological processes were identified. Pattern A revealed graft-associated genes known to be involved in key grafting processes including glycosyl hydrolases (β-1,3-glucanase), pectin methylesterase inhibitors (*PMEI*, *PMEI13*), NAC domain-containing proteins (*NAC007*, *NAC060*, *NAC073*, *NAC053*, *NAC036*, *NAC04*, *NAC02*, *VND1* and *VND7*), wound-responsive proteins (*WIP3* and *BBD2*), LOB domain-containing protein (*LBD4*), plasmodesmata protein (*PDLP2*), auxin-related proteins (*PIN3*, *PIN7*, *FL8*, *IAA17* and *DFL1*), brassinosteroid-related protein (*BRL1*), ethylene-responsive factors (*ERF1* and *ERF106*), sugar transporter (*STP14*) and homeobox proteins (*HAT14* and *KNAT7*) (Fig. 1C and Supplementary Dataset S1: Sheet 1). Genes in pattern B included previously known graft-associated genes such as IRREGULAR XYLEM 15-Like (*IRX15-L*), β-xylosidase 6 (*BXL6*), NAC domain-containing proteins (*NAC007* and *NAC060*), myb domain protein 83 (*MYB83*), peroxidase 17 (*PRX17*), basic helix-loop-helix DNA-binding protein (bHLH), LOB domain-containing protein 15 (*LBD15*), cyclin D2;1/cyclin A2;2 (*CYCD2;1*/*CYCA2;2*), and amine oxidase 1 (*AO1*) (Fig. 1C and Supplementary Dataset S1: Sheet 2). Genes identified in pattern C had the following previously known graft-associated genes including NAC domain-containing proteins (*NAC002*, *NAC007*, *NAC004*, *NAC029*, *NAC053* and *NAC060*), auxin efflux carrier family protein (*PIN3* and *PIN7*), basic helix-loop-helix (bHLH) DNA-binding superfamily protein (*bHLH121*), auxin canalization protein (*FL8*), BES1-interacting Myc-like protein 2 (*BES1*), homeobox leucine zipper protein (*HAT14*), wound-responsive family protein (*WIP3*), early nodulin-like protein 9 (*ENODL9*), rhamnogalacturonate lyase family protein, mitotic-like cyclin 3B (*CYC3B*) and cytokinin response factor 4 (*CRF4*) (Fig. 1C and Supplementary Dataset S1: Sheet 3). Compared with patterns A–C, genes belonging to patterns D, E, and F were scarce (Fig. 1C and Supplementary Dataset S1: Sheets 4, 5, and 6, respectively). Hereafter, the genes identified in all distinct expression patterns were employed for functional annotation and Bayesian gene regulatory network analysis to examine the molecular mechanism underlying *N. benthamiana*/*A. thaliana* interfamily grafting.

### Functional categorization of genes in each temporal gene expression pattern by Gene Ontology enrichment analysis

We performed GO enrichment analysis using GOATOOLS [38] to explore the relevant biological functions induced by the genes extracted from each specific expression pattern. For each expression pattern, the significantly enriched GO terms (*p* < 0.01) in the biological process (BP) category were screened. The long, redundant, and unintelligible lists of GO terms were summarized and visualized as treemaps using the REVIGO tool [39] following a simple clustering algorithm that relies on semantic similarity measures. The GO enrichment in the BP category revealed that the expression patterns share several superclusters. The “Organic cyclic compound metabolic process” supercluster appeared in expression patterns B, D, and F, while the “RNA modification” supercluster appeared in expression patterns C, D, and F. They are the only two superclusters shared by two or more expression patterns. Nine superclusters were summarized for 76 significant biological processes for genes in expression pattern A, as “DNA recombination” (including 17 terms), “DNA damage response” (including 9 terms), and “regulation of response to water deprivation” (including 8 terms) (Fig. 2A and Supplementary Dataset S2: Sheet 7). The 121 significantly enriched biological processes in expression pattern B were integrated into 9 superclusters, such as “negative regulation of mitotic cell cycle” (including 20 terms), “nucleic acid phosphodiester bond hydrolysis” (including 12 terms), and “fucose catabolic process” (including 10 terms) (Fig. 2B and Supplementary Dataset S2: Sheet 8). The expression pattern C had 90 significant biological processes summarized into 12 superclusters, including “RNA modification” (including 28 terms), “DNA damage response” (including 16 terms), and “Positive regulation of chromosome organization” (including 4 terms) (Fig. 2C and Supplementary Dataset S2: Sheet 9). Seven superclusters were classified for the 25 significantly enriched biological processes identified for the gene in expression pattern D, as “RNA modification” (including 10 terms), “Organic cyclic compound metabolic process” (including 6 terms), and “regulation of primary metabolic process” (including 2 terms) (Fig. 3A and Supplementary Dataset S2: Sheet 10). The minimum number of superclusters, 2, were summarized for the 7 significantly enriched biological processes identified in expression pattern E, including “auxin biosynthetic process” (including 5 terms), and “histone H3-K9 acetylation” (including 2 terms) (Fig. 3B and Supplementary Dataset S2: Sheet 11). Seven superclusters were summarized for 58 significant biological processes identified in expression pattern F, including “RNA modification” (including 17 terms), “regulation of intracellular signal transduction” (including 10 terms), and “organic cyclic compound metabolic process” (including 6 terms) (Fig. 3C and Supplementary Dataset S2: Sheet 12).

**Figure 2.**
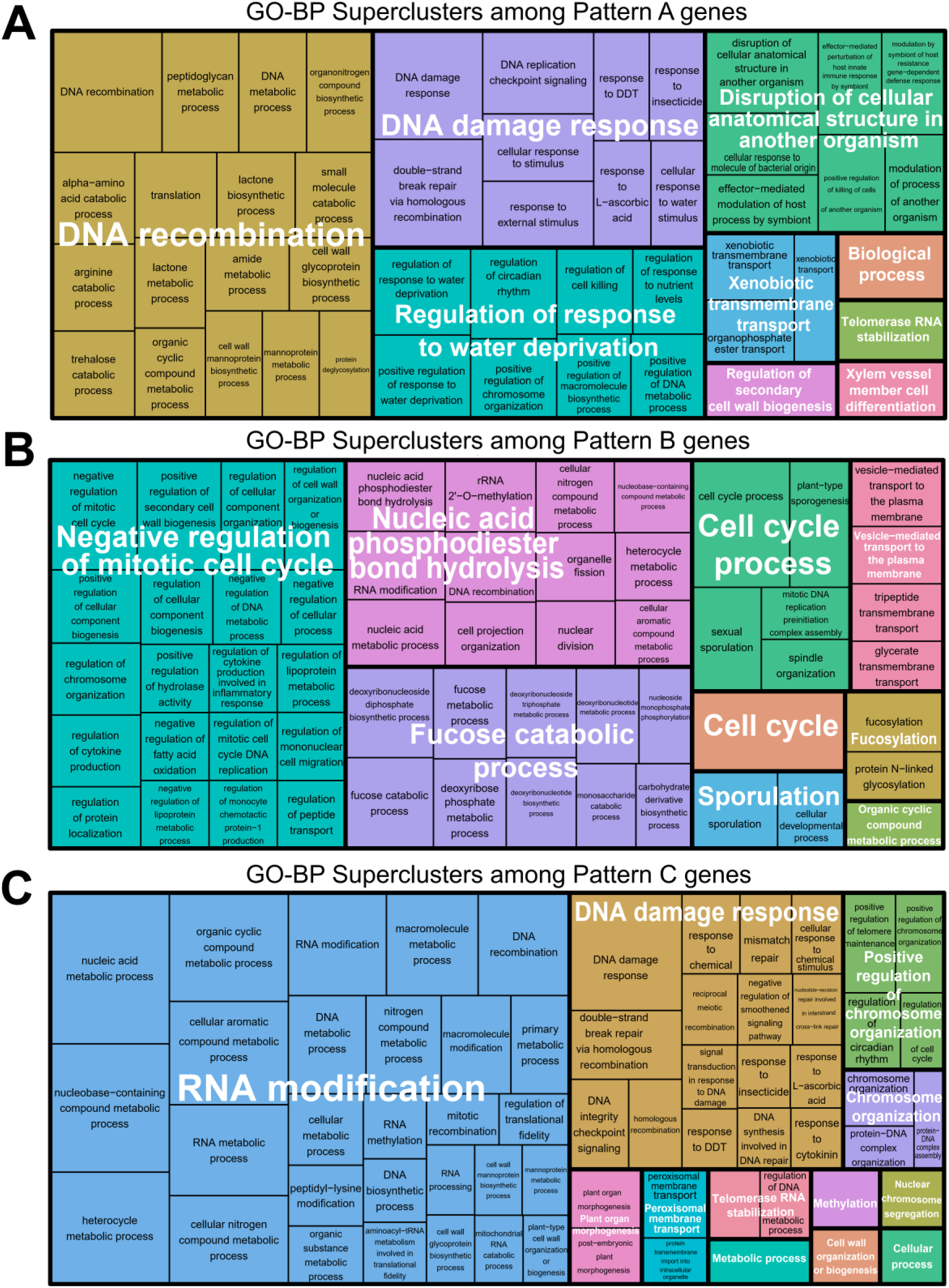
REVIGO treemap of GO-BP categories among genes in expression patterns A–C. Treemap views of summarized GO-BP categories were visualized using a simple clustering algorithm that relies on semantic similarity measurements. **(A)** Nine superclusters were summarized for 76 GO-BPs enriched in Pattern A. **(B)** Nine superclusters were summarized for 121 GO-BPs enriched in Pattern B. **(C)** Twelve superclusters were summarized for 90 GO-BPs enriched in Pattern C. Each rectangle is a single representative significant GO term joined into superclusters of loosely related terms and visualized with distinct colors, with supercluster names labeled in white fonts. The size of the rectangles is proportional to the p-value given by the GOATOOLS analysis, such that the larger the rectangle, the more significant the GO-term. Corresponding data are presented in Supplementary Dataset S2: Sheets 7–12, for details.

**Figure 3.**
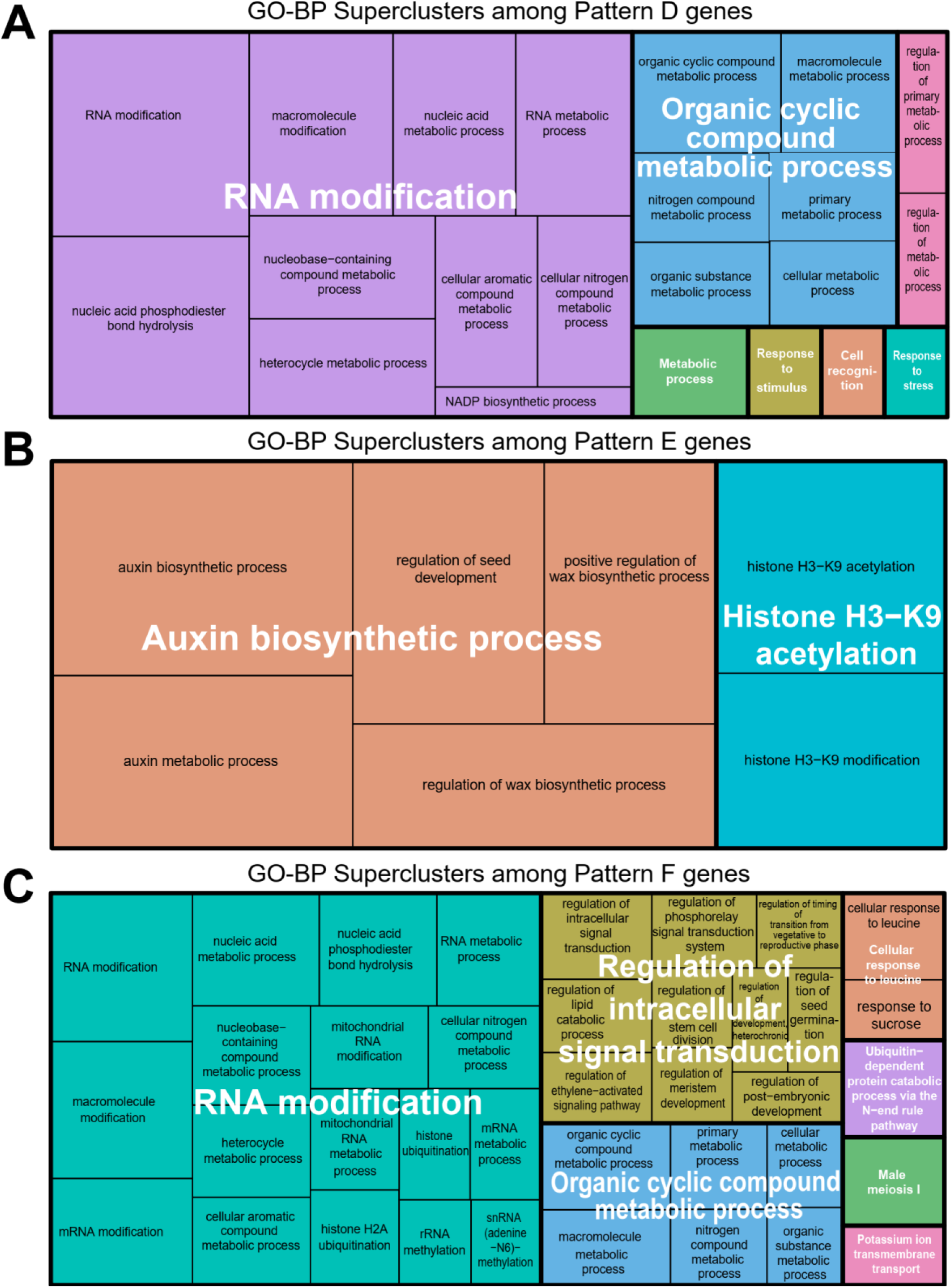
REVIGO treemap of GO-BP categories among genes in expression patterns D–F. Treemap views of summarized GO-BP categories were visualized using a simple clustering algorithm that relies on semantic similarity measures. **(A)** Seven superclusters were summarized for 25 GO-BPs enriched in pattern D. **(B)** Two superclusters were summarized for 7 GO-BPs enriched in pattern E. **(C)** Seven superclusters were summarized for 58 GO-BPs enriched in pattern F. Each rectangle is a single representative significant GO term joined into superclusters of loosely related terms and visualized with distinct colors, with supercluster names labeled in white squares. The size of the rectangles is proportional to the p-value given by the GOATOOLS analysis, such that the larger the rectangle, the more significant the GO-term. Corresponding data are presented in Supplementary Dataset S2: Sheets 10–12, for details.

### Gene network analysis and hub gene identification

To investigate the gene regulatory network and estimate regulatory dependencies between genes extracted from each distinct expression pattern, we employed Bayesian network construction using the SiGN-BN HC + Bootstrap program [34,35] implemented on our cluster server system. The 120 independent *N. benthamiana* transcriptome data, acquired using a consistent methodology in our previous study [37], were used in this study and enabled comparisons for estimating the gene regulatory networks. These data included the following categories: (1) a time-series of *N. benthamiana*/*A. thaliana* interfamily grafting samples collected at 2 hours after grafting and 1, 3, 5, 7, 10, 14, and 28 DAG along with intact controls (originally collected in [17] and reprocessed in [37]; (2) *N. benthamiana*/*N. benthamiana* homografting samples collected at 1, 3 and 7 DAG along with intact controls (originally collected in [17] and reprocessed in [37]); (3) various *N. benthamiana* tissues and organs including the shoot apices, cotyledons, hypocotyls, and roots at the seedling stage and young and mature leaves, flowers, stems, and roots at the mature plant stage [37] and (4) stem segments collected at five or six defined positions along the axis sampled 7 days after either wounding or grafting treatment, respectively [37]. The range of genes included in the pattern A–F in this study (51–1,136 genes) met the extreme input limit of this program (about 1,000 genes) and was optimum for network construction. The Bayesian network for genes in all six gene expression patterns was constructed and visualized by Cytoscape [40] (Fig. 4). The gene networks of patterns A and C were the largest, with 1,136 nodes; 19,331 edges, and 1,133 nodes; 18,376 edges, respectively (Fig. 4A, 4C, and Supplementary Dataset S3: Sheet 13 and 15). Patterns B and D had a much bigger network of 228 nodes; 3,138 edges and 176 nodes; 2,227 edges, respectively (Fig. 4B, D and Supplementary Dataset S3: Sheet 14 and 16) while patterns E and F had smaller gene networks with 51 nodes; 586 edges and 85 nodes; 991 edges, respectively (Fig. 4E, F and Supplementary Dataset S3: Sheet 17 and 18). The comparative topological parameters of the core networks of each expression pattern analyzed using the Network Analyzer Plugin in Cytoscape [41,42] are presented in Table 1. They include degree distribution, clustering coefficients, centrality, distribution of node degrees, neighborhood connectivity, and average shortest path lengths. Next, we used the CytoHubba plugin [43] to identify hub genes that may play crucial roles in plant grafting in the gene regulatory networks for each expression pattern. In this study, hub genes are described as genes with a higher degree of connectivity in a network. Hub genes were nodes with degree values greater than or equal to 0.95 quantiles (ranked top 5%) of the calculated degree distribution.

**Figure 4.**
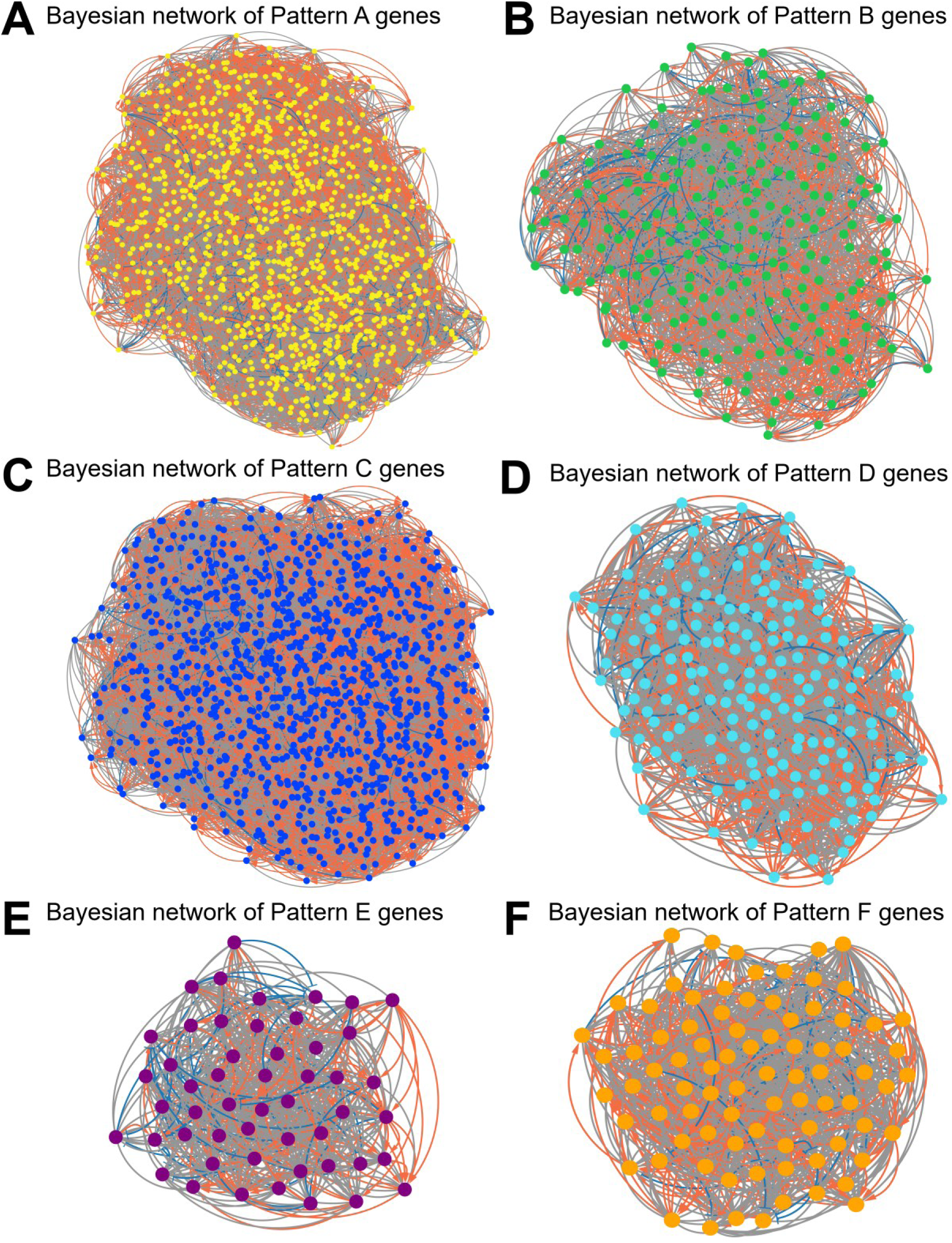
Bayesian network analysis of genes in expression patterns A–F. **(A)** Network diagram with only the nodes and edges for genes extracted from pattern A indicated by yellow dots and arrows/lines, respectively. **(B)** Network diagram with only the nodes and edges containing genes extracted from pattern B indicated by green dots and arrows/lines, respectively. **(C)** Network diagram with only the nodes and edges containing genes extracted from pattern C indicated by blue dots and arrows/lines, respectively. **(D)** Network diagram with only the nodes and edges containing genes extracted from pattern D indicated by turquoise dots and arrows/lines, respectively. **(E)** Network diagram with only the nodes and edges containing genes extracted from pattern E indicated by purple dots and arrows/lines, respectively. **(F)** Network diagram with only the nodes and edges containing genes extracted from pattern F indicated by orange dots and arrows/lines, respectively. Red arrows indicate gene up-regulation. Blue T-lines indicate gene down-regulation. Gray lines indicate uncertain gene regulation. Corresponding data are presented in Supplementary Dataset S3: Sheets 13–18.

**Table 1.** Topological parameters analysis of the core Bayesian networks of the expression patterns A–F.

| Network parameters | A | B | C | D | E | F |
| --- | --- | --- | --- | --- | --- | --- |
| Number of nodes | 1,136 | 228 | 1,133 | 176 | 51 | 85 |
| Number of edges | 19,331 | 3,128 | 18,376 | 2,227 | 586 | 984 |
| Average number of neighbors | 34 | 27 | 32 | 25 | 23 | 23 |
| Network diameter | 5 | 4 | 5 | 4 | 2 | 3 |
| Network radius | 3 | 3 | 3 | 3 | 2 | 2 |
| Characteristic path length | 2.689 | 2.097 | 2.656 | 2.026 | 1.540 | 1.741 |
| Clustering coefficient | 0.266 | 0.365 | 0.239 | 0.362 | 0.557 | 0.456 |
| Network density | 0.030 | 0.121 | 0.029 | 0.145 | 0.460 | 0.282 |
| Network heterogeneity | 0.375 | 0.240 | 0.380 | 0.245 | 0.192 | 0.259 |
| Network centralization | 0.063 | 0.096 | 0.069 | 0.114 | 0.209 | 0.242 |

These genes are the most significant in gene regulatory networks and may play an important role in each biological process included in the gene module. The network for the hub genes identified in each gene expression pattern and constructed using the CytoHubba plugin is presented in Fig. 5A–F. The color of nodes, from red to yellow, represents the degree of connectivity of nodes, with red being highly connected and yellow weakly connected. Detailed information on each hub gene, including ranks and names, is shown in Supplementary Dataset S4: Sheet 19–24. The expression patterns A, B, C, D, E, and F had 57, 11, 57, 57, 9, 3, and 4 hub genes, respectively.

**Figure 5.**
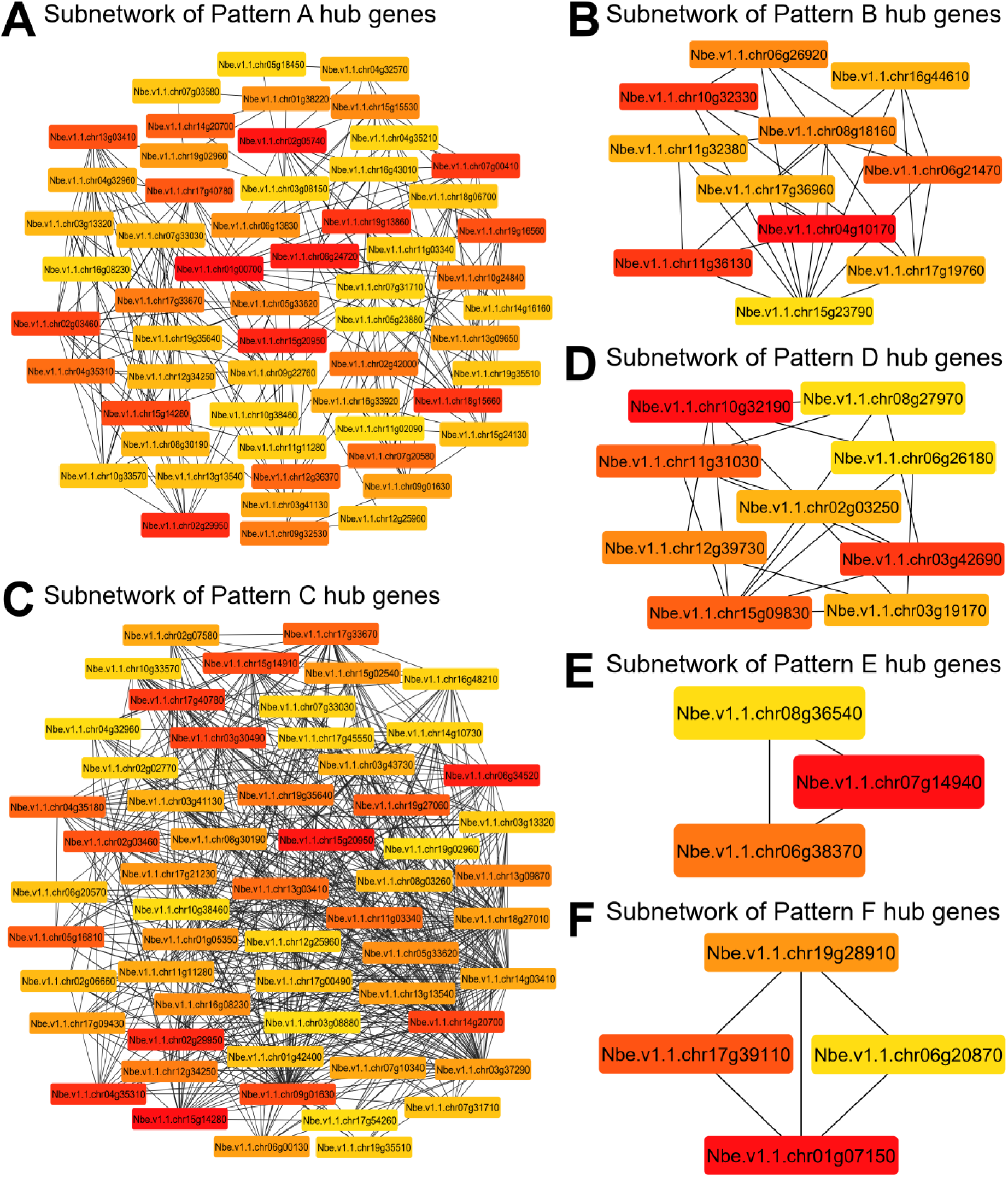
Hub gene identification in Bayesian gene networks for expression patterns A–F. **(A)** Subnetwork of the top 57 hub genes with the highest interaction degree ranked by the Degree algorithm in the expression pattern A. **(B)** Subnetwork of the top 11 hub genes with the highest interaction degree ranked by the Degree algorithm in the expression pattern B. **(C)** Subnetwork of the top 57 hub genes with the highest interaction degree ranked by the Degree algorithm in the expression pattern C. **(D)** Subnetwork of the top 9 hub genes with the highest interaction degree ranked by the Degree algorithm in the expression pattern D. **(E)** Subnetwork of the top 3 hub genes with highest interaction degree ranked by the Degree algorithm in the expression pattern E. **(F)** Subnetwork of the top 4 hub genes with highest interaction degree ranked by the Degree algorithm in the expression pattern F. The genes with the highest connections in each gene network, greater than or equal to the 0.95 quantiles of the degree distribution, are detected as hub genes by the CytoHubba Cytoscape plugin. The degree of importance of hub genes is depicted using the color scale ranging from red (high degree scores) to yellow (low degree scores). The detailed information of these hub genes is shown in Supplementary Dataset S4: Sheets 19–24.

The top hub genes identified in the gene regulatory network of expression pattern A, in which genes were continuously upregulated in the second and third states, include *Nbe.v1.1.chr01g00700* (a homolog gene encoding NB-ARC domain-containing disease resistance protein), a domain-containing disease resistance protein, and a nucleotide-binding adaptor involved in the regulatory core of the cell-death programmes and associated immunity responses in plants [44]. *Nbe.v1.1.chr02g05740* (a homolog gene encoding WRKY family transcription factor, *WRKY6*), a transcription factor that regulates the expression of phosphate1 (*PHOL1*) and is involved in phosphate stress tolerance and plant pathogen defense [45] and *Nbe.v1.1.chr06g24720* (a homolog gene encoding protein kinase superfamily protein, *RAF36*), which encodes a group C Raf-like protein kinase that is involved in the negative regulation of abscisic acid (ABA) responses during post-germination growth and plant resistance to pathogens [46].

In the gene regulatory network of expression pattern B, in which genes were upregulated in the second state only, the top hub genes identified include *Nbe.v1.1.chr04g10170* (a homolog gene encoding thaumatin-like protein 3, *TLP-3*), thaumatin-like proteins (TLPs) found in the class of pathogenesis-related (PR) proteins, and are involved in defense pathways and plant defense against fungal pathogens [47,48]. *Nbe.v1.1.chr10g32330* (a homolog of *CYCLIC NUCLEOTIDE-GATED CHANNEL 14*, *CNGC14*), which is a member of the cyclic nucleotide-gated channel family and localized in the plasma membrane, Ca^2+^ channel is involved in the regulation of auxin-mediated growth responses, gravitropism in plant roots, and mediates calcium influx required for tip growth in root hairs [49,50] and *Nbe.v1.1.chr11g36130* (a homolog gene encoding RNI-like superfamily protein) is involved in several plant biological processes, including SCF-dependent proteasomal ubiquitin-dependent protein catabolic process. It acts upstream of or within the ubiquitin-dependent protein catabolic process.

The top hub genes identified in the gene regulatory network of expression pattern C, in which genes were upregulated in the third state only, include *Nbe.v1.1.chr15g14280* (a homolog gene encoding zinc knuckle (CCHC-type) family protein), *Nbe.v1.1.chr15g20950* (a homolog gene encoding prolyl oligopeptidases (POPs) family protein), which is involved in the degradation of biologically important peptides [51] and *Nbe.v1.1.chr06g34520* (a homolog of tRNase Z4 gene, *TRZ4*), which encodes tRNase Z present in chloroplasts, and its deletion results in an embryo-lethal phenotype [52].

In the gene regulatory network of expression pattern D with downregulated genes in the third state only, the top hub genes included *Nbe.v1.1.chr10g32190* and *Nbe.v1.1.chr15g09830* (homolog genes encoding pentatricopeptide (PPR) repeat-containing proteins), which plays essential roles in organelle biogenesis, including constitutive and often essential roles in mitochondria and chloroplasts, through binding to organellar transcripts [53]; *Nbe.v1.1.chr03g42690* (a homolog gene encoding agenet domain (AGD)-containing protein 1, *AGDP1*), a heterochromatin-binding protein that binds to H3K9me2 tails and long transposable elements (TEs), was identified as a top hub gene in the pattern D and is involved in DNA methylation pathways and gene regulation [54,55].

The top hub genes identified in the gene regulatory network of expression pattern E, in which genes were downregulated in the second state, only include *Nbe.v1.1.chr07g14940* (a homolog of aldehyde oxidase 4 gene, *AAO4*), which codes for an aldehyde oxidase involved in delayed silique senescence via aldehyde detoxification by catalyzation, reactive oxygen species (ROS) generation, and response to water deprivation [56], *Nbe.v1.1.chr06g38370* (a homolog gene encoding RHOMBOID-like protein 12, *RBL12*), a serine protease class of mitochondrion-located rhomboid-like protein involved in mitochondrial processes and functional regulations [57], and *Nbe.v1.1.chr08g36540* (a homolog of UDP-glucosyl transferase 73B3 gene, *UGT73B3*), a UDP-glucosyl transferase 73B3 involved in the regulation of redox status, general detoxification of ROS-reactive secondary metabolites (SMs), and response to biotic and abiotic stresses [58,59].

The top hub genes identified in pattern F, with downregulated genes in the second and third states, include *Nbe.v1.1.chr01g07150* (a homolog gene encoding MUTL protein homolog 1, *MSH1*), which encodes a DNA-binding protein that promotes re-arrangements of the mitochondrial genome involved in abiotic stress tolerance, delayed maturity, and improved growth vigor of grafted plants [60], *Nbe.v1.1.chr17g39110* (a homolog gene encoding duplicated homeodomain-like superfamily protein, *GT2*), a plant transcriptional activator that contains trihelix DNA-binding domains involved in abiotic stress tolerance [61,62] and *Nbe.v1.1.chr19g28910* (a homolog gene encoding tetratricopeptide repeat (TPR)-like superfamily protein), which mediates protein-protein interactions in biological systems by binding to peptide ligands and is involved in the development and responses to light, phytohormones, and various stresses [63].

### Gene modules and hub genes identified in the Bayesian networks of six expression patterns

Two algorithms, MCODE [64] and GLay [65], were applied to identify the functional modules in the Bayesian networks of the six expression patterns A–F. We first utilized the gene module identification algorithm MCODE to perform module analysis in the gene regulatory networks of each gene expression pattern. This analysis was used to investigate the underlying biological mechanisms of *N. benthamiana*/*A. thaliana* interfamily grafting and reveal the core grafting-related processes by identifying modules that typically consist of genes that are highly interconnected and often share common biological functions or regulatory mechanisms. MCODE is a module clustering algorithm designed to detect tightly co-regulated gene sets or small dense modules of protein complexes in a gene regulatory network, leaving the majority of the nodes unclustered [64]. This algorithm uses a three-stage process of vertex weighting, complex prediction from high-weight nodes, and density filtering to detect protein complexes or functional modules. In a gene regulatory network like in this study, the detected modules represent co-regulated gene sets, and these genes may include transcription factors, enzymes, structural proteins, signaling proteins, transport and stress-related proteins, which imply their potential functions. The node with the highest weighted vertex, defined as the seed node, which serves as the origin of each MCODE, was identified. A threshold score > 5 was used to select significant modules in expression patterns A–F, respectively, and visualized with Cytoscape. To reveal the biological processes that the identified gene modules are associated with, the MCODE modules were evaluated for GO-enriched terms. The overrepresented biological processes in the module were evaluated using GOATOOLS [38]. For each MCODE module identified, nodes with a higher degree score over the 0.95 quantiles of the calculated degree distribution were designated as the hub genes, which indicate the overall biological significance of each module.

We then utilized the GLay community clustering algorithm implemented in Cytoscape for the identification of significant gene modules of gene expression patterns A–F. The GLay community clustering algorithm comprehensively and efficiently created large subnetworks by completely clustering and utilizing all the nodes in the large gene regulatory network with structured visualization. This algorithm was a method of choice for our analysis because it produces larger modules with greater sensitivity in recovering more biologically relevant information using all genes and implements the Girvan-Newman greedy algorithm to iteratively merge communities that maximize the modularity score, a metric measuring edge density within vs between communities [65,66]. GLay, particularly, was useful for the exploratory analysis and discovering broader functional relationships in the gene regulatory networks, while MCODE was more suited for identifying smaller, highly specific modules. Functionally related genes in each gene network, with the tendency of local neighborhood formation within large networks, were identified. GO enrichment analysis revealed that the gene networks from genes of expression patterns A–F were organized into several functionally distinct modules by the GLay algorithm and have relevant functional annotations associated with plant grafting. Next, for each GLay module identified, nodes with a higher degree score over 0.95 quantiles of the calculated degree distribution were designated as the hub genes, which indicate the overall biological significance of each module.

#### Pattern A

Using the MCODE algorithm, 15 modules (946 nodes, 2,258 edges, leaving 195 nodes unclustered) were identified in pattern A. There were 5 significant MCODE modules in pattern A with a threshold score of > 5, which included 412 nodes and 1,195 edges (Fig. 6 and Supplementary Dataset S5: Sheet 25). For pattern A, the modules were composed of various *N. benthamiana* homolog gene families and predominantly included transcription factors (such as MYB domain proteins, NAC domain-containing proteins, WRKY family transcription factor); enzymes (such as DNA glycosylase, UDP-glycosyltransferases, kinases); structural proteins (actin-related proteins, microtubule-associated proteins); signaling proteins (such as protein kinases, calcium-binding proteins); transport proteins (such as amino acid permeases, ABC transporters) and stress response proteins (such as heat shock proteins, disease resistance proteins).

**Figure 6.**
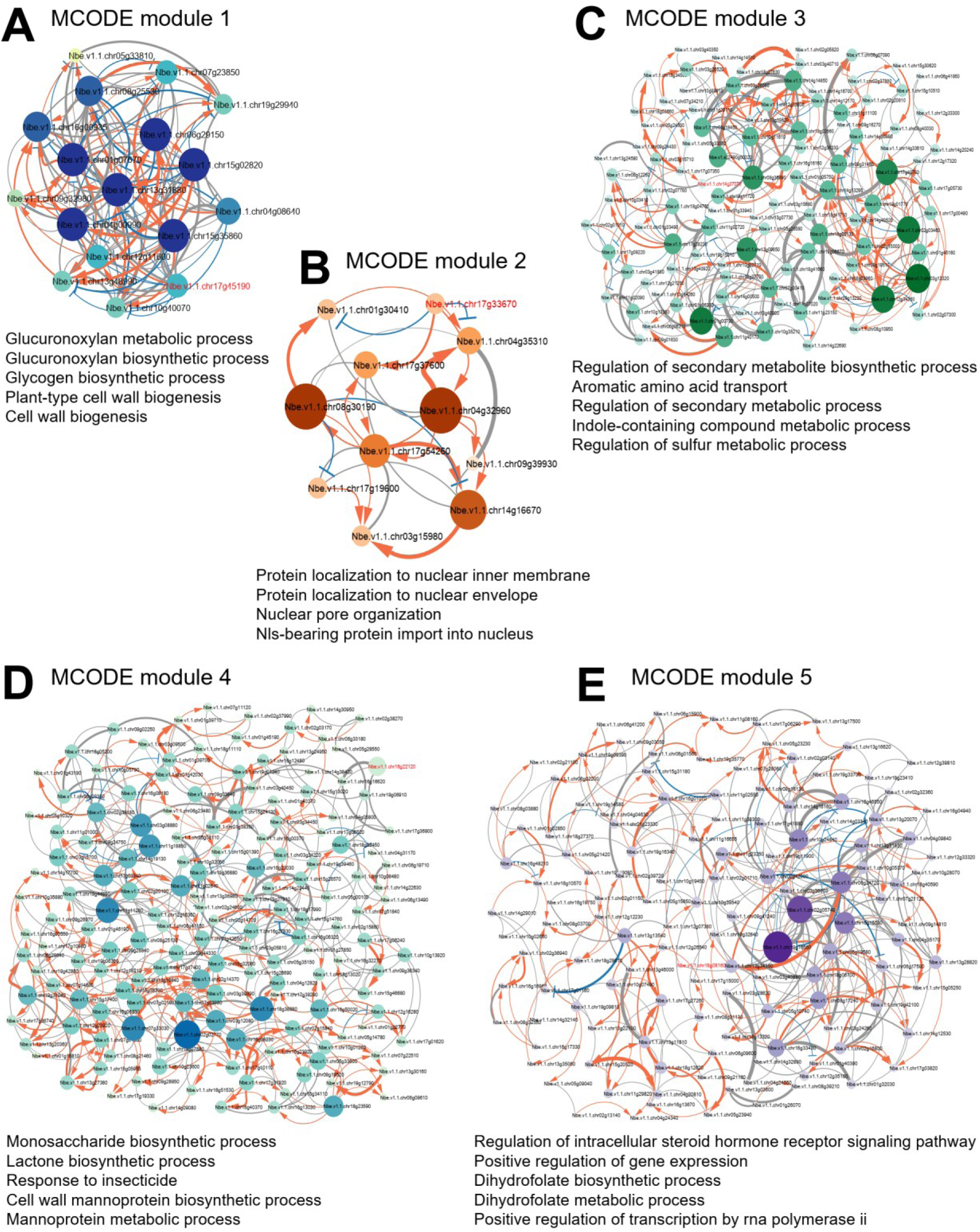
MCODE module analysis of the Bayesian network for expression pattern A. The module analysis using MCODE revealed 5 significant modules, which include 412 nodes and 1,195 edges, and the enrichment of several GO-BP terms. Modules 1–5 are the top five modules identified. **(A)** Module 1 (BTB/POZ complex) GO annotation has shown “glucuronoxylan metabolic process” and “cell wall biogenesis”. **(B)** Module 2 (Zinc finger complex) GO annotation has revealed “protein localization to nuclear inner membrane”, “nuclear envelope”, and “nuclear pore organization”. **(C)** Module 3 (UDP-glycosyltransferase 74 F1 complex) GO annotation has shown “secondary metabolite biosynthetic process” and “aromatic amino acid transport”. **(D)** Module 4 (O-fucosyltransferase complex) GO annotation has shown “monosaccharide biosynthetic process”, “lactone biosynthetic process”, and “response to insecticide”. **(E)** Module 5 (S-locus lectin protein kinase complex) GO annotation has “regulation of intracellular steroid hormone receptor signaling pathway”, “positive regulation of gene expression”, and “dihydrofolate biosynthetic process”. The seed node responsible for forming each module is indicated by red font. The module networks were visualized by Cytoscape (v3.10.2) by mapping the “degree parameter” to node size and color such that higher-degree nodes are darker in color and bigger in size. Red arrows indicate gene up-regulation. Blue T-lines indicate gene down-regulation. Gray lines indicate uncertain gene regulation. Line thickness indicates significance in gene regulation. All nodes are significantly enriched at p-value < 0.01. See Supplementary Dataset S5: Sheet 25 for details.

The following section describes the five significant MCODE modules (threshold score > 5) identified in the network of expression pattern A, which has genes predominantly highly expressed in the second (Ave 1–7 DAG) and third states (Ave 14–28 DAG). The seed node (central node with the strongest connections and highest weight score) in module 1 was *Nbe.v1.1.chr17g45190* (a homolog gene encoding BTB/POZ domain-containing protein) (Fig. 6A and Supplementary Dataset S5: Sheet 25), which plays a major role in regulating diverse biological processes related to plant growth, development, immunity, and stress adaptation [67]. Module 1 is mainly enriched in GO-BP terms associated with the “Glucuronoxylan metabolic process” and “cell wall biogenesis”. Hub genes in module 1 included *Nbe.v1.1.chr13g31880* (a homolog gene encoding core-2/I-branching beta-1,6-N-acetylglucosaminyltransferase family protein), *Nbe.v1.1.chr15g35860* (a homolog gene encoding plant glycogenin-like starch initiation protein 3, *PGSIP3*), *Nbe.v1.1.chr15g02820,* and *Nbe.v1.1.chr04g00990* (homolog genes encoding NAC domain-containing protein 73, *NAC073*), and *Nbe.v1.1.chr06g29150* (a homolog gene encoding RING/U-box superfamily protein, *ATL54*). The seed node of module 2 was *Nbe.v1.1.chr17g33670* (a homolog gene encoding zinc finger (CCCH-type) family protein, *TRM2B*) is involved in the stability and proper functioning of tRNA essential for protein synthesis [68,69] (Fig. 6B and Supplementary Dataset S5: Sheet 25). This module is mainly enriched in protein localization to the “nuclear inner membrane” and “nuclear pore organization” biological processes. Hub genes identified in module 2 were *Nbe.v1.1.chr04g32960* (a homolog gene encoding cysteine proteinases superfamily protein, *OTU5*) and *Nbe.v1.1.chr08g30190* (a homolog gene encoding RNA ligase/cyclic nucleotide phosphodiesterase family protein). The seed node of module 3 was *Nbe.v1.1.chr14g37070* (a homolog of UDP-glycosyltransferase 74 F1 gene, *UGT74F1*) (Fig. 6C and Supplementary Dataset S5: Sheet 25), which plays a key role in salicylic acid (SA) glycosylation, which regulates SA levels and plant immune responses [70–72]. This module was mainly enriched in GO-BP terms, “secondary metabolite biosynthetic process” and “aromatic amino acid transport”. Hub genes in module 3 included *Nbe.v1.1.chr01g00700* (a homolog gene encoding NB-ARC domain-containing disease resistance protein), *Nbe.v1.1.chr03g13320* (a homolog gene encoding hypothetical protein, *C53*), *Nbe.v1.1.chr12g34250* (a homolog gene encoding inner nuclear membrane protein B, *NEMP_B*), *Nbe.v1.1.chr02g03460* (a homolog gene encoding hypothetical protein 1, *OSCA2.3*) and *Nbe.v1.1.chr17g40780* (a homolog gene encoding TPR-like superfamily protein). The seed node of module 4 was *Nbe.v1.1.chr16g22120* (a homolog gene encoding o-fucosyltransferase family protein) (Fig. 6D and Supplementary Dataset S5: Sheet 25), which plays a key role in pollen tube growth [73]. This module was mainly enriched in GO-BP terms associated with the “monosaccharide biosynthetic process” and “lactone biosynthetic process”. Hub genes in module 4 included *Nbe.v1.1.chr11g18850* (a homolog gene encoding cell cycle checkpoint control protein family, *RAD9*), *Nbe.v1.1.chr18g38880* (a homolog of PASTICCINO 1 gene, *PAS1*), *Nbe.v1.1.chr07g33030* (a homolog of suppressor of k+ transport growth defect1 gene, *SKD1*), *Nbe.v1.1.chr11g11280* (a homolog gene encoding pre-rRNA-processing protein, *TSR2*), *Nbe.v1.1.chr03g08880* (a homolog of malectin domain kinesin 1 gene, *MDKIN1*), *Nbe.v1.1.chr03g12080* (a homolog of SC35-like splicing factor 33 gene, *SC35*) and *Nbe.v1.1.chr16g08230* (a homolog of *ORGANELLE TRANSCRIPT PROCESSING 439*, *OTP439*). The seed node of module 5 was *Nbe.v1.1.chr18g08160* (a homolog gene encoding S-locus lectin protein kinase family protein) (Fig. 6E and Supplementary Dataset S5: Sheet 25). The protein containing a G-type lectin domain is involved in diverse biological processes, including plant defense mechanisms and stress responses [74,75]. Module 5 was mainly enriched in GO-BP terms associated with the “regulation of intracellular steroid hormone receptor signaling pathway” and “regulation of gene expression”. Hub genes in module 5 included *Nbe.v1.1.chr19g16560* (a homolog of glutathione s-transferase TAU 8 gene, *GSTU8*), *Nbe.v1.1.chr02g05740* (a homolog of *WRKY6*), *Nbe.v1.1.chr15g15530* (a homolog of BRI1 like gene, *BRL1*), *Nbe.v1.1.chr06g24720* (a homolog of *RAF36*), *Nbe.v1.1.chr10g24840* (a homolog gene encoding drug/metabolite transporter superfamily protein) and *Nbe.v1.1.chr16g33430* (a homolog of callus expression of RBCS 101 gene, *CES101*) (Fig. 6E and Supplementary Dataset S5: Sheet 25).

Next, the GLay community clustering algorithm was applied. In the gene network for genes in expression pattern A, the GLay algorithm produced two large and two small modules (Fig. 7). Modules 1 (555 nodes and 8,538 edges) and 2 (508 nodes and 7,505 edges) were the largest modules, exhibiting high average connections per gene: 30.75 and 29.55 connections per gene, respectively. Modules 3 (38 nodes and 159 edges) and 4 (35 nodes and 180 edges) were the smallest modules with a more dispersed network topology and lower average connections per gene of 8.37 and 10.29, respectively. Modules 1 and 2 showed similar network characteristics, with clustering coefficients of approximately 0.30 and path lengths around 2.40. This showed that these modules have moderate connectivity but longer communication pathways. Meanwhile, modules 3 and 4, with similar network characteristics including strong local connectivity but shorter communication pathways, had clustering coefficients of approximately 0.50 and path lengths around 1.90. The modules contain several shared connecting nodes with other modules, indicating a potential relationship between their respective functions. The gene composition of the modules in expression pattern A includes a diverse array of enzymes, transcription factors, structural, signaling, stress response, and transport proteins, suggesting their involvement in signaling, transcriptional regulation, protein modification, and transport. The most frequently occurring *N. benthamiana* homolog genes in modules 1–4 were protein kinase superfamily protein, TPR-like superfamily protein, pleckstrin homology (PH) domain superfamily protein, and alpha/beta-hydrolases superfamily protein, respectively. Considering the 4 modules identified in the gene network of expression pattern A, Module 1 showed enrichment for several vascular tissue modification and defense response processes with evidence of involvement in plant grafting, including “regulation of secondary cell wall biogenesis” [17,76], “cellular response to molecule of bacterial origin” [77] and “xylem vessel member cell differentiation” [78] (Fig. 7 and Supplementary Dataset S6: Sheet 31). The top hub genes identified were *Nbe.v1.1.chr01g00700* (a homolog gene encoding NB-ARC domain-containing disease resistance protein), *Nbe.v1.1.chr02g05740* (a homolog of *WRKY6*), *Nbe.v1.1.chr06g24720* (a homolog of *RAF36*), and *Nbe.v1.1.chr19g13860* (a homolog gene encoding P-loop containing nucleoside triphosphate hydrolases superfamily protein) (Supplementary Dataset S7: Sheet 37). Biological processes such as the “nucleic acid metabolic process”, “DNA damage response”, and “nucleobase-containing compound metabolic process” which have been evident in plant grafting are captured in module 2 [79], and various catabolic processes including “alpha-amino acid”, “amino acid”, and “arginine catabolic process” in module 3 (Fig. 7 and Supplementary Dataset S6: Sheet 31). The top hub genes identified in module 2 were *Nbe.v1.1.chr02g29950* (a homolog of peroxin 6 gene, *PEX6*), *Nbe.v1.1.chr15g20950* (a homolog gene encoding prolyl oligopeptidase family protein), *Nbe.v1.1.chr02g03460* (a homolog of *OSCA2.3*) and *Nbe.v1.1.chr15g14280* (a homolog gene encoding zinc knuckle (CCHC-type) family protein) while *Nbe.v1.1.chr16g25250* (a homolog of ariadne 8 gene, *ARI8*), *Nbe.v1.1.chr15g09310* (a homolog of G protein-coupled receptor gene, *GPCR*) and *Nbe.v1.1.chr19g07140* (a homolog gene encoding RPAP2 IYO mate protein, *RIMA*) were in module 3 (Supplementary Dataset S7: Sheet 37). Module 4 revealed GO-BP terms “chromosome organization”, “nucleic acid metabolic process”, and “macromolecule methylation” (Fig. 7 and Supplementary Dataset S6: Sheet 31) with hub genes *Nbe.v1.1.chr03g08880* (a homolog of *MDKIN1*) and *Nbe.v1.1.chr08g22670* (a homolog gene encoding origin recognition complex protein 6, *ORC6*) (Supplementary Dataset S7: Sheet 37).

**Figure 7.**
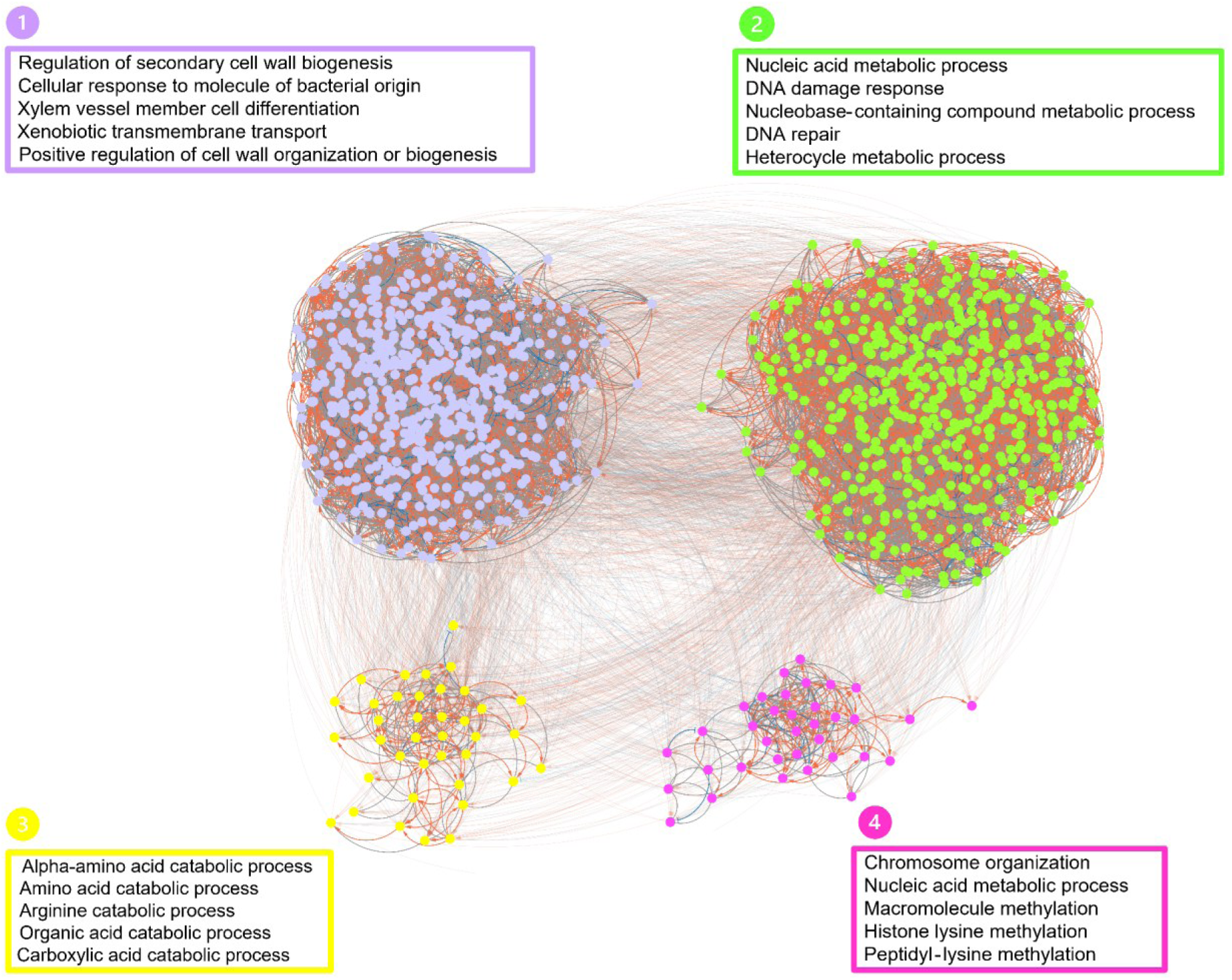
GLay module analysis of the Bayesian network for expression pattern A. The expression pattern A gene network was clustered into a total of four modules by the GLay community clustering algorithm in Cytoscape. GO enrichment analysis revealed that the modules identified had functionally distinct biological process GOs, including “regulation of secondary cell wall biogenesis”, “nucleic acid metabolic process”, “alpha-amino acid catabolic process”, and “chromosome organization”. Subnetworks are indicated by different colors and numbered sequentially as detected by the algorithm. Red arrows indicate gene up-regulation. Blue T-lines indicate gene down-regulation. Gray lines indicate uncertain gene regulation. See Supplementary Dataset S6: Sheet 31 for details.

#### Pattern B

Using the MCODE algorithm, a total of 9 modules (202 nodes and 578 edges, leaving 26 nodes unclustered) were identified in pattern B, however, 4 of the modules were significant with a threshold score > 5 and included 100 nodes and 393 edges (Fig. 8 and Supplementary Dataset S5: Sheet 2). For pattern B, the different *N. benthamiana* a homolog genes classes that formed the modules largely included transcription factors (such as ZIM-like 1, basic helix loop helix DNA-binding superfamily protein, AT-hook motif nuclear-localized protein 1, MYB family transcription factor, and homeodomain-like superfamily protein); enzymes (such as FKBP-like peptidyl-prolyl cis-trans isomerase family protein, glycosyl hydrolase superfamily protein, calcineurin-like metallo-phosphoesterase superfamily protein, fucosyltransferase 12, autoinhibited Ca^2+^/ATPase II, S-adenosyl-L-methionine-dependent methyltransferase superfamily protein and NAD(P)-linked oxidoreductase superfamily protein); signaling proteins (such as IQ-domain 8, rho GTPase-activating protein, cyclin-dependent kinase inhibitor SMR3-like protein, cyclic nucleotide-gated channel 14 and GCK domain-containing protein); stress response proteins (such as thaumatin-like protein 3, NB-ARC domain-containing disease resistance protein and disease resistance protein (CC-NBS-LRR class) family); cell cycle and division proteins (such as cell division cycle 45; mitotic-like cyclin 3B from *A. thaliana*, and BUB1-related (BUB1: budding uninhibited by benzymidazol 1) and transport proteins (such as organic solute transporter ostalpha protein (DUF300), nucleotide/sugar transporter family protein and endomembrane protein 70 protein family). A total of 4 significant MCODE modules were identified in the gene network of expression pattern B, which has genes predominantly highly expressed in the second state (Ave 1–7 DAG) only. The seed node of module 1 was *Nbe.v1.1.chr06g07540* (a homolog of cell division cycle 45 gene, *CDC45*) (Fig. 7A and Supplementary Dataset S5: Sheet 26), which plays a major role in regulating DNA replication initiation during the cell cycle process, including mitosis and meiosis [80]. This module was mainly enriched in GO-BP terms, “mitotic cell cycle checkpoint signaling”, “negative regulation of mitotic cell cycle”, and “cell cycle checkpoint signaling”. Hub genes in module 1 included *Nbe.v1.1.chr11g05390* (a homolog gene encoding brct5 domain containing protein 1, *BCP1*), *Nbe.v1.1.chr01g08340* (a homolog of phragmoplast-associated kinesin-related protein 2 gene, *PAKRP2*), *Nbe.v1.1.chr16g44610* (a homolog of *CYC3B*), *Nbe.v1.1.chr15g18390* (a homolog gene encoding Transducin/WD40 repeat-like superfamily protein) and *Nbe.v1.1.chr04g15130* (a homolog of budding uninhibited by benzymidazol 1 gene, *BUBR1*). The seed node of module 2 was *Nbe.v1.1.chr06g08060* (a homolog gene encoding AT-hook motif nuclear-localized protein 1, *AHL1*) (Fig. 8B and Supplementary Dataset S5: Sheet 26), which likely plays a role in chromatin organization and transcriptional regulation [81,82]. This module was mainly enriched in GO-BP terms, “establishment of spindle localization” and “protein signal transduction”. Hub genes in module 2 included *Nbe.v1.1.chr09g16700* (a homolog of *FASCIATED STEM 4*, *FAS4*), *Nbe.v1.1.chr02g29420* and *Nbe.v1.1.chr15g19470* (homolog genes encoding NB-ARC domain-containing disease resistance protein). The seed node of module 3 was *Nbe.v1.1.chr12g33170* (a homolog gene encoding CASP-like protein 5C2, *CASPL5C2*), a member of the CASP-like protein family, which may play a role in Casparian strip formation in plant roots that regulates the transport of water and solutes [83–85]. This module was mainly enriched in GO-BP terms, “response to maltose” and “fucose catabolic process”. Hub genes in module 3 included *Nbe.v1.1.chr12g33170* (a homolog of *CASPL5C2*), *Nbe.v1.1.chr11g20870* (a homolog gene encoding S-adenosyl-L-methionine-dependent methyltransferase superfamily protein) and *Nbe.v1.1.chr16g45560* (a homolog gene encoding transmembrane proteins) (Fig. 8C and Supplementary Dataset S5: Sheet 26). The seed node of module 4 was *Nbe.v1.1.chr01g39610* (a homolog gene encoding o-fucosyltransferase family protein), which is involved in several plant processes, including signaling pathways regulation, cell adhesion, and cell wall biosynthesis [86,87]. This module was mainly enriched in GO-BP terms, “dGMP metabolic process”, “glycoprotein transport”, and “GDP biosynthetic process”. Hub genes in module 4 included *Nbe.v1.1.chr01g39610* (a homolog gene encoding o-fucosyltransferase family protein), *Nbe.v1.1.chr17g07990* (a homolog gene encoding the PADRE gene family protein) and *Nbe.v1.1.chr06g21470* (a homolog of polyamine oxidase 4 gene, *PAO4*) (Fig. 8D and Supplementary Dataset S5: Sheet 26).

**Figure 8.**
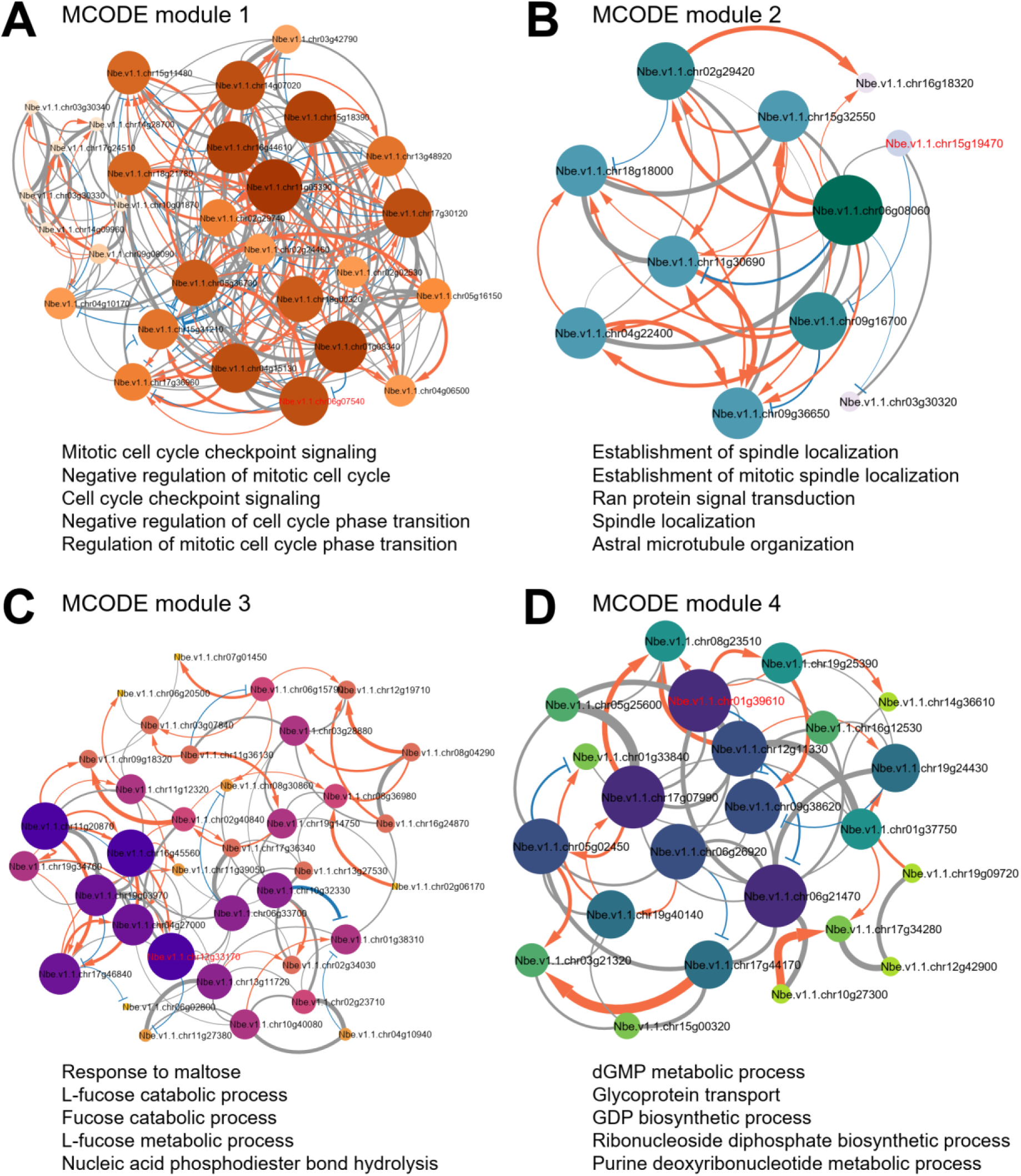
MCODE module analysis of the Bayesian network for expression pattern B. The module analysis using MCODE resulted in 4 significant modules, which included 100 nodes and 393 edges, and the enrichment of several GO-BP terms. Modules 1– 4 are the top four modules identified. **(A)** Module 1 (CDC45 complex) GO annotation has shown “mitotic cell cycle checkpoint signaling” and “negative regulation of mitotic cell cycle”. **(B)** Module 2 (AHL1 complex) GO annotation has revealed “establishment of spindle localization” and “establishment of mitotic spindle localization”. **(C)** Module 3 (CASPL5C2 complex) GO annotation has shown “response to maltose” and “L-fucose catabolic process”. **(D)** Module 4 (O-fucosyltransferase complex) GO annotation has shown “dGMP metabolic process”, “glycoprotein transport”, and “GDP biosynthetic process”. The seed node responsible for forming each module is indicated by red font. The module networks were visualized by Cytoscape (v3.10.2) by mapping the “degree parameter” to node size and color such that higher-degree nodes are darker in color and bigger in size. Red arrows indicate gene up-regulation. Blue T-lines indicate gene down-regulation. Gray lines indicate uncertain gene regulation. Line thickness indicates significance in gene regulation. All nodes are significantly enriched at p-value < 0.01. See Supplementary Dataset S5: Sheet 26 for details.

The GLay community clustering algorithm divided the network for genes in expression pattern B into 3 large modules (Fig. 9). Module 1 (79 nodes and 773 edges), Module 2 (89 nodes and 878 edges), and Module 3 (56 nodes and 545 edges) showed similar network characteristics, with clustering coefficients of approximately 0.48, high average connections per gene of 19.46–19.47, and path lengths of around 1.79. They had longer communication pathways with an average characteristic path length of 1.80. The small module 4 (4 nodes and four edges) showed a high clustering coefficient (0.58) and compactness (diameter 2), characteristics of a highly specialized functional unit. These GLay modules in expression pattern B have shared connecting nodes with other modules, indicating a potential relationship between their respective functions, and they consist mainly of enzymes, including transferases, hydrolases, and oxidoreductases. *N. benthamiana* homologs of PPR superfamily protein, NB-ARC domain-containing disease resistance protein, and P-loop containing nucleoside triphosphate hydrolases superfamily protein were the most frequently occurring genes in the modules and are involved in a complex network of cellular processes, including “organelle RNA regulation”, “plant immunity”, and “energy-dependent cellular activities”. A total of 4 GLay modules were also identified in the gene network of expression pattern B; however, module 4 had few nodes with no enriched GO-BP terms (Fig. 9). Biological process enrichment of “regulation of cell wall” and “secondary cell wall biogenesis” in module 1 links to its necessity for successful graft union formation and vascular reconnection in plant grafting [17] (Fig. 9 and Supplementary Dataset S6: Sheet 32) with main hub genes *Nbe.v1.1.chr11g36130* (a homolog gene encoding RNI-like superfamily protein), *Nbe.v1.1.chr11g32380* (a homolog of embryo defective 2739 gene, *EMB2739*), *Nbe.v1.1.chr08g18160* (a homolog gene encoding heat shock protein 20-like (HSP20-like) chaperones superfamily protein, *HSP17.6A*) and *Nbe.v1.1.chr10g01870* (a homolog gene encoding CCCH-type zinc finger family protein, *C3H17*) (Supplementary Dataset S7: Sheet 38). Biological processes such as “RNA metabolic process”, “nucleic acid metabolic process”, and “RNA modification” which have been evident in plant grafting are captured in module 2 [79], and several cell cycle processes including “cell cycle process”, “meiotic cell cycle process” and “mitotic cell cycle checkpoint signaling” in module 3 (Fig. 9 and Supplementary Dataset S6: Sheet 32). The top hub genes identified in module 2 were *Nbe.v1.1.chr06g26920* (a homolog gene encoding disease resistance (CC-NBS-LRR class) protein family) and *Nbe.v1.1.chr06g33700* (a homolog gene encoding NAD(P)-binding Rossmann-fold superfamily protein), whiles *Nbe.v1.1.chr18g00320* (a homolog gene encoding FKBP-like peptidyl-prolyl cis-trans isomerase family protein), *Nbe.v1.1.chr16g44610* (a homolog of *CYC3B*) and *Nbe.v1.1.chr15g18390* (a homolog gene encoding Transducin/WD40 repeat-like superfamily protein) were in module 3 (Supplementary Dataset S7: Sheet 38).

**Figure 9.**
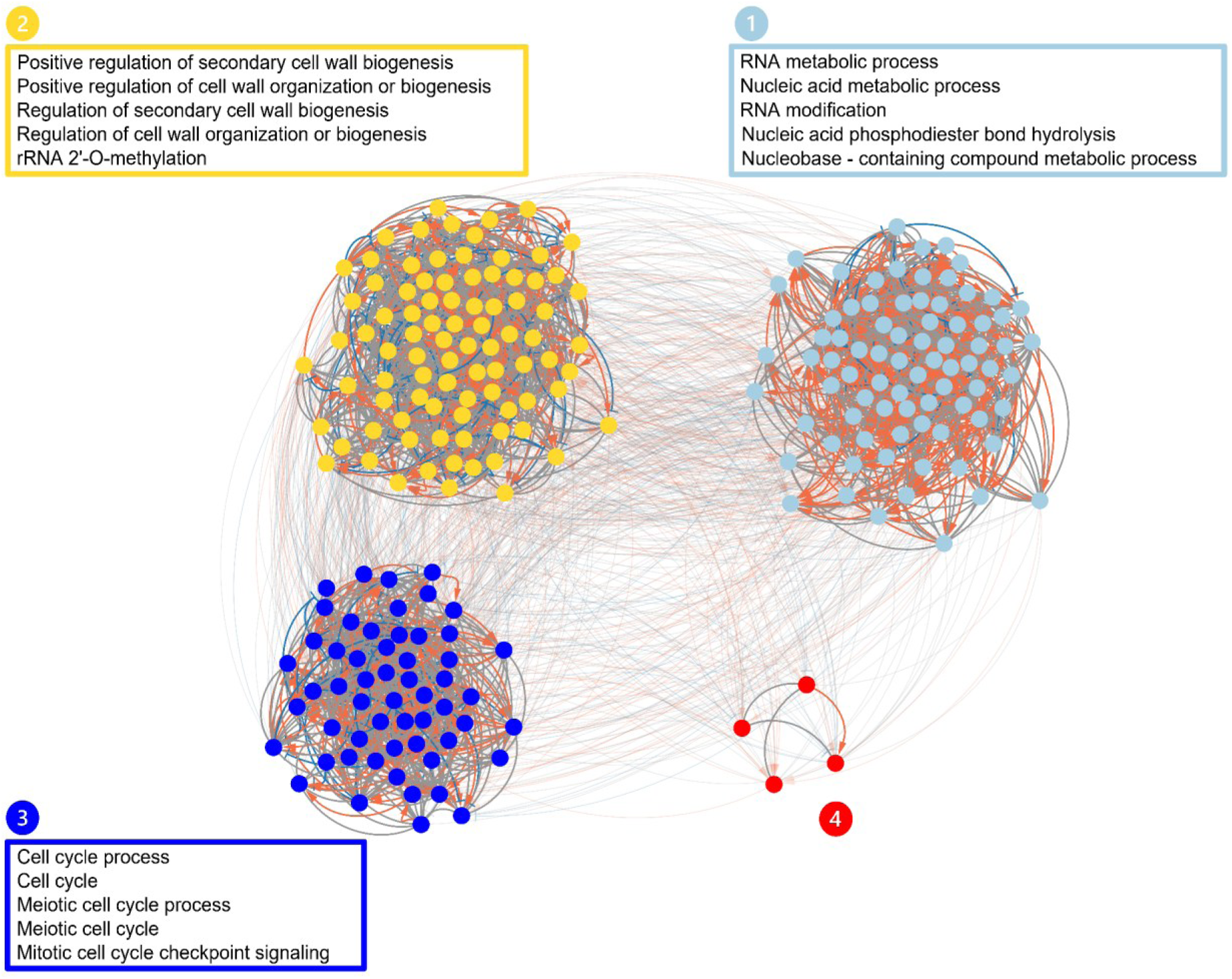
GLay module analysis of the Bayesian network for expression pattern B. The expression pattern B gene network was clustered into a total of four modules by the GLay community clustering algorithm in Cytoscape. GO enrichment analysis revealed that the modules identified had functionally distinct biological process GOs, including “RNA metabolic process”, “positive regulation of secondary cell wall biogenesis”, and “cell cycle process”. Subnetworks are indicated by different colors and numbered sequentially as detected by the algorithm. Red arrows indicate gene up-regulation. Blue T-lines indicate gene down-regulation. Gray lines indicate uncertain gene regulation. See Supplementary Dataset S6: Sheet 32 for details.

#### Pattern C

In pattern C, MCODE revealed a total of 18 modules (873 nodes and 1997 edges, leaving 260 nodes unclustered). There were 6 significant MCODE modules in pattern C, which included 375 nodes and 1,041 edges (Fig. 10 and Supplementary Dataset S5: Sheet 27). The types of *N. benthamiana* homologs that formed the modules in the pattern predominantly included some stress response proteins (such as NPR1-like protein 3, disease resistance protein (CC-NBS-LRR class) family, DNA heat shock family protein, disease resistance protein (TIR-NBS-LRR class) family and RPM1 interacting protein 13); cell cycle and division proteins (such as origin recognition complex protein 6, cell division control, cdc6 and mitotic-like cyclin 3B from *A. thaliana*); transport proteins (such as MATE efflux family protein, K^+^ uptake transporter 3 and lipid transporter); signaling proteins (such as protein kinase superfamily protein, cyclic nucleotide gated channel 8, phototropic-responsive NPH3 family protein, and BRI1 like); transcription factors (such as SEUSS-like 2, bzip transcription factor family protein, myelin transcription factor-like protein, DNA-binding transcription factor 2 and NAC-like, activated by AP3/PI) and enzymes (such as Lipase class 3 family protein, oxidoreductases, acting on NADH or NADPH and FAD-binding berberine family protein [86]. In the expression pattern C gene regulatory network, which has genes predominantly highly expressed in the third state, only 6 significant MCODE modules were identified from the analysis. The seed node of module 1 was *Nbe.v1.1.chr10g12880* (a homolog gene encoding bromodomain protein) (Fig. 10A and Supplementary Dataset S5: Sheet 27), which is involved in key processes in plants, including chromatin remodeling, transcriptional regulation, development, and stress responses [88,89]. This module is mainly enriched in GO-BP terms, “regulation of cell killing”, “host innate immune response by symbiont”, and “modulation by symbiont of host programmed cell death”. Hub genes in module 1 included *Nbe.v1.1.chr15g31440* (a homolog gene encoding transmembrane protein), *Nbe.v1.1.chr13g10520* (a homolog gene encoding the phosphatase 2C family protein), *Nbe.v1.1.chr15g47660* and *Nbe.v1.1.chr05g05430* (homolog genes encoding major facilitator superfamily proteins). The seed node of module 2 was *Nbe.v1.1.chr08g22670* (a homolog of *ORC6*) (Fig. 10B and Supplementary Dataset S5: Sheet 27), which is reported to be involved in plant processes including DNA replication initiation [90]. This module mainly enriched GO-BP terms, “synaptic vesicle priming”, “establishment of organelle localization”, and “chromosome organization”. Hub genes in module 2 included *Nbe.v1.1.chr15g07140* (a homolog gene encoding RING/FYVE/PHD zinc finger superfamily protein), *Nbe.v1.1.chr13g22810* (a homolog of *TRITHORAX-RELATED PROTEIN 6*, *ATXR6*) and *Nbe.v1.1.chr08g22670* (a homolog of *ORC6*). The seed node of module 3 was *Nbe.v1.1.chr02g09880* (a homolog gene encoding PPR repeat-containing protein) (Fig. 10C and Supplementary Dataset S5: Sheet 27), involved in photosynthesis, respiration, cytoplasmic male sterility, and seed development through the regulation of organelle gene expression [91–94]. This module was mainly enriched in GO-BP terms, “nucleic acid phosphodiester bond hydrolysis”, and “organic cyclic compound metabolic process”. Hub genes in module 3 included *Nbe.v1.1.chr05g33620* (a homolog gene encoding PPR superfamily protein), *Nbe.v1.1.chr02g29950* (a homolog of *PEX6*), *Nbe.v1.1.chr17g09430* (a homolog of the DNA polymerase kappa gene, *POLK*), *Nbe.v1.1.chr19g27060* (a homolog gene encoding the DNA/RNA polymerases superfamily protein), *Nbe.v1.1.chr18g27360* (a homolog gene encoding the ankyrin repeat family protein) and *Nbe.v1.1.chr02g08930* (a homolog gene encoding LEUNIG-like protein, *LUH*). The seed node of module 4 was *Nbe.v1.1.chr05g33620* (a homolog gene encoding helicase/SANT-associated protein) (Fig. 10D and Supplementary Dataset S5: Sheet 27), which is involved in key processes in plants, including chromatin remodeling, transcriptional regulation, and various aspects of RNA metabolism [95,96]. This module was mainly enriched in GO-BP terms, “amyloplast organization”, “detection of gravity”, and “histone H2A acetylation”. Hub gene in module 4 included *Nbe.v1.1.chr12g26540* (a homolog of NDR1/HIN1-like 8 gene, *NHL8*). The seed node of module 5 was *Nbe.v1.1.chr02g36940* (a homolog gene encoding PPR repeat superfamily protein) (Fig. 10E and Supplementary Dataset S5: Sheet 27), which is involved in photosynthesis, respiration, cytoplasmic male sterility, and seed development through the regulation of organelle gene expression [91–94]. This module was mainly enriched in GO-BP terms, “regulation of termination of DNA-templated transcription”, and “positive regulation of telomere maintenance via telomerase”. Hub genes in module 5 included *Nbe.v1.1.chr08g03260* (a homolog of histone-lysine N-methyltransferase gene, *ASHH3*), *Nbe.v1.1.chr10g38460* (a homolog gene encoding TPX2-like protein 3, *TPXL3*), *Nbe.v1.1.chr16g06830* (a homolog gene encoding CW14 protein (DUF1336), *CW14*), *Nbe.v1.1.chr15g19050* (a homolog gene encoding DNA repair (Rad51) family protein, *DMC1*), *Nbe.v1.1.chr07g29080* (a homolog of RPA70-kDa subunit B gene, *RPA70B*), *Nbe.v1.1.chr07g03980* (a homolog of *bHLH121*) and *Nbe.v1.1.chr05g16810* (a homolog of maternal effect embryo arrest 47 gene, *MEE47*). The seed node of module 6 was *Nbe.v1.1.chr13g00825* (a homolog of FORKED-LIKE8 gene, *FL8*), involved in auxin transport, leaf morphogenesis, and vascular patterning [97,98]. This module was mainly enriched in “regulation of actin filament bundle assembly”, “intron homing”, and “protein serine/threonine kinase activity”. Hub genes in module 6 included *Nbe.v1.1.chr13g06550* (a homolog gene encoding cysteinyl-tRNA synthetase class Ia family protein, *SYCO*), *Nbe.v1.1.chr01g33940* (a homolog of *REDUCED CHLOROPLAST COVERAGE*, *REC1*), *Nbe.v1.1.chr13g00825* (a homolog of *FL8*) and *Nbe.v1.1.chr05g05310* (a homolog gene encoding zinc-finger protein 14, *ZAT14*) (Fig. 10F and Supplementary Dataset S5: Sheet 27).

**Figure 10.**
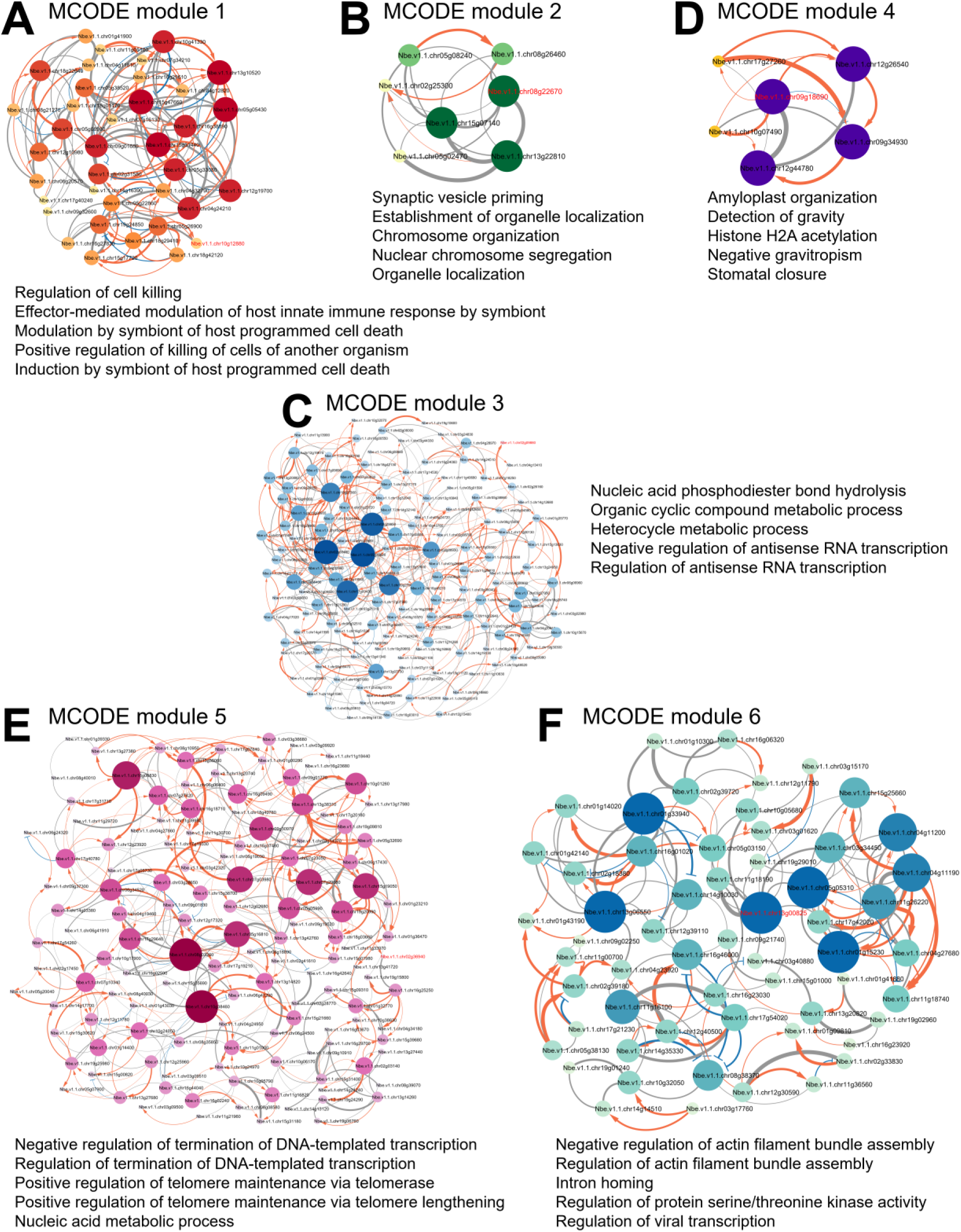
MCODE module analysis of the Bayesian network for expression pattern C. The module analysis using MCODE revealed 6 significant modules, which include 375 nodes and 1,041 edges, and the enrichment of several GO-BP terms. Modules 1 – 6 are the top six modules identified. **(A)** Module 1 (Bromodomain complex) GO annotation has shown regulation of “cell killing”, “effector-mediated modulation of host innate immune response by symbiont”, and “modulation by symbiont of host programmed cell death”. **(B)** Module 2 (Origin recognition complex) GO annotation has revealed “synaptic vesicle priming”, “establishment of organelle localization”, and “chromosome organization”. **(C)** Module 3 (PPR repeat-containing complex) GO annotation has shown “nucleic acid phosphodiester bond hydrolysis”, “organic cyclic compound metabolic process”, and “heterocycle metabolic process”. **(D)** Module 4 (Helicase/SANT-associated DNA binding complex) GO annotation has shown “amyloplast organization”, “detection of gravity”, and “histone H2A acetylation”. **(E)** Module 5 (PPR repeat superfamily complex) GO annotation has shown “negative regulation of termination of DNA-templated transcription” and “regulation of termination of DNA-templated transcription”. **(F)** Module 6 (Auxin canalization complex) GO annotation has shown regulation of “negative regulation of actin filament bundle assembly”, “regulation of actin filament bundle assembly”, “intron homing”, and “regulation of protein serine/threonine kinase activity”. The seed node responsible for forming each module is indicated by red font. The module networks were visualized by Cytoscape (v3.10.2) by mapping the “degree parameter” to node size and color such that higher-degree nodes are darker in color and bigger in size. Red arrows indicate gene up-regulation. Blue T-lines indicate gene down-regulation. Gray lines indicate uncertain gene regulation. Line thickness indicates significance in gene regulation. All nodes are significantly enriched at p-value < 0.01. See Supplementary Dataset S5: Sheet 27 for details.

In pattern C, GLay community clustering analysis revealed 3 large modules (Module 1: 169 nodes and 1,687 edges; Module 2: 434 nodes and 5,164 edges; Module 3: 520 nodes and 7,319 edges) and a small module (10 nodes and 17 edges) (Fig. 11). Module 1, though the third largest, had the highest clustering coefficient (0.44) and average density (0.12) among all modules, indicating dense local neighborhoods. Modules 2 and 3 showed similar network characteristics, with clustering coefficients of approximately 0.28, path lengths of around 2.50, high average connections per gene, 23.79 and 28.15 connections per gene, respectively. This showed that these modules have moderate connectivity but longer communication pathways. Module 4 had loosely connected nodes with a low average degree (3.4 connections per gene) and shorter communication pathways with path lengths of 1.93. Similarly, the GLay modules in expression pattern C comprise several shared connecting nodes and various enzymes, transcription factors, structural proteins, signaling proteins, stress response proteins, and transport proteins. Notably includes *N. benthamiana* homologs of kinases, NAC domain-containing proteins, transferases, disease resistance protein (CC-NBS-LRR class) family, and NB-ARC domain-containing disease resistance protein. Following GLay module analysis, 4 modules were identified in the gene regulatory network of expression pattern C (Fig. 11). In module 1 enriched GO terms in the biological process were related to the regulation of cellular processes including “entrainment of circadian clock”, “regulation of actin filament bundle assembly”, “negative regulation of actin filament bundle assembly” and “positive regulation of JNK cascade” (Fig. 11 and Supplementary Dataset S6: Sheet 33) with main hub genes, *Nbe.v1.1.chr09g01630* (a homolog of detoxification 29 gene, *DTX29*), *Nbe.v1.1.chr03g28630* (a homolog gene encoding inositol 1,3,4-trisphosphate 5/6-kinase family protein, *ITPK1*) and *Nbe.v1.1.chr18g29410* (a homolog of phosphate transporter PHO1 homolog 3 gene, *PHO1;H3*) (Supplementary Dataset S7: Sheet 39). These significant processes may be tightly linked to the regulation of the cellular process during grafting [99–101]. In module 2, enriched GO terms in the biological process category identified related to “nucleic acid metabolism” and “genetic exchange” which have been observed to contribute to graft success [102] including “nucleic acid metabolic process”, “heterocycle metabolic process”, “DNA damage response” and “RNA metabolic process” (Fig. 11 and Supplementary Dataset S6: Sheet 33) with main hub genes, *Nbe.v1.1.chr14g20700* (a homolog gene encoding the 2-oxoglutarate (2OG) and Fe(II)-dependent oxygenase superfamily protein), *Nbe.v1.1.chr02g29950* (a homolog of *PEX6*), *Nbe.v1.1.chr18g27010* (a homolog gene encoding the F-box/RNI-like superfamily protein) and *Nbe.v1.1.chr19g02960* (a homolog gene encoding phloem protein 2-B1, *PP2-B1*) (Supplementary Dataset S7: Sheet 39). Similarly, in module 3, enriched GO terms such as “nucleic acid metabolic process”, “nucleobase-containing compound metabolic process”, and “heterocycle metabolic process” were identified with hub genes *Nbe.v1.1.chr15g14280* (a homolog gene encoding the Zinc knuckle (CCHC-type) family protein), *Nbe.v1.1.chr15g20950* (a homolog gene encoding the prolyl oligopeptidase family protein), *Nbe.v1.1.chr06g34520* (a homolog of *TRZ4*) and *Nbe.v1.1.chr17g40780* (a homolog gene encoding TPR-like superfamily protein). In module 4, several steroid metabolism-related GO terms with intricate evidence in plant grafting [103], including “steroid biosynthetic process”, “sterol biosynthetic process”, “protein peptidyl-prolyl isomerization”, and “peptidyl-proline modification” were enriched (Fig. 11 and Supplementary Dataset S6: Sheet 33) with *Nbe.v1.s00050g15120* (a homolog of small defense-associated protein 1 gene, *SDA1*) identified as the main hub gene (Supplementary Dataset S7: Sheet 39).

**Figure 11.**
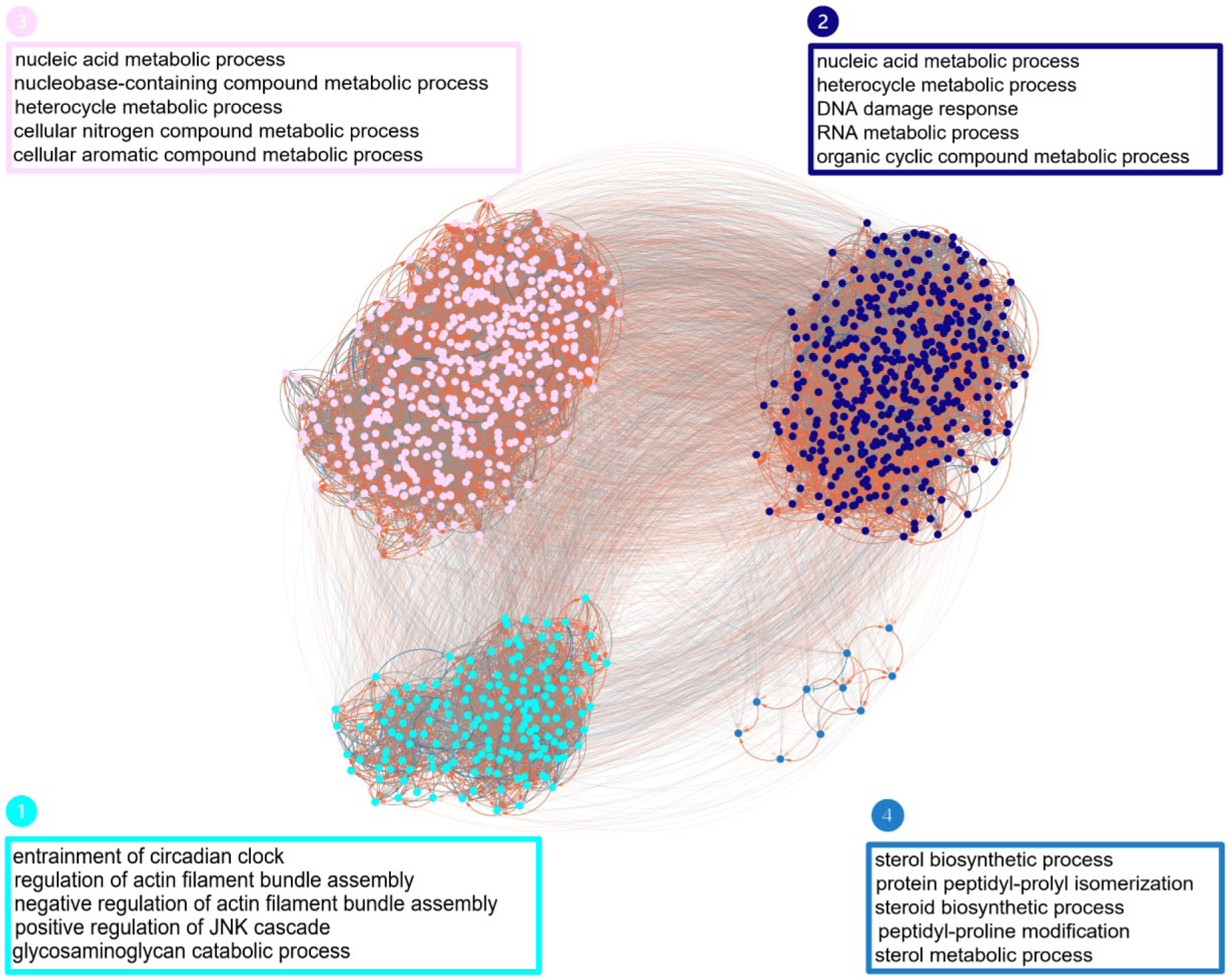
GLay module analysis of the Bayesian network for expression pattern C. The expression pattern C gene network was clustered into a total of four modules by the GLay community clustering algorithm in Cytoscape. GO enrichment analysis revealed that the modules identified had functionally distinct biological process GOs, including “entrainment of circadian clock”, “nucleic acid metabolic process”, and “sterol biosynthetic process”. Subnetworks are indicated by different colors and numbered sequentially as detected by the algorithm. Red arrows indicate gene up-regulation. Blue T-lines indicate gene down-regulation. Gray lines indicate uncertain gene regulation. See Supplementary Dataset S6: Sheet 33 for details.

#### Pattern D

In pattern D, the MCODE algorithm revealed a total of 7 modules (153 nodes and 463 edges, leaving 23 nodes unclustered). The significant MCODE modules with a threshold score of > 5 were 4 and comprised 98 nodes and 352 edges (Fig. 12 and Supplementary Dataset S5: Sheet 28). For pattern D, the different *N. benthamiana* homolog gene families that formed the modules mostly included transcription factors (such as E2F transcription factor 3, transcription factor IIIB and NAC domain-containing protein 100); enzymes (such as cellulose-synthase-like C6, glucan synthase-like 10 and NAD(H) kinase 3); signaling proteins (such as protein kinase superfamily protein, leucine-rich repeat (LRR) family protein and K^+^ uptake transporter 3); transport proteins (such as amino acid permeases, ABC transporters) and DNA/RNA-Related Proteins (such as SET domain group 4, nuclear RNA polymerase C1 and plastid transcriptionally active 3). In the expression pattern D gene regulatory network, which has genes predominantly lowly expressed in the third state only, a total of 4 significant modules were identified in the MCODE analysis. The seed node of module 1 was *Nbe.v1.1.chr06g12060* (a homolog gene encoding ARM repeat superfamily protein, *CAP-G2*) (Fig. 12A and Supplementary Dataset S5: Sheet 28), which plays a major role in regulating cell division through the spatial arrangement of centromeres mediated by ARM repeats [104,105]. This module is mainly enriched in GO-BP terms, “positive regulation of the meiotic cell cycle” and “DNA replication initiation”. Hub genes in module 1 included *Nbe.v1.1.chr18g11830* (a homolog of cyclin A3;4 gene, *CYCA3;4*), *Nbe.v1.1.chr15g00270* (a homolog gene encoding growing plus-end tracking protein 2, *GPT2*), *Nbe.v1.1.chr19g05150* (a homolog of condensin gene, *CAP-D3*) and *Nbe.v1.1.chr05g21840* (a homolog of TORTIFOLIA1-like protein 2 gene, *TOR1L2*). The seed node of module 2 was *Nbe.v1.1.chr15g09830* (a homolog gene encoding PPR repeat superfamily protein) (Fig. 12B and Supplementary Dataset S5: Sheet 28) is involved in photosynthesis, respiration, cytoplasmic male sterility, and seed development through the regulation of organelle gene expression [91–94]. This module was mainly enriched in GO-BP terms, “RNA modification” and “nucleic acid phosphodiester bond hydrolysis”. Hub genes in module 2 included *Nbe.v1.1.chr19g27950* and *Nbe.v1.1.chr15g09830* (homolog genes encoding TPR-like superfamily proteins), *Nbe.v1.1.chr04g21100* (a homolog of pigment defective 338 gene, *PDE338*) and *Nbe.v1.1.chr03g20220* (a homolog of embryo defective 3103 gene, *EMB3103*). The seed node of module 3 was *Nbe.v1.1.chr19g08450* (a homolog gene encoding the galactose oxidase/kelch repeat superfamily protein) (Fig. 12C and Supplementary Dataset S5: Sheet 28), which is involved in plant iron deficiency response [106]. This module was mainly enriched in GO-BP terms, “regulation of response to extracellular stimulus” and “regulation of response to nutrient levels”. The hub gene identified in module 3 was *Nbe.v1.1.chr03g42690* (a homolog of *AGDP1*). The seed node of module 4 was *Nbe.v1.1.chr03g04570* (a homolog gene encoding the serine/threonine kinase), which regulates signaling pathways, stress acclimation, and development [107,108]. This module was mainly enriched in the GO-BP terms “RNA modification”, “NADP biosynthetic process”, and “peptidyl-lysine demethylation”. Hub genes in module 4 included *Nbe.v1.1.chr02g03250* (a homolog gene encoding ran BP2/NZF zinc finger-like superfamily protein) and *Nbe.v1.1.chr07g11510* (a homolog of *NAC100*) (Fig. 12D and Supplementary Dataset S5: Sheet 28).

**Figure 12.**
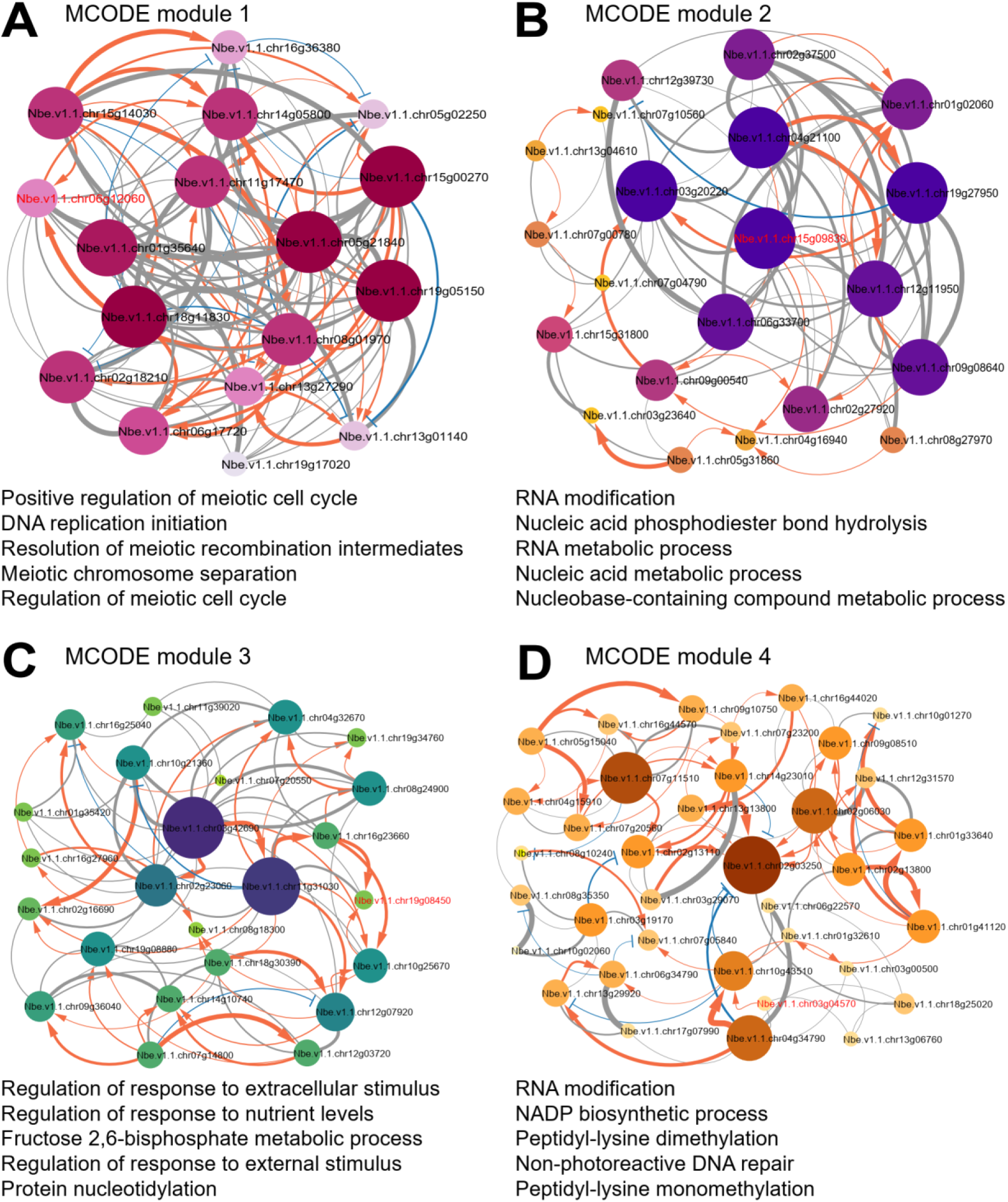
MCODE module analysis of the Bayesian network for expression pattern D. The module analysis using MCODE resulted in 4 significant modules, which included 98 nodes and 352 edges, and the enrichment of several GO-BP terms. Modules 1–4 are the top four modules identified. **(A)** Module 1 (ARM repeat complex) GO annotation has shown “positive regulation of meiotic cell cycle”, “DNA replication initiation”, and “resolution of meiotic recombination intermediates”. **(B)** Module 2 (PPR repeat complex) GO annotation has revealed “RNA modification”, “RNA metabolic process”, and “nucleic acid phosphodiester bond hydrolysis”. **(C)** Module 3 (Galactose oxidase/kelch complex) GO annotation has shown “regulation of response to extracellular stimulus”, “regulation of response to nutrient levels”, and “fructose 2,6-bisphosphate metabolic process”. **(D)** Module 4 (Serine/Threonine-kinase complex) GO annotation has shown “RNA modification”, “NADP biosynthetic process”, and “peptidyl-lysine dimethylation”. The seed node responsible for forming each module is indicated by red font. The module networks were visualized by Cytoscape (v3.10.2) by mapping the “degree parameter” to node size and color such that higher-degree nodes are darker in color and bigger in size. Red arrows indicate gene up-regulation. Blue T-lines indicate gene down-regulation. Gray lines indicate uncertain gene regulation. Line thickness indicates significance in gene regulation. All nodes are significantly enriched at p-value < 0.01. See Supplementary Dataset S5: Sheet 28 for details.

In the gene network for genes in expression patterns D, the GLay community clustering algorithm produced 3 modules (Fig. 13). Module 1 (23 nodes and 132 edges) was the smallest with a high average degree and clustering coefficient of 0.52 and 0.66, indicating a high degree of local connectivity but a low average connection per gene and path lengths of 11.48 and 1.48, respectively. The **c**omposition of genes in module 1 is a mix of structural proteins, transcription factors, transport, and stress-responsive proteins. Module 2 (74 nodes and 657 edges) and Module 3 (79 nodes and 751 edges) showed similar network characteristics, with clustering coefficients of approximately 0.40 and path lengths around 1.80. This showed that these modules have moderate connectivity but longer communication pathways. The GLay modules shared connecting nodes. The SKU5 similar protein, TPR-like, and ARM repeat superfamily protein were the most frequently occurring *N. benthamiana* homolog gene families in modules 2 and 3. GLay module analysis revealed 3 modules in the gene network of the expression pattern D (Fig. 13). In module 1, enriched GO terms in the biological process related to RNA metabolism were “RNA modification”, “nucleic acid phosphodiester bond hydrolysis”, “RNA metabolic process”, and “organic cyclic compound metabolic process” with *Nbe.v1.1.chr03g19170* (a homolog gene encoding xylulose kinase-1, *XK-1*) as the main hub gene screened (Fig. 13, Supplementary Dataset S6: Sheet 34 and Supplementary Dataset S7: Sheet 40). These RNA metabolism-related biological processes potentially influence RNA stability and signaling pathways [109]. Biological processes such as “regulation of cell morphogenesis”, “macromolecule modification”, “nucleic acid phosphodiester bond hydrolysis” and “arp2/3 complex-mediated actin nucleation” which have been intricated in plant grafting are captured in modules 2 [110–112], and several nucleic acid metabolisms GO terms including “RNA modification”, “nucleic acid phosphodiester bond hydrolysis”, “RNA metabolic process”, “macromolecule modification” and “nucleic acid metabolic process” in module 3 (Fig. 13, and Supplementary Dataset S6: Sheet 34). The top hub genes identified in module 2 were *Nbe.v1.1.chr15g00270* (a homolog of *GPT2*), *Nbe.v1.1.chr02g06170* (a homolog of syntaxin of plants 112 gene, *SYP112*) and *Nbe.v1.1.chr02g18210* (a homolog *ORC5*) while *Nbe.v1.1.chr11g31030* (a homolog of phosphate transporter 3;1 gene, *PHT3;1*), *Nbe.v1.1.chr03g42690* (a homolog of *AGDP1*), and *Nbe.v1.1.chr08g05060* (a homolog of PPR repeat superfamily protein) were in module 3 (Supplementary Dataset S7: Sheet 40).

**Figure 13.**
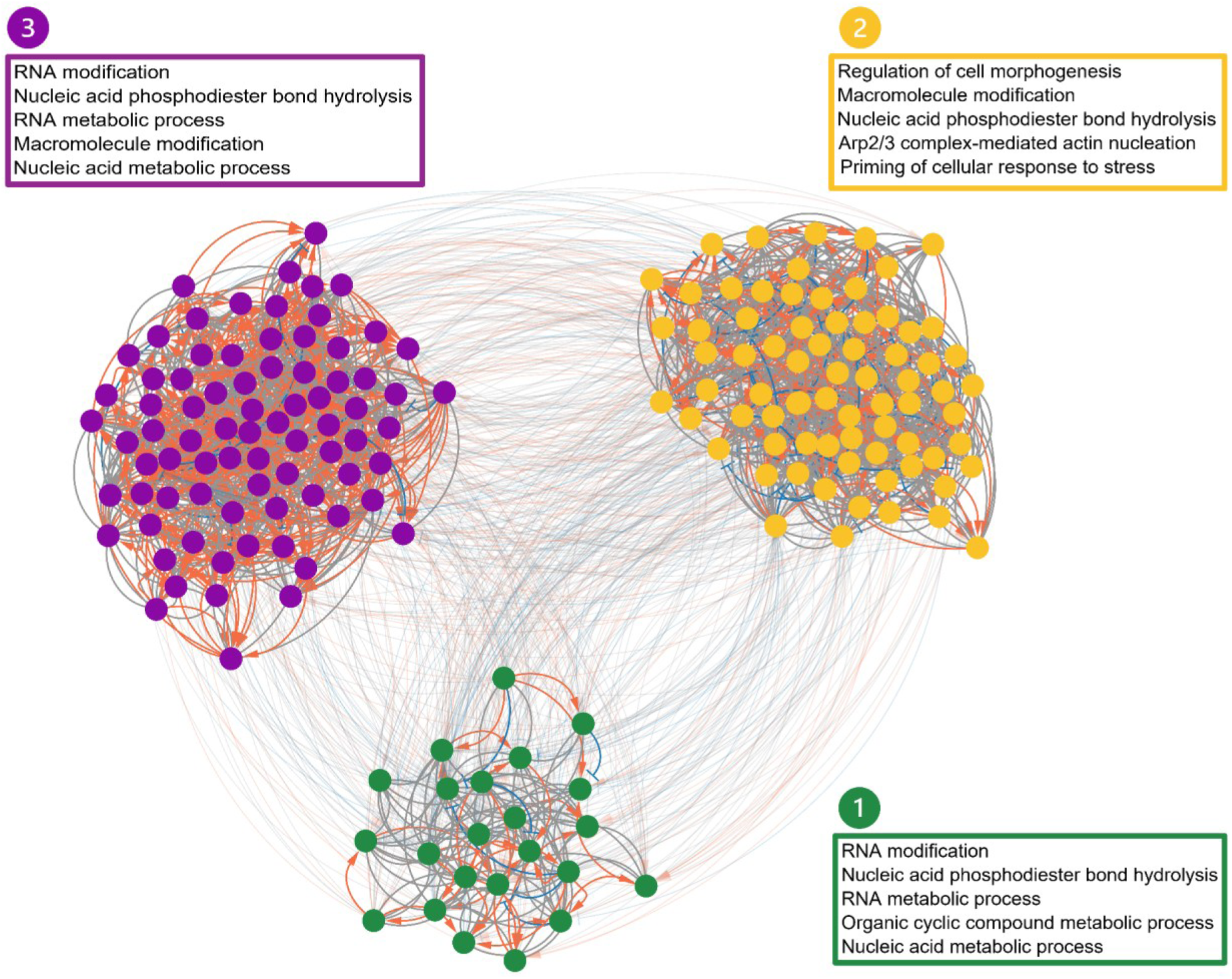
GLay module analysis of the Bayesian network for expression pattern D. The expression pattern D gene network was clustered into a total of three modules by the GLay community clustering algorithm in Cytoscape. GO enrichment analysis revealed that the modules identified had functionally distinct biological process GOs, including “RNA modification”, “regulation of cell morphogenesis”, and “RNA modification”. Subnetworks are indicated by different colors and numbered sequentially as detected by the algorithm. Red arrows indicate gene up-regulation. Blue T-lines indicate gene down-regulation. Gray lines indicate uncertain gene regulation. See Supplementary Dataset S6: Sheet 34 for details.

#### Pattern E

The total MCODE modules in pattern E were 3 (48 nodes and 210 edges, leaving 3 nodes unclustered), and all 3 were significant modules with a threshold score > 5 (Fig. 14 and Supplementary Dataset S5: Sheet 29). For pattern E, the different types of *N. benthamiana* homolog genes that formed the modules primarily included some transcription factors (such as cycling DOF factor 2, OBF-binding protein 3, plant regulator RWP-RK family protein and NAC domain-containing protein 83); enzymes (such as FAD/NAD(P)-binding oxidoreductase family protein, aldehyde oxidase 4 and UDP-glucosyl transferase 73B3); stress response proteins (such as senescence-associated gene 101); and DNA/RNA-related proteins (such as R3H domain protein, sirtuin 1 and chromatin remodeling 5protein 70 protein family). In the expression pattern E gene regulatory network, which has genes predominantly lowly expressed in the second state, 3 significant modules were identified in the MCODE analysis. The seed node of module 1 was *Nbe.v1.1.chr05g16670* (a homolog gene encoding leucine-rich repeat protein kinase family protein) (Fig. 14A and Supplementary Dataset S5: Sheet 29), whose precise role in plants is unknown, may have similar functions as the receptor kinase family in plants, including hormone signaling and defense mechanisms [113,114]. This module was mainly enriched in GO-BP terms, “chloroplast-nucleus signaling pathway”, “apocarotenoid biosynthetic process”, and “tertiary alcohol biosynthetic process”. The hub gene in module 1 was *Nbe.v1.1.chr05g01150* (a homolog of *PPR19*). The seed node of module 2 was *Nbe.v1.1.chr05g22030* (a homolog gene encoding o-glycosyl hydrolases family 17 protein, *GH17*) (Fig. 14B and Supplementary Dataset S5: Sheet 29) is involved in callose degradation, plasmodesmata regulation, and stress response [115,116]. This module was mainly enriched in GO-BP terms, “histone H3-K9 acetylation”, “histone H3-K9 modification”, and “regulation of seed growth”. The hub gene in module 2 was *Nbe.v1.1.chr02g21910* (a homolog of the senescence-associated gene 101 gene, *SAG101*). The seed node of module 3 was *Nbe.v1.1.chr02g08190* (a homolog gene encoding the alpha 1,4-glycosyltransferase family protein) is involved in plant cell wall polysaccharides biosynthesis, specifically a component of pectin known as galactan [117]. This module was mainly enriched in GO-BP terms, “positive regulation of wax” and “abscisic acid biosynthetic processes”. The hub gene identified was *Nbe.v1.1.chr15g07360* (a homolog of *TRICHOME BIREFRINGENCE-LIKE 41*, *TBL41*) (Fig. 14C and Supplementary Dataset S5: Sheet 29).

**Figure 14.**
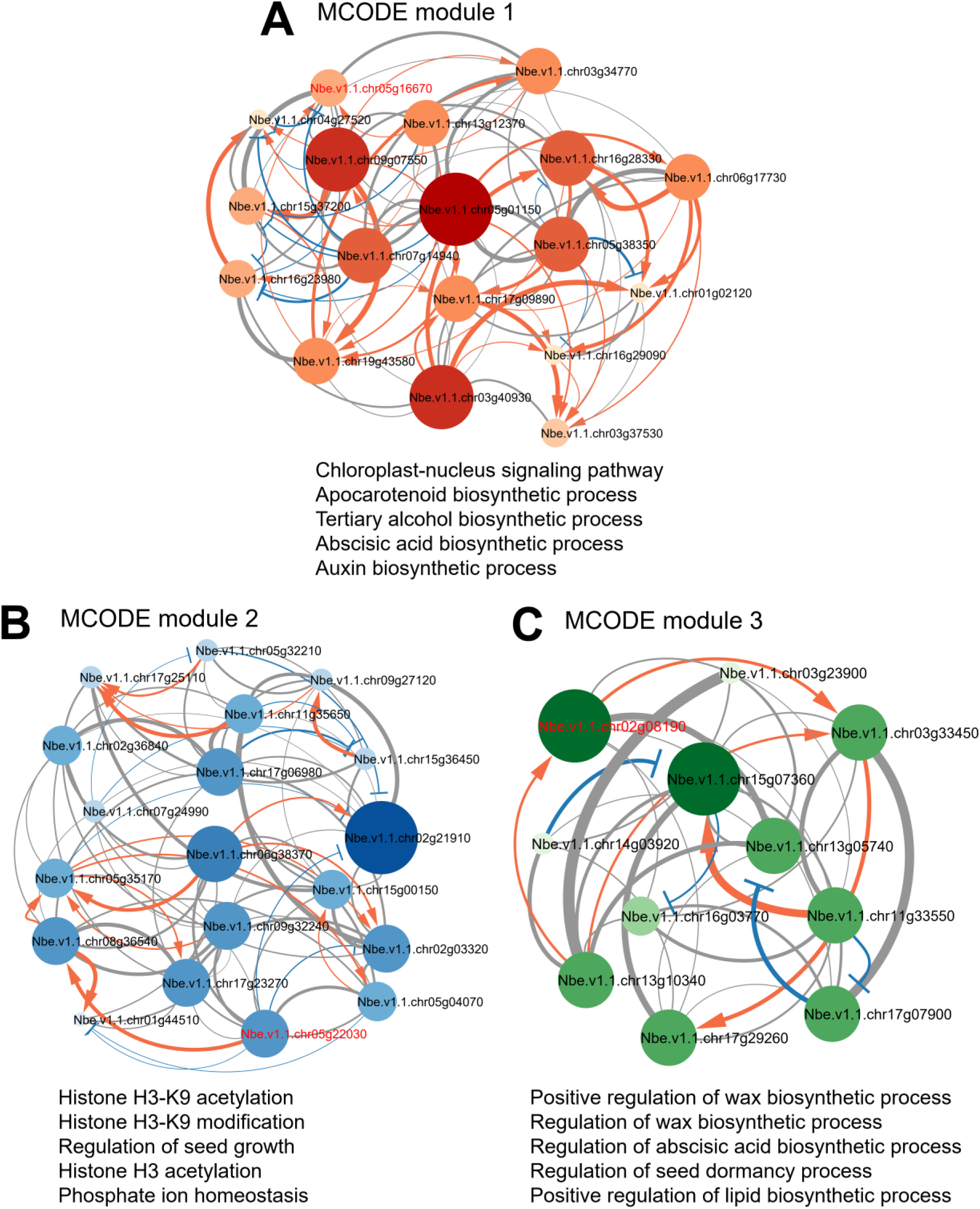
MCODE module analysis of the Bayesian network for expression pattern E. The module analysis using MCODE revealed 3 significant modules, which include 48 nodes and 210 edges, and the enrichment of several GO-BP terms. Modules 1 – 3 are the top three modules identified. **(A)** Module 1 (Leucine-rich repeat complex) GO annotation has shown “chloroplast-nucleus signaling pathway”, “apocarotenoid biosynthetic process”, and “tertiary alcohol biosynthetic process”. **(B)** Module 2 (O-Glycosyl hydrolases complex) GO annotation has revealed histone “H3-K9 acetylation and modification” and “regulation of seed growth”. **(C)** Module 3 (Alpha 1,4-glycosyltransferase complex) GO annotation has shown “positive regulation of wax biosynthetic process” and “regulation of abscisic acid biosynthetic process”. The seed node responsible for forming each module is indicated by red font. The module networks were visualized by Cytoscape (v3.10.2) by mapping the “degree parameter” to node size and color such that higher-degree nodes are darker in color and bigger in size. Red arrows indicate gene up-regulation. Blue T-lines indicate gene down-regulation. Gray lines indicate uncertain gene regulation. Line thickness indicates significance in gene regulation. All nodes are significantly enriched at p-value < 0.01. See Supplementary Dataset S5: Sheet 29 for details.

The GLay community clustering analysis revealed 3 modules in the gene regulatory network for expression pattern E (Fig. 15). Module 1 (20 nodes and 140 edges) and Module 2 (19 nodes and 116 edges) showed similar network characteristics, with clustering coefficients of approximately 0.80 and path lengths around 1.30. NAC domain-containing proteins and various enzymes, including oxidoreductases, hydrolases, transferases, ligases, and kinases, were the most frequently occurring *N. benthamiana* homolog genes in modules 1–2. Module 3 (12 nodes and 59 edges), the smallest module, had a low average degree and characteristic path length of 9.80 and 1.11, respectively, but a high clustering coefficient of 0.91, indicating a low degree of connections. The gene regulatory networks of the expression pattern E and F produced 3 GLay modules each (Figs. 16 and 17). The overrepresented GO-enriched terms in each of them were assessed. For modules in pattern E, module 1 is enriched with GO-BP terms associated with the regulation of plant seed and wax processes including “regulation of seed development”, “regulation of wax biosynthetic process”, “regulation of ABA biosynthetic process” and “regulation of seed dormancy process” which has been involved in hormone signaling such as strigolactones, gene expression, and control water uptake during the grafting process [118] and other roles in seed development, dormancy, and plant stress response [119–121] The main hub genes identified were, *Nbe.v1.s00030g01150* (a homolog of *PPR19*) and *Nbe.v1.1.chr06g38370* (a homolog of *RBL12*) (Fig. 15, Supplementary Dataset S6: Sheet 35 and Supplementary Dataset S7: Sheet 41). Biological processes related to cellular processes involving molecule modification and signaling pathways such as “histone H3-K9 acetylation”, “histone H3-K9 modification”, “chloroplast-nucleus signaling pathway”, “tertiary alcohol biosynthetic process” and “apocarotenoid biosynthetic process” are captured in modules 2 and auxin metabolism and biosynthesis GO-BP terms including “indoleacetic acid metabolic process”, “indoleacetic acid biosynthetic process”, “auxin biosynthetic process”, “indole-containing compound biosynthetic process” and “auxin metabolic process” in module 3 (Fig. 15, and Supplementary Dataset S6: Sheet 35). The top hub genes identified in modules 2 and 3 were two protein molecules, *Nbe.v1.1.chr17g06980* (a homolog of *NODULE INCEPTION-LIKE PROTEIN 4*, *NLP4*) and *Nbe.v1.1.chr03g40930* (a homolog gene encoding the putative methyltransferase family protein), respectively (Supplementary Dataset S7: Sheet 41).

**Figure 15.**
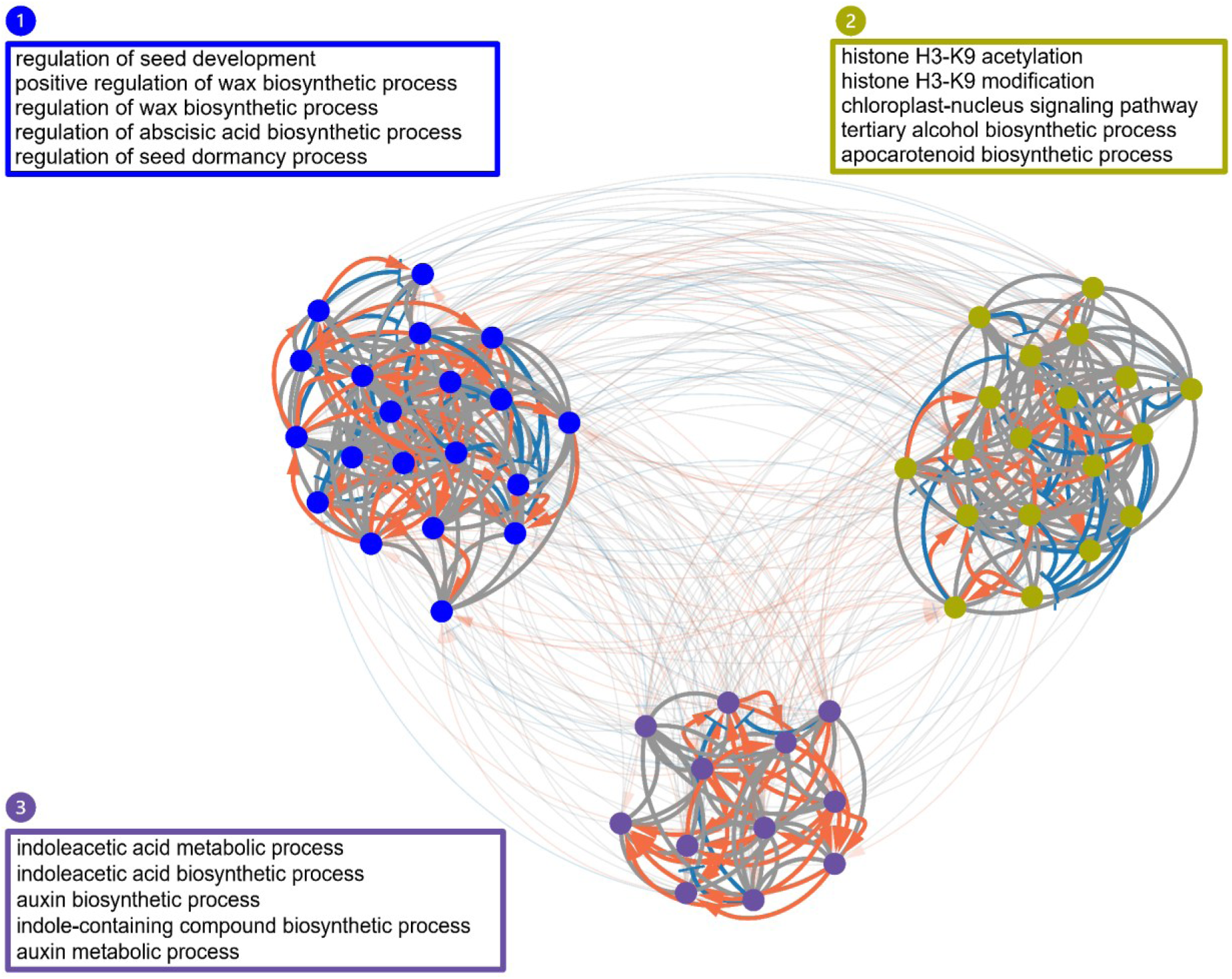
GLay module analysis of the Bayesian network for expression pattern E. The expression pattern E gene network was clustered into a total of three modules by the GLay community clustering algorithm in Cytoscape. GO enrichment analysis revealed that the modules identified had functionally distinct biological process GOs, including “regulation of seed development”, “histone H3-K9 acetylation”, and “indoleacetic acid metabolic process”. Subnetworks are indicated by different colors and numbered sequentially as detected by the algorithm. Red arrows indicate gene up-regulation. Blue T-lines indicate gene down-regulation. Gray lines indicate uncertain gene regulation. See Supplementary Dataset S6: Sheet 35 for details.

**Figure 16.**
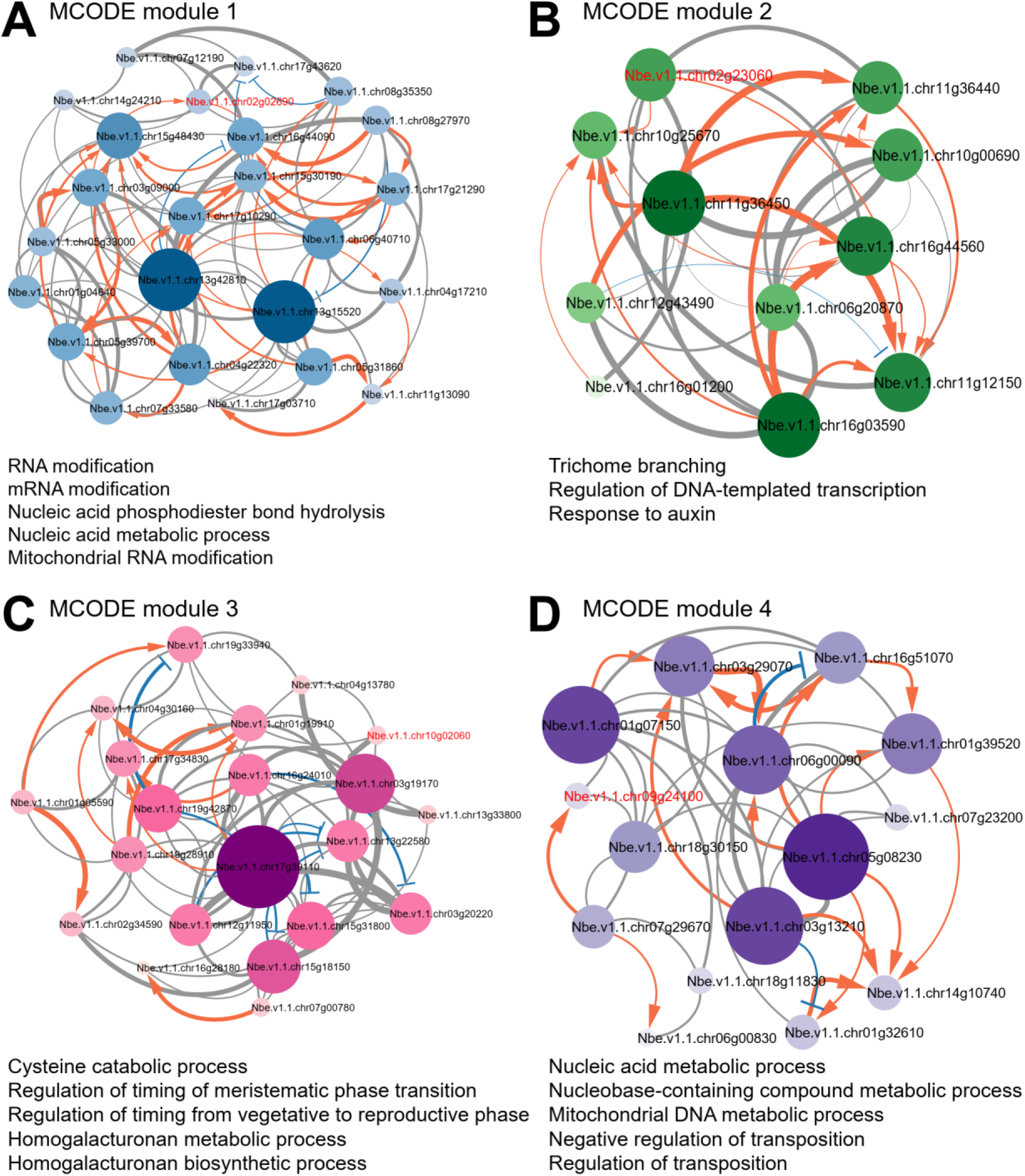
MCODE module analysis of the Bayesian network for expression pattern F. The module analysis using MCODE resulted in 4 significant modules, which include 71 nodes and 251 edges, and the enrichment of several GO-BP terms. Modules 1 – 4 are the top four modules identified. **(A)** Module 1 TPR-like complex) GO annotation has shown “RNA modification”, “mRNA modification”, and “nucleic acid phosphodiester bond hydrolysis”. **(B)** Module 2 (LYSM-CONTAINING RECEPTOR-LIKE KINASE 5 complex) GO annotation has revealed “trichome branching”, “regulation of DNA-templated transcription”, and “response to auxin”. **(C)** Module 3 (Terpenoid cyclases complex) GO annotation has shown “cysteine catabolic process”, “regulation of timing of meristematic phase transitions”, and “regulation of timing of transition from vegetative to reproductive phase”. **(D)** Module 4 (Transcription initiation factor TFIID subunit A complex) GO annotation has shown “nucleic acid metabolic process”, “nucleobase-containing compound metabolic process”, and “mitochondrial DNA metabolic process”. The seed node responsible for forming each module is indicated by red font. The module networks were visualized by Cytoscape (v3.10.2) by mapping the “degree parameter” to node size and color such that higher-degree nodes are darker in color and bigger in size. Red arrows indicate gene up-regulation. Blue T-lines indicate gene down-regulation. Gray lines indicate uncertain gene regulation. Line thickness indicates significance in gene regulation. All nodes are significantly enriched at p-value < 0.01. See Supplementary Dataset S5: Sheet 30 for details.

**Figure 17.**
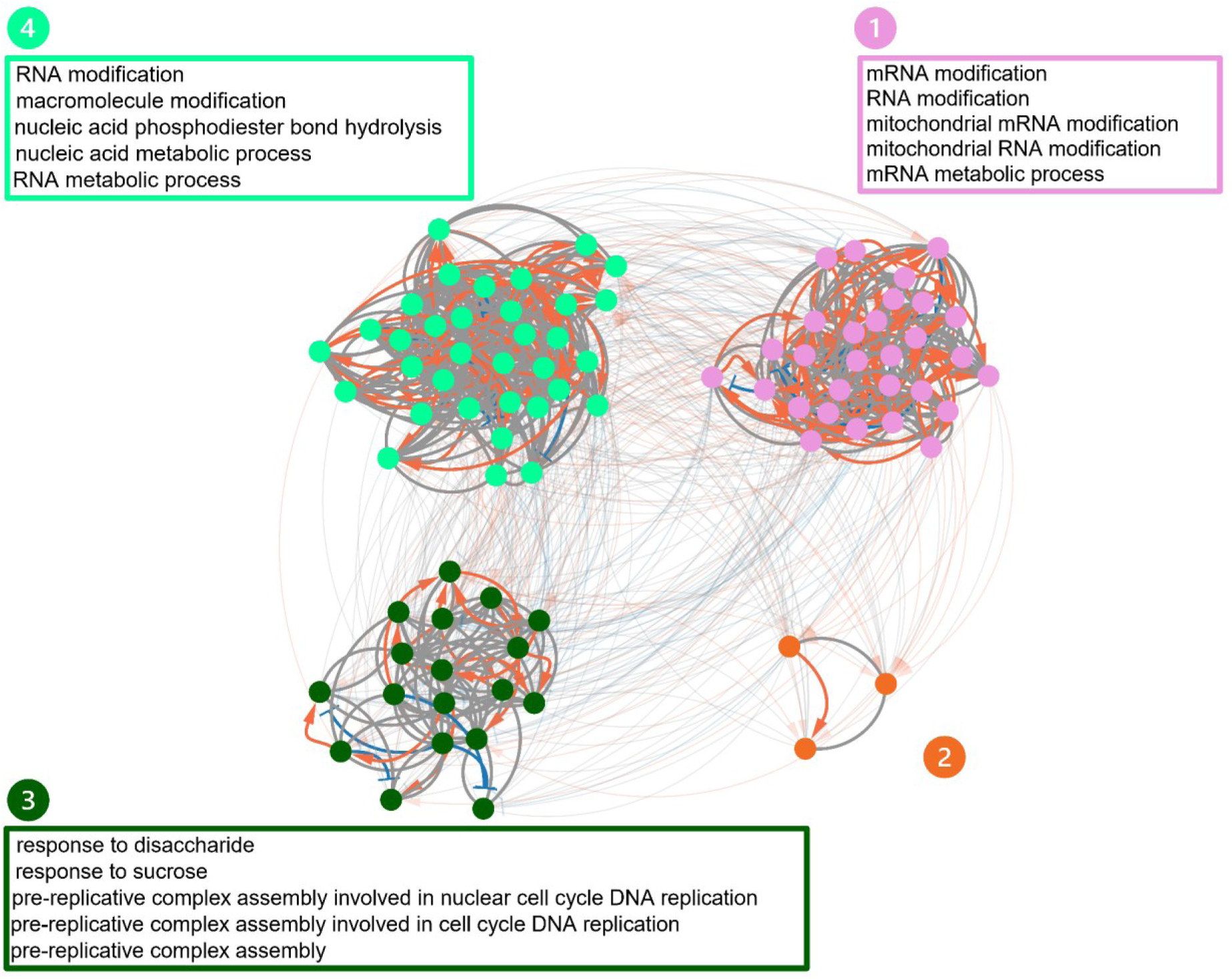
GLay module analysis of the Bayesian network for expression pattern F. The expression pattern F gene network was clustered into a total of four modules by the GLay community clustering algorithm in Cytoscape. GO enrichment analysis revealed that the modules identified had functionally distinct biological process GOs, “mRNA modification”, response to disaccharide”, and “RNA modification”. Subnetworks are indicated by different colors and numbered sequentially as detected by the algorithm. Red arrows indicate gene up-regulation. Blue T-lines indicate gene down-regulation. Gray lines indicate uncertain gene regulation. See Supplementary Dataset S6: Sheet 36 for details.

#### Pattern F

In pattern F, MCODE revealed a total of 5 modules (78 nodes and 263 edges, leaving 7 nodes unclustered). The significant MCODE modules with a threshold score > 5 were 4 and comprised 71 nodes and 251 (Fig. 16 and Supplementary Dataset S5: Sheet 30). The types of *N. benthamiana* homolog genes that formed the modules in the pattern mainly included cell cycle and division proteins (such as origin recognition complex subunit 3 and cyclin A3;4); transport proteins (such as K^+^ uptake transporter 3, potassium transporter 1 and MATE efflux family protein); signaling proteins (such as poltergeist-like 4, CRINKLY4 related 3 and PB1 domain-containing protein tyrosine kinase); transcription factors (such as auxin response factor 16, GATA transcription factor 11 and RNA polymerase sigma subunit 2) and enzymes (such as Zinc-binding alcohol dehydrogenase family protein, FIDGETIN-LIKE-1 INTERACTING PROTEIN (*FLIP*) and proteolysis 6). A total of 4 significant MCODE modules were identified in the gene network of expression pattern F, which has genes predominantly lowly expressed in the second (Ave 1–7 DAG) and third states (Ave 14–28 DAG). The seed node of module 1 was *Nbe.v1.1.chr02g02690* (a homolog gene encoding TPR-like superfamily protein) (Fig. 16A and Supplementary Dataset S5: Sheet 30), which is involved in various plant processes such as stress responses [122,123], hormone signaling [124], and chloroplast biogenesis [125]. This module was mainly enriched in GO-BP terms, “RNA modification”, “mRNA modification”, and “nucleic acid phosphodiester bond hydrolysis”. Hub genes in module 1 included *Nbe.v1.1.chr13g15520* (a homolog gene encoding mitochondrial transcription termination factor family protein, *MTERF19*) and *Nbe.v1.1.chr13g42810* (a homolog gene encoding the alkaline-phosphatase-like family protein). The seed node of module 2 was *Nbe.v1.1.chr02g23060* (a homolog gene encoding LysM-containing receptor-like kinase 5, *LYK5*) (Fig. 16B and Supplementary Dataset S5: Sheet 30), involved in the regulation of signaling pathways of plant hormones, regulation of metabolic and cellular processes [126], and abiotic stresses [127]. Hub genes in module 2 included *Nbe.v1.1.chr16g03590* (a homolog gene encoding *FLIP*) and *Nbe.v1.1.chr11g36450* (a homolog of *GATA TRANSCRIPTION FACTOR 11*, *GATA11*), however, there was no enrichment for this module due to the small number of nodes (11 nodes). The seed node of module 3 was *Nbe.v1.1.chr10g02060* (a homolog of beta-amyrin synthase, *BAS*) (Fig. 16C and Supplementary Dataset S5: Sheet 30), which plays a key role in the production of terpenoid natural products in plants [128,129]. This module was mainly enriched in GO-BP terms, “cysteine catabolic process”, “regulation of timing of meristematic phase transition”, and “transition from vegetative to reproductive phase”. The hub gene in module 3 was *Nbe.v1.1.chr17g39110* (a homolog *GT2*). The seed node of module 4 was *Nbe.v1.1.chr09g24100* (a homolog of TBP-associated factor 12b gene, *TAF12b*), which is crucial for gene expression regulation and RNA polymerase II-induced transcription initiation in plants [130]. This module was mainly enriched in GO-BP terms, “nucleic acid metabolic process”, “nucleobase-containing compound metabolic process”, and “mitochondrial DNA metabolic process”. The hub gene in module 4 was *Nbe.v1.1.chr05g08230* (a homolog of proteolysis 6 gene, *PRT6*) (Fig. 16D and Supplementary Dataset S5: Sheet 30). Details of genes in the subnetworks of the MCODE modules of the expression patterns A–F are shown in Supplementary Dataset S5: Sheet 25–30.

GLay community clustering analysis in pattern F revealed 3 large modules and a small module (Fig. 17). Module 1 (29 nodes and 211 edges), Module 3 (18 nodes and 81 edges) and Module 4 (34 nodes and 270 edges) were moderate-sized, densely connected, and highly efficient network structures with similar characteristics, including the average number of neighbors, characteristic path length, clustering coefficient and network density of approximately 13.14, 1.49, 0.63 and 0.51. Module 2 (3 nodes and 3 edges) was the smallest GLay module in the gene regulatory network of expression pattern F and was made up of *Nbe.v1.1.chr01g19910* (a homolog of *TOXICOS EN LEVADURA 4*, *ATL4*), *Nbe.v1.1.chr15g48430* (a homolog gene encoding DUO1-activated unknown 1, *DAU1*) and *Nbe.v1.1.chr04g30160* (a homolog of the potassium transporter 1 gene, *KT1*), suggesting it has a potential function in toxicity responses, RNA processing, and ion transport in plants. For modules in pattern F, module 1 is enriched with GO-BP terms associated with RNA modification including “mRNA modification”, “RNA modification”, and “mitochondrial mRNA modification” whereas carbohydrate response and pre-replicative complex assembly terms including “response to disaccharide”, “response to sucrose” which have been involved in grafting [131,132], pre-replicative complex assembly and pre-replicative complex assembly involved in “nuclear cell cycle DNA replication” were enriched in module 3 (Fig. 17 and Supplementary Dataset S6: Sheet 36). The top hub genes identified in modules 1 and 3 were *Nbe.v1.1.chr05g08230* (a homolog of *PRT6*) and *Nbe.v1.1.chr03g19170* (a homolog gene encoding xylulose kinase-1, *XK-1*), respectively (Supplementary Dataset S7: Sheet 42). Module 4 is enriched in GO-BP terms associated with RNA processing terms, including “RNA modification”, “macromolecule modification”, “nucleic acid phosphodiester bond hydrolysis”, “nucleic acid metabolic process”, and “RNA metabolic process” with the hub gene, *Nbe.v1.1.chr16g03590* (a homolog of *FLIP*) (Fig. 17, Supplementary Dataset S6: Sheet 36 and Supplementary Dataset S7: Sheet 42). Collectively, our module analysis using both MCODE and GLay algorithms revealed the abundance of organized subnetworks or modules of distinct and overlapping biological processes and hub genes that potentially have important roles in each biological process. Details of genes in the subnetworks of the GLay modules of the expression patterns A–F are shown in Supplementary Dataset S6: Sheet 31–36. It was observed after the analysis that MCODE produces much smaller modules than GLay, leaving the majority of the nodes unclustered. Therefore, GLay outperforms MCODE in terms of structural partitioning of the original gene regulatory networks analyzed. GLay has higher sensitivity in identifying functional modules than MCODE, which has higher specificity in identifying more tightly connected modules. Overall, while MCODE’s specificity excels at identifying highly specific, tightly connected modules, GLay’s higher sensitivity makes it more suitable for broader functional interpretation of biological networks.

## Discussion

In the present study, we identified distinct gene expression patterns and gene modules to reveal biological processes associated with graft union formation in the *N. benthamiana*/*A. thaliana* interfamily grafting. Moreover, after clustering genes by their up- and down-regulation patterns in the three states, genes in each expression pattern were further analyzed through Bayesian network analysis using the SiGN-BN HC + Bootstrap program [34,35]. Subsequently, two types of network clustering algorithms, MCODE [64] and GLay [65], were used to identify the core modules. In each module, hub genes that are potentially critical for the module were identified using the degree algorithm of the CytoHubba plugin [43] in Cytoscape software (v3.10.2) [40].

### Dynamics of gene expression after grafting

We started the processing of the RNA-seq data from the *N. benthamiana*/*A. thaliana* interfamily graft samples into the gene expression levels (TPM) with a newly updated genome database (Nben v1.1, [37]). The datasets were pre-processed, and PCA was utilized to estimate the similarity between the intact and grafted samples based on distance threshold relationships. From the total gene expression levels (TPM) calculated, distance threshold-based clustering was performed, and six distinct temporal gene expression patterns A–F were identified, which also ensured narrowing down the number of target genes for network estimation. A greater number of genes were up-regulated and classified into patterns A–C (Fig. 1C, D). In contrast, down-regulated genes classified into patterns D–F were only 11.1% (Fig. 1C, D). The genes present in patterns A-C include genes associated with wounding and immunity, indicating that wound damage response and pre-immune reaction rely on transcriptional regulation. A previous study revealed that autophagy is required for decay of the previous cell state during tissue regeneration after callus induction [133]. Proteomic exchanges caused by auxin and abscisic acid treatment were drastically decreased by shifting plants to callus induction medium in wild-type plants, but less so in autophagy mutants. Autophagy action was also observed during grafting [37]. Autophagy was activated in cells near the graft boundary in the scion side, where callus tissues were mainly formed to promote tissue reconnection. Analysis of autophagy mutants indicates its role in callus cell proliferation during grafting. Based on these observations, it is possible that degeneration and regeneration by decay of previous cell states rely on proteomic exchange through actions such as autophagy and proteasome rather than transcriptional regulation.

Patterns A–C following the enriched GO-BP terms and hub genes identified in the functional modules indicated events that were promoted during grafting to drive tissue reunion and successful graft union formation, including wound response, cell division and differentiation, cell adhesion, hormone signaling, and vascular formation. Meanwhile, pattern D–F indicated events that were suppressed after grafting, representing a trade-off that prioritizes regeneration and integration over normal physiological activities such as photosynthesis and chloroplast activity, mitochondrial energy chain, nutrient uptake and assimilation, flowering, DNA recombination and epigenetic silencing, protein degradation, and non-essential stress adaptation pathways.

GO terms enrichment was further performed by analysis of gene annotations in each expression pattern. From the analysis, pattern A (early and continuous upregulation response) functions in the initiation and maintenance of graft union formation initiated early events such as wound and immune response (NB-ARC domain-containing disease resistance protein gene and *WRKY6*), cell wall remodeling (*β-1,3-glucanases*), plasmodesmata connectivity (*PDLP2*) and late events such as vascular development (*VND1, VND7*, *PIN3* and *PIN7*). Pattern B (transient early upregulation response) functions in rapid early-stage events such as wound repair and defense priming (*TLP-3*), auxin signaling (*CNGC14*), cell cycle process (*CYCD2;1* and *CYCA2;2*), and cell wall modification (*PRX17*). These processes in pattern B are downregulated in the late stage to transition from tissue proliferation events to tissue maturation events. Pattern C (late upregulation response) are more involved in tissue maturation and long-term union cohesiveness and stabilization events such as stress adaptation (prolyl oligopeptidases family protein gene), cell wall strengthening (*IRX15-L* and pectin lyase-like superfamily protein gene), hormonal coordination (*BES1*, *CRF4*, *PIN3* and *PIN7*) and RNA processing and peptide regulation (Zinc knuckle family protein genes and *TRZ4*). Among suppressed expression patterns D–F, pattern D (late downregulation response) which functions in the suppression of growth-related and energy-intensive events were involved in the suppression of cell cycle and DNA replication (downregulation of *CYCA3;4*), photosynthesis and energy chains (downregulation of *PHT3;1*, *XK1*, *GPT2* and K+ uptake transporter 3 gene) and restriction of cell wall remodeling and excessive cell wall degradation (downregulation of *BXL6*). Pattern E (early downregulation response) suppresses reproductive and lignification processes by inhibiting flowering and senescence pathways, mitochondrial functional regulation, non-essential translation processes and protein synthesis, nitrate uptake and transport, lignin biosynthesis following the downregulation of key genes including *NAC075*, *RAP2.7*, *RBL12*, *PPR19*, *BXL6,* and *NLP4*. Pattern F (continuous downregulation response) reflects an overall shift in energy-intensive and developmental processes in favor of graft healing and tissue regeneration by suppressing events including mitochondrial and chloroplast gene expression, homologous recombination, protein degradation, chlorophyll biosynthesis, chloroplast development, photosynthesis, light signaling, photomorphogenesis, nutrient uptake and plant growth following the repression of the genes, *MTERF19*, *SLG1*, *GT2, PRT6, SIG2*, alkaline-phosphatase-like family protein gene, *GATA11* and *FLIP*.

### Modules and hub genes in the gene network of *N. benthamiana*/*A. thaliana* grafting

In this current network analysis, the SiGN-BN HC + Bootstrap program, a gene network estimation software that uses Bayesian network models and a B-spline nonparametric regression to estimate regulatory dependencies between genes and model parent-child relationships as gene networks from gene expression data, was used [34,35] To ensure that genuine biological relationships among genes in each expression pattern are estimated following the Bayesian networks estimation, p-values and population sizes of the gene regulatory networks of expression patterns were used as criteria to eliminate possible artificial bias that could arise from statistical fluctuations. This approach increased the reliability and robustness of the inferred gene regulatory networks and minimized spurious associations. Further, a series of gene network topological analyses was performed to identify the gene modules in each gene network using two module clustering algorithms, MCODE and GLay. The MCODE algorithm identified small and tightly connected gene modules within the network. The nodes had many strong connections to each other and represented molecular complexes likely to be functionally related. However, the MCODE algorithm left some of the nodes unclustered. Meanwhile, the GLay clustering algorithm uncovered larger, more loosely connected modules spanning multiple biological functions by considering the overall structure and connections within the network. The GLay algorithm divided the network into groups by iteratively removing edges that connect different communities, resulting in a hierarchical structure of modules. As a result, some GLay modules were distinct, while others were nested within larger structures. This hierarchical organization helps in understanding how genes function in both small, specialized modules and broader, more general biological processes.

With the MCODE algorithm, the number of modules identified from the gene expression patterns A–F was 5,4,6,4,3, and with the GLay algorithm, 4, 4, 4, 3, 3 modules in the gene network of expression patterns A–F, respectively. In these modules, we identified hub genes corresponding to established sequential grafting processes associated with upregulated expression patterns A–C (Fig. 18) and the non-essential trade-off events to grafting in downregulated expression patterns D–F (Fig. 19). The established plant grafting processes identified to be associated with upregulated expression patterns A–C include immune response, hormone signaling, cell wall biogenesis, cell division/proliferation, and vascular differentiation (Fig. 18). We identified a total of 47 hub genes potentially associated with these established early and later developmental stage processes in *N. benthamiana*/*A. thaliana* grafting. Interestingly, these genes are being described as graft-related for the first time, except for one (*LBD4*), which has previously been predicted as a hub gene potentially involved in tomato and pepper graft vascular cell proliferation (callus production) [7]. In expression pattern A, a total of 5 immune-related hub genes were identified (NB-ARC protein gene, *RAF36*, *RFO3*, *LECRK-VII.1*, and *LOL2*), 4 hormone signaling hub genes (*GPCR*, *BRL1*, *PAS1*, and *UGT74F1*), 3 hub genes involved in cell wall biogenesis (*NAC073*, *ATL54*, and *PGSIP3*), and 7 cell division/proliferation hub genes (*MDKIN1*, *PAS1*, *BRL1*, *LBD4*, *TPLX3*, *RIMA*, and *ORC6*). Pattern B contained 4 immune response hub genes (NB-ARC protein gene, PADRE protein gene, NBS-LRR protein gene, and *ZAR1*), 1 hormone signaling hub gene (*CNGC14*), 1 cell wall biogenesis hub gene (*FAS4*), and 6 cell division/proliferation hub genes (*PAKRP2*, *KINESIN-12E*, *CDC45*, *CYC3B*, RNI-like protein gene, and *FKBP43*). Pattern C included 5 immune response hub genes (*NHL8*, *SYCO*, *ZAT14*, NBS-LRR protein gene, and *SDA1*), 4 hormone signaling hub genes (*ITPK1*, *FL8*, *JMJ20*, and *ADTO1*), 1 cell wall biogenesis gene (*LUH*), 3 cell division/proliferation hub genes (*ATXR6*, *DMC1*, and *TPXL3*), and 3 vascular differentiation genes (*FL8*, *PP2-B1*, and *ADTO1*) (Fig. 18). These hub genes may collectively coordinate the regulation of downstream genes and exhibit their roles essential for successful grafting. On the other hand, we identified hub genes that are involved in non-essential trade-off events to grafting and downregulated to ensure resource allocation for improved grafting. The non-essential trade-off events identified to be associated with the downregulated expression patterns D–F include photosynthesis, excessive cell wall modelling, reproductive process, mitochondrial process, and nutrient uptake (Fig. 19). These hub genes are potentially associated with photosynthesis (*XK1*, *GATA11*, *GT2,* and *SIG2*), excessive cell wall remodeling (*BXL6* and *TBL41*), reproductive process (*NAC100* and *FLIP*), mitochondrial process (*PHT3;1*, *RBL12*, *MTERF19,* and *SLG1*), and nutrient uptake (*NLP4*) (Fig. 19). Further detailed discussion with information on each gene function that has been revealed in previous studies is described in Supplementary Text 1.

**Figure 18.**
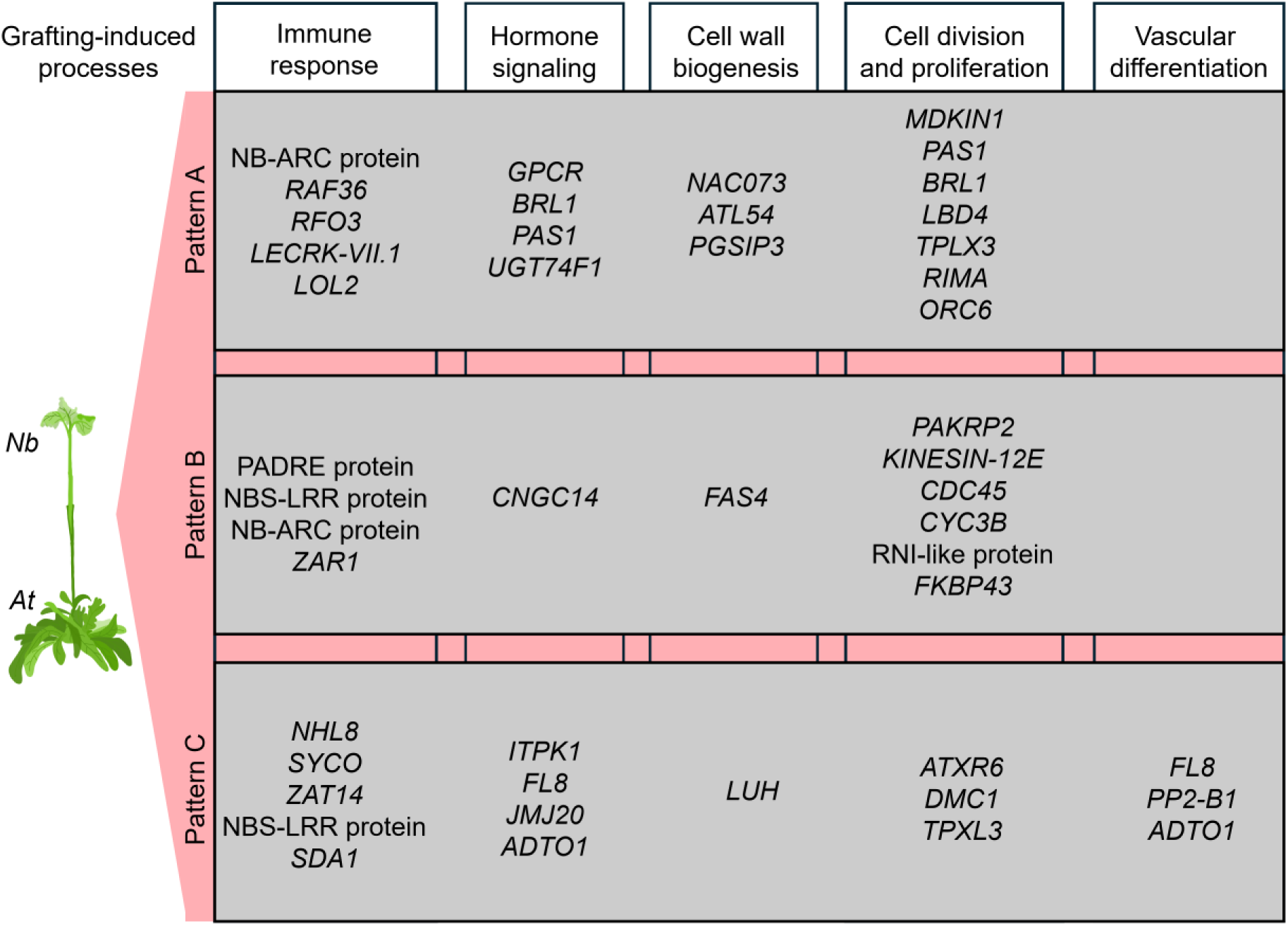
Network hub genes upregulated during grafting in MCODE and GLay modules for patterns A–C. The identified hub genes are potentially associated with early-stage processes, including immune responses such as disease resistance (NB-ARC domain-containing disease resistance protein gene, *RAF36*, *RFO3*, *LECRK-VII.1*, PADRE protein gene, CC-NBS-LRR class protein gene, *ZAR1*, *NHL8,* and *SDA1*) and cell death (*LOL2* and *ZAT14*). Hub genes potentially involved in later developmental stage processes were identified, including hormonal signaling (*GPCR*, *BRL1*, *PAS1*, *UGT74F1*, *CNGC14*, *ITPK1*, *FL8*, *JMJ20* and *ADTO1*), cell wall biogenesis (*NAC073*, *ATL54*, *PGSIP3*, *FAS4* and *LUH*), cell division and proliferation (*MDKIN1*, *LBD4*, *TPLX3*, *RIMA*, *ORC6*, *PAKRP2*, *KINESIN-12E*, *CDC45*, *CYC3B*, RNI-like protein gene, *FKBP43*, *ATXR6* and *DMC1*) and vascular development (*FL8*, *PP2-B1* and *ADTO1*).

**Figure 19.**
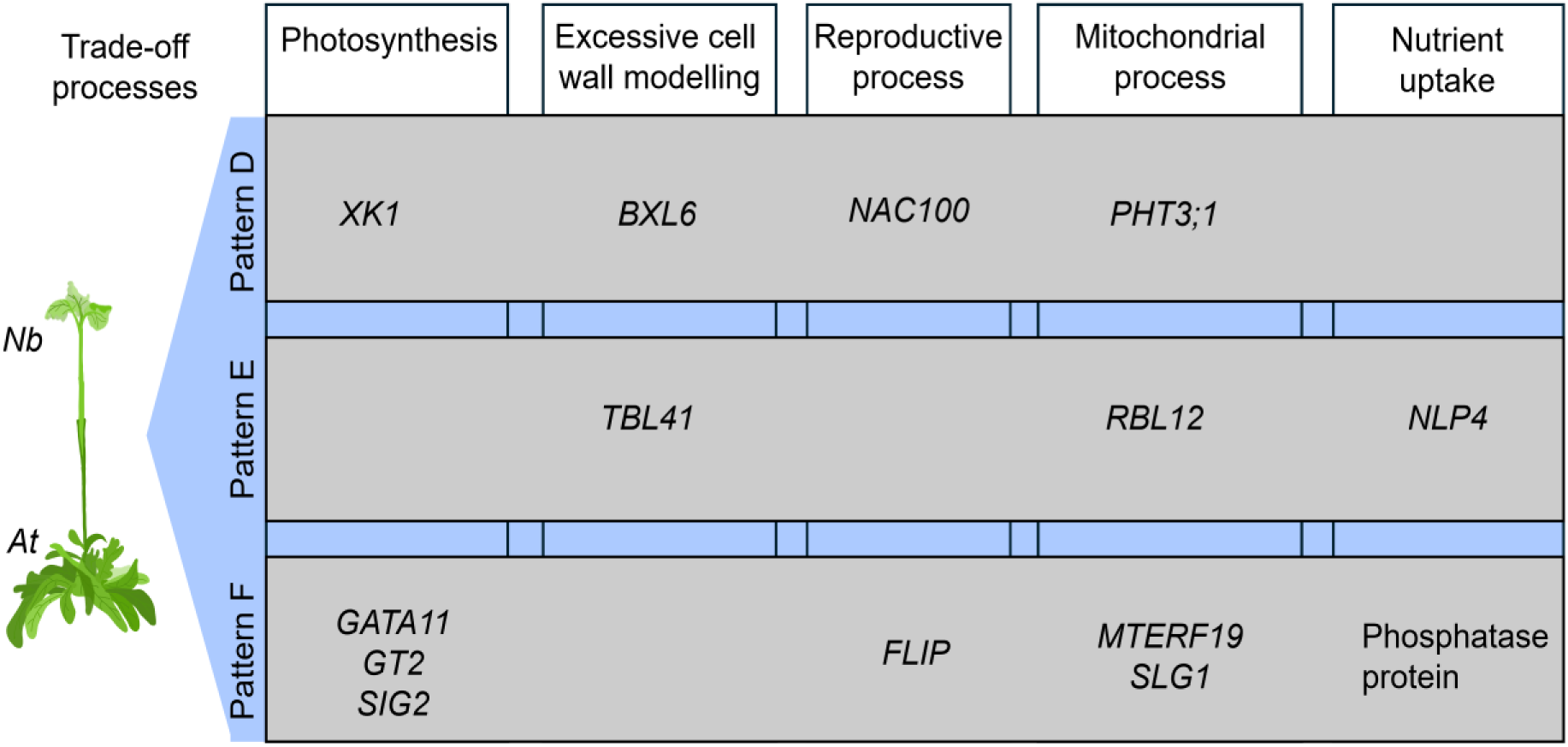
Network hub genes repressed during grafting in MCODE and GLay modules for patterns D–F. The hub genes are potentially associated with non-essential trade-off events to grafting, including photosynthesis (*XK1*, *GATA11*, *GT2*, and *SIG2*), excessive cell wall remodeling (*BXL6* and *TBL41*), reproductive process (*NAC100* and *FLIP*), mitochondrial process (*PHT3;1*, *RBL12*, *MTERF19*, and *SLG1*), and nutrient uptake (*NLP4* and phosphate protein gene).

### Stringent distance thresholds exclude established grafting genes from network analysis

Our network analysis successfully identified key gene modules and hub genes associated with grafting responses across distinct expression patterns. However, it is important to note that several previously characterized grafting-related genes including aberrant lateral root formation 4 (*ALF4*), auxin-resistant 1 (*AXR1*), transport inhibitor response 1 (*TIR1*), auxin signaling f-box 2 (*AFB2*) [134], auxin response factors (*ARF8*) [135], cellulase3 (*CEL3*/*GH9B3*) [17], target of monopteros6 (*TMO6*), pectin methylesterase inhibitor 5 (*PMEI5*), DNA binding with one finger (*DOF2*.1 and *DOF6*), high cambial activity2 (*HCA2*), KORRIGAN1 (*KOR1*) [136], NAC domain containing protein 71 (*ANAC071*) [22,137,138], knotted-like from A. thaliana (*KNAT1*), LOB domain-containing protein 25 (*LBD25*), LOB domain-containing protein 18 (*LBD18*), myb domain protein 86 (*MYB86*), NAC secondary wall thickening promoting factor (*NST1*) [7], phytochrome a signal transduction1 (*PAT1*) [139], xyloglucan endotransglucosylase/hydrolases family (*XTH28*) [17], 1-amino-cyclopropane-1-carboxylate synthase 2 (*ASC2*) [22], wound-induced dedifferentiation 1 and 4 (*WIND1*) [140–142] and WUSCHEL-related homeobox (*WOX4* and *WOX13*) [7,143] were not included in the network analysis as 2,808 genes with high expressional changes during grafting were selected in this study (Fig. 1C). These genes were indeed present in the initial expression dataset; however, the stringent distance threshold-based cutoff criteria applied resulted in their exclusion from subsequent network analysis. Despite their exclusion from the main network analysis, we have included the expression profiles of these genes to provide insight into their transcriptional dynamics (Supplementary Fig. S1). This approach highlights the advantage of balancing stringent threshold cutoffs for high-confidence gene selection with the inclusion of a broader range of biologically relevant genes. Alternative threshold-based strategies and integrative approaches using genes specifically associated with each biological process for network analyses will provide further knowledge on the molecular mechanism of grafting from multiple aspects.

### Graft-related genes identified in this study

In this study, we aimed to capture a long-term transcriptional dynamic shift from ground state (intact stem, before grafting) to second and third states, corresponding to the first week after grafting and two or later weeks after grafting, respectively, in the *N. benthamiana*/*A. thaliana* interfamily grafts. In view of the morphological and cytological observations during grafting [110,111,144], we considered that the second state corresponds to events involved in the primary wound healing process, and the third state reflects events related to the latter coordination process to reestablish and retain homeostasis of stem function. This study identified core gene regulatory modules as well as hub genes connecting a higher number of genes associated with each biological event. As a result, this study captured novel sets of graft-related genes in gene modules (Fig. 4–17), helping our construction of an entire view of grafting processes.

Genes associated with defense responses to abiotic and biotic stresses were induced during grafting. The induction of ankyrin repeat family protein (*Nbe.v1.1.chr18g27360*), heat shock proteins (*HSP17.6A* and *HSP70*), NB-ARC domain-containing disease resistance protein (*Nbe.v1.1.chr01g00700*), salicylic acid-related genes (*UGT74F1*), osmotic-stress gene (*OSCA2.*3), ROS-mediated and ABA signaling related-genes (*ABO5*, *GPCR*, *MC3, MPK9*, *MTERF19*, *PP2C, PRT6* and *RAF36*), pathogenesis-related protein (*TLP-3*), WRKY transcription factor (*WRKY6*) and wound-responsive proteins (*WIP3* and *BBD2*) during grafting highlights a critical overlap between biotic (pathogen and immune defense activation) and abiotic (dehydration, oxidative, and heat) stress responses and grafting-induced immunity. This crosstalk suggests that grafting co-upregulates conserved stress adaptation mechanisms to facilitate graft healing and tissue regeneration while initiating defense pathways.

Moreover, graft union formation potentially shifts the transcriptional reprogramming to support rapid cell division and differentiation, which may be accompanied by the utilization of sugars and hormone metabolism. Cell division and differentiation genes such as *CDC45*, *CYCD2;1*, *CYCA2;2*, *CYC3B*, *PAS1*, *PP2-B1,* and *RIMA* were identified in the upregulated expression patterns, suggesting promoted meristematic activity and vascular tissue differentiation at the graft site. This proliferative response is accompanied by enhanced sugar utilization with the induction of sugar transporters (*STP14*) and hormonal metabolism, particularly auxin (*PIN3*, *PIN7*, *FL8*, *IAA17* and *DFL1*), brassinosteroid-related protein (*BRL1*), ethylene (*ERF1* and *ERF106*), cytokinin (*CRF4*), and jasmonic acid (*LECRK-VII.1*) which were activated probably to coordinate tissue regeneration and stress responses. In contrast, repressed genes linked to photosynthesis, mitochondrial energy production, and the electron transport chain, nutrient uptake, and assimilation suggest a temporary suppression of normal cellular activities, which likely serves as an energy redirection strategy towards graft healing processes.

Further, while our Bayesian network analysis revealed regulatory cascades including NAC, WRKY, AP2/ERF, MYB, and bHLH transcription factors, many members of these cascades may be explored in *N. benthamiana* interfamily grafting because, in this system, many active and longer responses occur to overcome a less compatible situation rather than conventional compatible homo- or closed relatives-grafting. For example, although NACs such as *NAC071* and *NAC096* are known to regulate vascular development in other plant species, including *A. thaliana* (Asahina, 2011), many of these NACs, including *NAC007, NAC02, NAC04, NAC036, NAC060, NAC073, NAC53, NAC076,* and *NAC083* identified in our network, lack known roles in grafting, suggesting hindered regulatory activity in grafting.

### Hormone signaling during graft formation

Phytohormones play key roles in graft formation. The hormone-associated hub genes identified include auxin-related genes (*CNGC14, PAS1, FL8,* and *ITPK1*), brassinosteroid-related genes (*BRL1*, *ADTO1*), gibberellic acid-related gene (*JMJ20*), ABA signaling (*GPCR*), and salicylic acid-related gene (*UGT74F1*). Auxin is a phytohormone central to graft formation, playing critical roles in cell division, callus formation, vascular differentiation, and tissue reunion [NO_PRINTED_FORM]. The identification of these auxin-related hub genes highlights the importance of auxin response in interfamily grafting. Previous studies have shown that grafting induces the expression of auxin biosynthesis and signaling genes [137,145–148], where it promotes vascular regeneration and callus proliferation. This is evident in delayed *A. thaliana* hypocotyl graft healing and phloem reconnection due to the disruption of auxin transport or signaling by the removal of cotyledons or application of auxin transport inhibitor, triiodobenzoic acid [137]. The upregulation of transcription factors in auxin signaling is essential for cell proliferation during tissue reunion [138]. Above the grafting junction, activation of transcripts encoding efflux carriers in auxin transport has been identified to regulate graft development [19].

This study identified BRL1, which plays a partially redundant role with a Brassinosteroid (BRs) receptor, BRI1, implying the occurrence of BR signaling to facilitate graft formation. BRs are plant steroid hormones that regulate cell growth and morphogenesis as well as responses to both biotic and abiotic stresses [149,150]. BR biosynthesis and signaling induce xylem formation, and they interact with other essential grafting phytohormones such as auxin for overall graft success, such that inhibition of BR biosynthesis represses tracheary element differentiation in *Zinnia elegans* cells [151] and represses the formation of secondary xylem in cress plants [152]. Also, BR accumulation has been identified to peak before tracheary element differentiation [153]. Ethylene is an endogenous stimulator that promotes secondary growth [154] and likely acts synergistically with auxin to promote vascular cell divisions [110]. This synergy is mediated in part through the receptor kinase *PHLOEM INTERCALATED WITH XYLEM* (*PXY*)/*TRACHEARY ELEMENT DIFFERENTIATION INHIBITOR FACTOR* (*TDIF*) *RECEPTOR* (*TDR*), which promotes vascular cell proliferation by regulating the auxin-inducible transcription factor *WOX4*, a key component of the *PXY*/*TDR* signaling pathway [155,156]. During grafting and wounding, ethylene biosynthesis genes are activated at the graft junction [147] and around the wounding site [22,157], indicating the significance of ethylene signaling for tissue healing and reunion.

Gibberellic acid (GA) biosynthesis, represented by the hub gene *JMJ20*, may be a crucial graft candidate for promoting cell elongation, division, and vascular tissue development. GAs are diterpene phytohormones known to enhance xylogenesis in cambium tissue by accumulating in developing xylem tissues [158,159]. GA biosynthesis inhibition during grafting suppresses cortex cell expansion at the graft junction [137]. GAs have also been reported to modulate the turnover of PIN proteins, thereby promoting auxin transport [160,161], suggesting the complementary roles of these hormones in cell expansion, tissue proliferation, and reunion at the graft junction [134,137].

From the current study, the presence of *UGT74F1*, a salicylic acid (SA) metabolism-associated gene, suggests a potential role of SA in grafting. SA is an essential hormone tightly regulated to ensure plant immunity [162]. SA accumulates in both roots and leaves during grafting. This accumulation stimulates the biosynthesis of abscisic acid and hydrogen peroxide and upregulates the expression of cold-responsive (COR) genes, such as *CsCOR47* and *CsICE1,* in enhancing chilling tolerance in grafted cucumber and pumpkin [163,164]. The identification of multiple hormone-related hub genes from the Bayesian network modules highlights the complexity and precision of hormone regulations during graft healing and vascular reconnection.

## Conclusion

This study investigated the molecular events during plant grafting and identified crucial gene modules as key modulators in the process of *N. benthamiana*/*A. thaliana* interfamily grafting. The Bayesian network approach further explored the transcriptome including, (1) modules involved in cell wall biogenesis and defense responses essential for establishing a successful graft union and preventing infection, (2) modules related to cell cycle regulation and DNA replication critical for cell division and tissue regeneration at the graft interface, and (3) modules associated with RNA processing and protein modification important in regulating gene expression and protein function during the grafting process. We identified hub genes in the regulatory networks and module subnetworks involved in a range of these cellular and biological processes critical for plant growth, adaptation, and graft union formation. Overall, the study provided insight and enhanced our understanding of the crucial molecular events and biological processes during plant grafting. The key hub genes identified provide a foundation for further research and can serve as targets for novel grafting strategies and improvement. In the future, further investigations may provide us with new insights and elucidate the specific function of these genes involved in crucial biological processes such as “cell wall biogenesis”, “defense responses”, “DNA replication”, “RNA processing”, and “protein modification” in enhancing graft compatibility and accelerating graft union formation. This knowledge may inform the development of strategies, potentially through targeted genetic engineering, to improve plant grafting success. For example, the effect of these genes on graft union formation and success in plants can be observed by overexpressing or suppressing their expression. Moreover, network analysis focusing on the hub genes identified for module-specific sequential biological processes during grafting would help to describe the entire gene network or map for these events. In particular, applying a Bayesian network approach at defined upstream and downstream levels can provide a comprehensive map of the causal interactions and regulatory hierarchies in the modules involved in graft formation and development. This strategy will enhance our understanding of the detailed gene regulatory map that will reveal the causal hierarchies and interactions during grafting, providing a predictive framework for graft biology.

## Methods

### Identification of gene expression patterns during *N. benthamiana*/*A. thaliana* interfamily grafting

The RNA-seq dataset collected in a previous study [17] was used in this study. The dataset contained *N. benthamiana* intact stem samples and *N. benthamiana*/*A. thaliana* interfamily grafted stem samples from six time points, 1, 3, 5, 7, 14, and 28 days after grafting (DAG), with three respective biological replicates. Based on principal component analysis (PCA), the time-course samples were divided into three states: ground state (before grafting [intact]), second state (1, 3, 5, and 7 DAG), and third state (14 and 28 DAG). Then, six unique mathematical thresholds were set according to the changing gene expression profiles at the different time points (1, 3, 5, 7, 14, and 28 DAG) after grafting relative to the expression of the intact samples. To set the defined mathematical thresholds, the Log_2_TPM (Transcripts Per Million) values of genes were classified into 3 data points including “Intact” (data of intact plants, fixed at 1), “Ave 1–7 DAG” (average of 1–7 DAG data points), and “Ave 14–28 DAG” (average of 14 and 28 DAG data points). The six mathematical thresholds set were (A) Intact = 1 TPM; Ave 1–7 DAG ≥ 2 TPM; Ave 14–28 DAG > 1 TPM, (B) Intact = 1 TPM; Ave 1–7 DAG ≥ 2 TPM; Ave 14–28 DAG ≤ 1 TPM, (C) Intact = 1 TPM; 0.5 ≤ Ave 1–7 DAG ≤ 2 TPM; Ave 14–28 DAG ≥ 2 TPM, (D) Intact = 1 TPM; 0.5 ≤ Ave 1–7 DAG ≤ 2 TPM; Ave 14–28 DAG ≤ 0.5 TPM, (E) Intact = 1 TPM; Ave 1–7 DAG < 0.5 TPM; Ave 14–28 DAG ≥ 0.5 TPM, and (F) Intact = 1 TPM; Ave 1–7 DAG ≤ 0.5 TPM; Ave 14–28 DAG < 0.5 TPM. Genes with similar expression patterns were extracted using the thresholds set, and plots of the expression profiles were processed using the GraphPad Prism Software (Version 6.0, GraphPad Software, La Jolla, USA).

### Gene ontology enrichment analyses

For each distinct temporal gene expression pattern, gene ontology (GO) enrichment analyses were performed by GOATOOLS [38]. The most significant enriched GO terms (sorted according to their *P*-value) at *P* < 0.01 were selected by the biological process category. REVIGO was used to reduce redundancy and complexity in the GO terms, and the similarity between terms is reflected by the semantic space [39]. The most informative results were visualized as treemaps using the treemap package in R, with GO-related terms grouped in superclusters under a collective name.

### Construction of Bayesian gene network

Bayesian network analysis of genes belonging to each expression pattern was estimated on 120 RNA-seq data, collected in a previous study [37], using the SiGN-BN HC + Bootstrap program (http://sign.hgc.jp/signbn/index.html; [34,35]) implemented on a cluster server system. The dataset comprises time-series transcriptome profiles from site-specific sections of grafted and intact stem tissues providing temporal resolution of graft-induced gene expression and transcriptome profiles from tissues and organs of *N. benthamiana* across young to mature developmental stages including, the stem apex, cotyledons, hypocotyls, and roots at the seedling stage; young and fully expanded leaf apical regions; floral organs; stems; and roots at the mature plant stage. Next, we analyzed the gene network using the Cytoscape software (v3.10.2) [40].

### Topological analysis of networks and detection of hub genes

The Cytoscape plugin, Network Analyzer [41,42] was used for the analysis of the topological properties of each interaction network, including degree, clustering coefficients, centrality, distribution of node degrees, neighborhood connectivity, and average shortest path lengths. Hub genes are nodes (genes) with a higher degree of connectivity in a network [42]. In this study, to detect the hub genes in each interaction network, the node degree distribution [165] was calculated using the Network Analyzer plugin in Cytoscape software [40]. The genes with an interaction degree that lies over 0.95 quantiles of the degree distribution were selected as hub genes. The subnetwork of hub genes was constructed using the Degree algorithm of the CytoHubba plugin [43] in Cytoscape software (v3.10.2), with the degree of importance of hub genes depicted using a color scale ranging from red to yellow.

### Network module analysis

We aimed to identify and characterize the functional modules within our Bayesian regulatory networks. To identify the significant clustering modules in the Bayesian network of each expression pattern, we performed module analysis using two widely used classical network clustering algorithms, including MCODE (v2.0.3) [64] and GLay community clustering algorithm (v2.3.4) [65] in Cytoscape (v3.10.2). The MCODE identifies seeds (anchor genes) in each module, which are nodes with the highest weighted vertex (highest connectivity) for expansion by computing a score of local density for each node. Modules with higher clustering scores were considered the most significant biologically functional modules in interfamily grafting. All filtering parameters, including a degree threshold of 2, a node score threshold of 0.2, a k-core threshold equal to 2, and a maximum depth of 100. Hence, a threshold score of ≥ 5 was set to select significant modules and visualized with Cytoscape (v3.10.2). The second module identifier used was GLay, a comprehensive algorithm noted for identifying stable community structures within large networks by iteratively removing the highest betweenness edges within communities. GO term enrichment analyses were performed by GOATOOLS at *p* < 0.05 for each interfamily grafting-related gene module identified by both MCODE and GLay network clustering algorithms to highlight the significantly enriched GO biological processes. For the enrichment, only communities with at least 10 nodes were evaluated. In each module, the Degree algorithm of the CytoHubba plugin was used to identify hub genes and a subnetwork of hub genes in Cytoscape software (v3.10.2).

## Additional Files

**Supplementary Figure S1.** Expression levels of previously identified grafting-responsive genes. The expression levels of these genes were established following transcriptomic analysis of the intact *Nb* and *N. benthamiana*/*A. thaliana* grafted samples. TPM, transcripts per million. Error bars denote SD.

**Supplementary Dataset S1:**

**Sheet 1.** Annotation and TPM values of *N. benthamiana* genes in expression pattern A.

**Sheet 2.** Annotation and TPM values of *N. benthamiana* genes in expression pattern B.

**Sheet 3.** Annotation and TPM values of *N. benthamiana* genes in expression pattern C.

**Sheet 4.** Annotation and TPM values of *N. benthamiana* genes in expression pattern D.

**Sheet 5.** Annotation and TPM values of *N. benthamiana* genes in expression pattern E.

**Sheet 6.** Annotation and TPM values of *N. benthamiana* genes in expression pattern F.

**Supplementary Dataset S2:**

**Sheet 7.** Biological Process Gene ontology analysis for *N. benthamiana* genes in expression pattern A.

**Sheet 8.** Biological Process Gene ontology analysis for *N. benthamiana* genes in expression pattern B.

**Sheet 9.** Biological Process Gene ontology analysis for *N. benthamiana* genes in expression pattern C.

**Sheet 10.** Biological Process Gene ontology analysis for *N. benthamiana* genes in expression pattern D.

**Sheet 11.** Biological Process Gene ontology analysis for *N. benthamiana* genes in expression pattern E.

**Sheet 12.** Biological Process Gene ontology analysis for *N. benthamiana* genes in expression pattern F.

**Supplementary Dataset S3:**

**Sheet 13.** Bayesian network data for *N. benthamiana* genes in expression pattern A.

**Sheet 14.** Bayesian network data for *N. benthamiana* genes in expression pattern B.

**Sheet 15.** Bayesian network data for *N. benthamiana* genes in expression pattern C.

**Sheet 16.** Bayesian network data for *N. benthamiana* genes in expression pattern D.

**Sheet 17.** Bayesian network data for *N. benthamiana* genes in expression pattern E.

**Sheet 18.** Bayesian network data for *N. benthamiana* genes in expression pattern F.

**Supplementary Dataset S4:**

**Sheet 19.** Details of the hub genes identified by the CytoHubba algorithm using the Degree Method in the core Bayesian network of the expression pattern A.

**Sheet 20.** Details of the hub genes identified by the CytoHubba algorithm using the Degree Method in the core Bayesian network of the expression pattern B.

**Sheet 21.** Details of the hub genes identified by the CytoHubba algorithm using the Degree Method in the core Bayesian network of the expression pattern C.

**Sheet 22.** Details of the hub genes identified by the CytoHubba algorithm using the Degree Method in the core Bayesian network of the expression pattern D.

**Sheet 23.** Details of the hub genes identified by the CytoHubba algorithm using the Degree Method in the core Bayesian network of the expression pattern E.

**Sheet 24.** Details of the hub genes identified by the CytoHubba algorithm using the Degree Method in the core Bayesian network of the expression pattern F.

**Supplementary Dataset S5:**

**Sheet 25.** Details of genes and Biological Process Gene Ontology categories significantly over-represented in the subnetworks of the MCODE modules of the expression pattern A.

**Sheet 26.** Details of genes and Biological Process Gene Ontology categories significantly over-represented in the subnetworks of the MCODE modules of the expression pattern B.

**Sheet 27.** Details of genes and Biological Process Gene Ontology categories significantly over-represented in the subnetworks of the MCODE modules of the expression pattern C.

**Sheet 28.** Details of genes and Biological Process Gene Ontology categories significantly over-represented in the subnetworks of the MCODE modules of the expression pattern D.

**Sheet 29.** Details of genes and Biological Process Gene Ontology categories significantly overrepresented in the subnetworks of the MCODE modules of the expression pattern E.

**Sheet 30.** Details of genes and Biological Process Gene Ontology categories significantly over-represented in the subnetworks of the MCODE modules of the expression pattern F.

**Supplementary Dataset S6:**

**Sheet 31.** Details of genes and Biological Process Gene Ontology categories significantly over-represented in the subnetworks of the GLay modules of the expression pattern A.

**Sheet 32.** Details of genes and Biological Process Gene Ontology categories significantly over-represented in the subnetworks of the GLay modules of the expression pattern B.

**Sheet 33.** Details of genes and Biological Process Gene Ontology categories significantly over-represented in the subnetworks of the GLay modules of the expression pattern C.

**Sheet 34.** Details of genes and Biological Process Gene Ontology categories significantly over-represented in the subnetworks of the GLay modules of the expression pattern D.

**Sheet 35.** Details of genes and Biological Process Gene Ontology categories significantly over-represented in the subnetworks of the GLay modules of the expression pattern E.

**Sheet 36.** Details of genes and Biological Process Gene Ontology categories significantly over-represented in the subnetworks of the GLay modules of the expression pattern F.

**Supplementary Dataset S7:**

**Sheet 37.** Details of the hub genes identified by the CytoHubba algorithm using the Degree Method in the subnetworks of the GLay modules of the expression pattern A.

**Sheet 38.** Details of the hub genes identified by the CytoHubba algorithm using the Degree Method in the subnetworks of the GLay modules of the expression pattern B.

**Sheet 39.** Details of the hub genes identified by the CytoHubba algorithm using the Degree Method in the subnetworks of the GLay modules of the expression pattern C.

**Sheet 40.** Details of the hub genes identified by the CytoHubba algorithm using the Degree Method in the subnetworks of the GLay modules of the expression pattern D.

**Sheet 41.** Details of the hub genes identified by the CytoHubba algorithm using the Degree Method in the subnetworks of the GLay modules of the expression pattern E.

**Sheet 42.** Details of the hub genes identified by the CytoHubba algorithm using the Degree Method in the subnetworks of the GLay modules of the expression pattern F.

**Supplementary Dataset S8:**

**Sheet 43.** Annotation of selected upregulated genes during grafting in expression pattern A.

**Sheet 44.** Annotation of selected upregulated genes during grafting in expression pattern B.

**Sheet 45.** Annotation of selected upregulated genes during grafting in expression pattern C.

**Sheet 46.** Annotation of selected downregulated genes during grafting in expression pattern D.

**Sheet 47.** Annotation of selected downregulated genes during grafting in expression pattern E.

**Sheet 48.** Annotation of selected downregulated genes during grafting in expression pattern F.

**Supplementary Text S1.** Gene profiles, functional modules, and hub genes of expression patterns A–F.

## Author Contributions

MN and KK conceived and designed the study. FO and KK performed the formal analysis and prepared figures and/or tables. MN and KK provided funding and project supervision. FO wrote the original draft manuscript. All authors reviewed & edited the manuscript. All authors read and approved the final manuscript for publication.

## Data Availability

All data generated or analyzed during this study are included in this published article and additional files. All the raw datasets used in this manuscript are available from the DNA Data Bank of Japan (DDBJ; http://www.ddbj.nig.ac.jp/) under accession numbers DRA009936 and DRA017203.

