## Supplemental Files (PDF and Excel) for "A network analysis for the identification of gene modules in the transcriptome during *Nicotiana benthamiana* interfamily grafting": OpokuAgyemang_et_al_Supplementary_Text_1 - bioRxiv.pdf

Here, we describe the gene profiles, functional modules, and hub genes of each gene expression pattern. These patterns A–F reflect the transcriptional dynamics of gene sets during the grafting process. Patterns A–C, with the upregulated gene sets at defined stages, are potentially associated with key graft formation processes. In contrast, patterns D–F, with gene sets exhibiting reduced gene expression during graft healing, may be repressed to facilitate the strategic reallocation of cellular resources in favor of healing processes. Pattern A genes had sustained high expression, indicating continuous roles as key factors throughout the entire period for the initiation and maintenance of graft union formation. Pattern B genes were temporally upregulated specifically during the early stage, suggesting their potentially specialized involvement in immediate response during grafting, but rapidly declined for subsequent processes. Pattern C genes were predominantly upregulated during the late periods, and these gene sets may be involved in the long-term stability and function of the graft. Among the repressed patterns D–F, pattern D is unique, with genes specifically upregulated at the early stage but downregulated at the late stage. Transient expression of pattern D genes indicates their potential roles in the early response to grafting, but must be actively repressed later to allow for the proper function of graft-related genes in the later stages. Pattern E genes are downregulated specifically at the early stage, suggesting they may be antagonistic to the immediate wound response, and their suppression is required to enable the expression of early graft-healing genes similar to the pattern F genes repressed throughout the entire period.

### Pattern A

#### *Gene profiles of pattern A.*

Pattern A shows genes that, compared to the starting point (intact), were upregulated from 1 to 7 DAG and continuously upregulated at 14 to 28 DAG after grafting. It had the highest number of genes (1,136 genes) out of the six patterns. These genes largely belong to major functional gene classes related to transcriptional regulation, signal transduction (particularly protein kinases), disease resistance, RNA processing, transport, protein modification and degradation, and metabolism. This indicates grafting triggers significant changes in gene regulation, cellular signaling, and metabolic processes.

In the Bayesian gene regulatory network of pattern A, we identified some graft-related genes with known functional roles in plant grafting, including cell wall modification/reconstruction, vascular development, wound and stress response, plasmodesmata and intercellular communication, auxin and brassinosteroid signaling, ethylene response, cell cycle and division, sugar transport, and gene regulation. These genes related to such functional processes during grafting include genes encoding NAC domain-containing proteins (*NAC007*, *NAC060*, *NAC073*, *NAC53*, *NAC036*, *NAC04*, *NAC02*, *VND1* and *VND7*), LOB domain-containing proteins (*LBD4*), homeobox proteins (*HAT14* and *KNAT7*), ethylene-responsive factors (*ERF1* and *ERF106*), auxin-related proteins (*PIN3*, *PIN7*, *FL8*, *IAA17* and *DFL1*), glycosyl hydrolases (*β-1,3-glucanase* and *ENGase85B*), pectin methylesterase inhibitors (*PMEI* and *PMEI13*), wound-responsive proteins (*WIP3* and *BBD2*), plasmodesmata proteins (*PDLP2*), sugar transporter (*STP14*) and brassinosteroid-related proteins (*BRL1*) (Supplementary Dataset S8: Sheet 43).

Moreover, the hub genes identified in the Bayesian gene regulatory network of pattern A using the degree algorithm of the CytoHubba plugin, including *Nbe.v1.1.chr01g00700* (a homolog gene encoding NB-ARC domain-containing disease resistance protein) is involved in pathogen recognition and activation of immune responses [166], *Nbe.v1.1.chr02g05740* (a homolog of *WRKY6*) is involved in plant pathogen defense [45], and *Nbe.v1.1.chr06g24720* (a homolog of *RAF36*), a novel plant susceptibility factor that negatively regulates plant resistance to pathogens [46], highlights the complex regulatory network involved in immunity and defense regulation during grafting. These genes have established roles in plant defense and immune responses and may be promising candidates to unravel immunity mechanisms during grafting.

In the Bayesian gene regulatory network of pattern A, 5 significant modules (threshold score > 5) were identified using the MCODE algorithm, and 4 modules were identified using the GLay community clustering algorithm. MCODE module 1 is mainly enriched in cell wall polysaccharide biosynthesis, metabolism, and biogenesis-related biological processes. MCODE module 2 is mainly enriched in nuclear protein localization and organization-related biological processes. MCODE module 3 is mainly enriched in the regulation of secondary metabolism, biosynthesis, and amino acid transport-related biological processes. MCODE module 4 is mainly enriched in cell wall glycoprotein biosynthesis, carbohydrate metabolism, and chemical stress responses related to biological processes. MCODE module 5 is mainly enriched in the regulation of gene expression, steroid receptor signaling, and protein homeostasis-related biological processes. GLay module 1 is mainly enriched in cell wall biogenesis, host-symbiont immune dynamics, and defense responses related to biological processes. GLay module 2 is mainly enriched in nucleic acid metabolism, recombination, and DNA repair-related biological processes. GLay module 3 is mainly enriched in amino acid metabolism and catabolism-related biological processes. GLay module 4 is mainly enriched in chromosome organization and epigenetic regulation-related biological processes. These complementary approaches allow for the identification of functional graft-related modules within the network of expression pattern A, from small, tightly connected modules with strong node interactions using MCODE to larger, loosely connected, hierarchically organized communities covering multiple biological functions using GLay.

The GO superclusters identified through REVIGO treemap analysis for pattern A represented broad biological themes including "DNA recombination", "DNA damage response", "regulation of response to water deprivation", "disruption of cellular anatomical structure in another organism" "xenobiotic transmembrane transport", "telomerase RNA stabilization", "regulation of secondary cell wall biogenesis" and "xylem vessel member cell differentiation", while the GO terms enriched in the MCODE and GLay modules provide a more detailed view of gene function within specific network structures identified. The MCODE modules identified were independent with no interconnections, likely representing distinct functional pathways. However, their enriched GO terms still align with the broader GO superclusters identified to be enriched by the genes in pattern A and presented in the REVIGO treemap, highlighting conserved functional networks that underpin key biological processes. For example, MCODE module 1 in pattern A enriched in "regulation of secondary cell wall biogenesis" is associated with the "regulation of secondary cell wall biogenesis"

GO supercluster in the REVIGO treemap. MCODE 3 in pattern A enriched in "regulation of cell killing", "response to external stimulus" and "xenobiotic transmembrane transport" may appear independent in the network clustering but, all contribute to the "DNA damage response", "regulation of response to water deprivation", "xenobiotic transmembrane transport" and "disruption of cellular anatomical structure in another organism" GO superclusters in the REVIGO treemap. This reveals the contribution of distinct molecular functions to higher-order biological processes. Similarly, MCODE module 4 in pattern A enriched in "alpha−amino acid catabolic process", "cell wall glycoprotein biosynthetic process", "cell wall mannoprotein biosynthetic process", "DNA replication checkpoint signaling" and "telomerase RNA stabilization" are associated with "DNA recombination", "DNA damage response", "regulation of response to water deprivation" and "telomerase RNA stabilization" GO superclusters in the REVIGO treemap. MCODE module 5 in pattern A enriched in the "trehalose catabolic process" is associated with the "DNA recombination" GO supercluster in the REVIGO treemap.

The GLay modules identified exhibited connections among a few genes, suggesting a potential functional linkage between the biological processes identified in each GLay module, and these processes can be matched to the organized biological themes captured in the GO superclusters. For example, GLay module 1 in pattern A enriched in "xylem vessel member cell differentiation", "xenobiotic transmembrane transport", "regulation of secondary cell wall biogenesis", "lactone biosynthetic process", "regulation of cell killing", "regulation of response to water deprivation", "response to external stimulus", and "effector-mediated modulation of host process by symbiont" contributes to broader GO superclusters related to "regulation of secondary cell wall biogenesis", "xylem vessel member differentiation", "DNA recombination", "xenobiotic transmembrane transport" and "disruption of cellular anatomical structure in another organism". GLay module 2 in pattern A enriched in "DNA recombination", "DNA metabolic process", "trehalose catabolic process", "DNA damage response", "positive regulation of DNA metabolic process" and "telomerase RNA stabilization" contributes to the GO superclusters "DNA recombination", "DNA damage response", "regulation of response to water deprivation" and "telomerase RNA stabilization". GLay module 3 in pattern A enriched in "alpha−amino acid catabolic process", "arginine catabolic process", "small molecule catabolic process", and "cellular response to stimulus" contributes to the GO superclusters "DNA recombination" and "DNA damage response".

***Functional enrichments and hub genes of MCODE modules in pattern A.***

MCODE module 1 in pattern A is related to cell wall biogenesis and organization, polysaccharide metabolism, energy metabolism, and cytoskeleton organization. These processes may enhance successful graft union formation by promoting cell adhesion, tissue regeneration, and vascular reconnection between the scion and rootstock. For this module, we identified hub genes with established roles in secondary cell wall formation (*Nbe.v1.1.chr04g00990* and *Nbe.v1.1.chr15g02820*, homologs of *NAC073*), secondary cell wall lignification (*Nbe.v1.1.chr06g29150*, a homolog of *ATL54*) and cell wall component synthesis (*Nbe.v1.1.chr15g35860*, a homolog of *PGSIP3*), indicating their potential function in grafting (Supplementary Dataset S5: Sheet 25). *Nbe.v1.1.chr06g29150* (a homolog of

*ATL54*) encodes a ubiquitin ligase gene that has been identified to be expressed in fiber cells and vessel elements of secondary cell walls. This gene is known to be directly regulated by *MYB46*, which is a key transcriptional activator of secondary cell wall formation [167], and so *ATL54* is probably involved in the secondary wall formation process of grafting. *Nbe.v1.1.chr04g00990* and *Nbe.v1.1.chr15g02820* (homologs of *NAC073*) are associated with secondary cell wall specificity, and suppression of its expression levels results in slow thickening of secondary cell walls [168,169]. *NAC073* is involved in lignifying secondary cell walls [170], and so *NAC073* is probably involved in the lignification and secondary cell wall thickening process of grafting. *Nbe.v1.1.chr15g35860* (a homolog of *PGSIP3*) encodes a glycosyl transferase involved in glucuronoxylan biosynthesis and hemicellulose synthesis [171,172]. These are the principal hemicelluloses in secondary cell walls [173], and so *PGSIP3* is probably involved in the synthesis of cell wall components during grafting.

MCODE module 2 in pattern A is related to protein localization to the nuclear inner membrane and nuclear envelope, nuclear pore organization, and NLS-bearing protein import into the nucleus. These processes may regulate stress responses, gene expression, signaling, and cell cycle during graft union formation. We identified hub genes involved in intracellular calmodulin-binding activity (*Nbe.v1.1.chr08g30190*, a homolog gene encoding RNA ligase/cyclic nucleotide phosphodiesterase family protein), and cysteine-type deubiquitinase activity (*Nbe.v1.1.chr04g32960*, a homolog of *OTU5*) (Supplementary Dataset S5: Sheet 25). *Nbe.v1.1.chr08g30190* (a homolog gene encoding RNA ligase/cyclic nucleotide phosphodiesterase family protein) belongs to the 2H phosphoesterase superfamily. This family of enzymes is involved in various RNA metabolic processes, including RNA processing and degradation [174,175], and so, it may probably be involved in the regulation of gene expression and cellular responses during grafting, which will specifically enhance stress response and cellular communication. *Nbe.v1.1.chr04g32960* (a homolog of *OTU5*) encodes an ovarian tumor domain (OTU)-containing a deubiquitinating enzyme family protein involved in cysteine-type deubiquitinase activity. The deubiquitination activity of the genes ensures the stability of target proteins through the removal of ubiquitin, and regulates genes associated with root development [176], root responses to phosphate starvation [177], and activates the major flowering repressors *FLC*, *MAF4*, and *MAF5* [178]. This gene may modulate the expression, stability, and activity of genes crucial for grafting processes such as nutritional stress responses, gene expression, and signaling.

MCODE module 3 in pattern A is related to metabolic processes (regulation of secondary metabolite and carbohydrate biosynthesis, sulfur and glucose metabolism), transport processes (aromatic amino acid and p-coumaryl alcohol transport), defense and stress responses (external and extracellular stimulus, nutrient levels) and cell death (modulation of host programmed cell death, symbiont of host resistance gene-dependent defense response and positive regulation of killing of cells of another organism). These processes may be crucial for graft healing, vascular reconnection, and compatibility between scion and rootstock, which may enhance successful graft union formation. In this module, hub genes involved in immune response and cell death (*Nbe.v1.1.chr01g00700*, a homolog gene encoding NB-ARC domain-containing disease resistance protein), selective autophagy (*Nbe.v1.1.chr03g13320*, a homolog of *C53*), and mechanically activated ion channel gene (*Nbe.v1.1.chr02g03460*, a homolog of *OSCA2.3*) were identified (Supplementary Dataset S5:

Sheet 25). *Nbe.v1.1.chr01g00700* (a homolog gene encoding NB-ARC domain-containing disease resistance protein) contains an NB-ARC domain, a nucleotide-binding adaptor shared by APAF-1, certain R gene products, and CED-4, and is typically found in disease resistance proteins [179]. The nucleotide-binding site of NB-ARC has ARC1 (Apoptotic protease activation factor 1, APAF-1, and cell death protein 4, CED4) and ARC2 regions that activate downstream signaling targets [180,181]. This gene may play a crucial role in priming the immune response and disease resistance processes to prepare for potential risks of pathogen attacks during grafting. *Nbe.v1.1.chr03g13320* (a homolog of *C53*) has been shown to encode a conserved reticulophagy receptor associated with selective autophagy and ribosome stalling at the endoplasmic reticulum (ER) [182]. *C53* is crucial for ER homeostasis and stress tolerance in plants; hence, during grafting, it may enhance ER-related stress response and maintain cellular homeostasis. *Nbe.v1.1.chr02g03460* (a homolog of *OSCA2.3*) encodes a hyperosmolality-gated calcium-permeable channel family [183] involved in osmotic adjustments [183,184], with differential roles in stress response and tolerance [185,186], and so *OSCA2.3* is probably involved in osmo-sensing and stress response processes during grafting. This module's hub genes involved in immunity and selective autophagy are candidates to study underlying grafting mechanisms, such as coordinated defense and cellular maintenance.

MCODE module 4 in pattern A is related to autophagy, cell wall processes, and response to stimuli. The hub genes identified were involved in DNA damage repair and homologous recombination (*Nbe.v1.1.chr11g18850*, a homolog of *RAD9*), cell division, pollen and seed development (*Nbe.v1.1.chr03g08880*, a homolog of *MDKIN1*), and cell proliferation and differentiation (*Nbe.v1.1.chr18g38880*, a homolog of *PAS1*) (Supplementary Dataset S5: Sheet 25). The respective roles of the hub genes screened in this module suggest that they are potential genes in maintaining genome stability, cell division, and tissue regeneration during grafting. *Nbe.v1.1.chr11g18850* (a homolog of *RAD9*) encodes a key component of the DNA damage response pathway and genotoxic stress response in plants. This gene is crucial for maintaining genomic integrity by participating in cell-cycle checkpoint control and DNA repair [187,188], and so *RAD9* probably enhances the DNA damage repair caused by the physical wounding during grafting. Also, during plant grafting, *RAD9* may support the response to stresses by ensuring genomic stability and proper cell-cycle regulation. *Nbe.v1.1.chr03g08880* (a homolog of *MDKIN1*) encodes a malectin domain kinesin, which is a type of motor protein shown to be associated with cell division throughout the plant, pollen, and seed development [189]. So, *MDKIN1* is probably involved in cell division and tissue regeneration during grafting. *Nbe.v1.1.chr18g38880* (a homolog of *PAS1*) encodes an FK506-binding protein, an immunophilin, involved in the hormonal regulation of cell proliferation and differentiation during plant development [190–192]. *PAS1* regulates and balances the effect of key hormones, including cytokinin and auxin, for coordinated cell division and differentiation [191], and so *PAS1* is probably involved in the regulation of hormonal signaling and ensuring proper cell proliferation and tissue differentiation process during grafting.

MCODE module 5 in pattern A is related to gene expression, protein processing, cell cycle regulation, growth, transport, and structural modifications. Following network analysis, we identified hub genes with established roles in stress response and ROS scavenging

(*Nbe.v1.1.chr19g16560*, a homolog of *GSTU8*), pathogen resistance and stress response (*Nbe.v1.1.chr06g24720*, a homolog of *RAF36*), immune response (*Nbe.v1.1.chr16g33430*, a homolog of *RFO3*) and meristem activity (*Nbe.v1.1.chr15g15530*, a homolog of *BRL1*) (Supplementary Dataset S5: Sheet 25). *Nbe.v1.1.chr19g16560* (a homolog of *GSTU8*) encodes a glutathione S-transferase (GST), which is part of a large family of enzymes involved in detoxification and response to abiotic stresses, including drought, salt, and oxidative stress, by scavenging ROS and protecting cells from oxidative damage [193]. The role of the *GSTU8* genes suggests that they are probably involved in improving stress response and reducing oxidative damage during grafting. *Nbe.v1.1.chr06g24720* (a homolog of *RAF36*) encodes a Raf-like kinase that plays a role in negatively regulating plant resistance to pathogens and stress signaling and hormonal responses [93]. The role of the gene in pathogen resistance, stress signaling, and ABA response regulation may be crucial for wound healing and stress mitigation during grafting. *Nbe.v1.1.chr16g33430* (a homolog of *RFO3*) encodes a receptor-like kinase (RLK), a disease-resistance protein that plays a significant role in plant defense mechanisms and confers resistance to a broad spectrum of pathogens, including wilt fungus *Fusarium oxysporum*. This gene functions as a pattern-recognition receptor (PRR) that helps detect pathogen-associated molecular patterns (PAMPs) and activate defense responses. *RFO3* is highly expressed in vascular tissues, which are critical sites for pathogen entry and spread [194,195]. Hence, *RFO3* is probably involved in wound healing at the graft junction by activating defense mechanisms against potential pathogens. *Nbe.v1.1.chr15g15530* (a homolog of *BRL1*) encodes a leucine-rich repeat receptor-like kinase (LRR-RLK) involved in brassinosteroid (BR) signaling [196] and plays a specific role in vascular differentiation and maintaining proper xylem and phloem ratios [197,198]. This is essential for the development of functional vascular tissues, including protophloem differentiation in root meristems [197,198]. Hence, *BRL1* is probably involved in the formation of new vascular tissues, cell elongation, and differentiation processes, further supporting the role of BRs in promoting xylem formation [110] at the graft junction during grafting. During grafting, the suggested coordinated function of these genes in mitigating oxidative damage to promote cell viability, defense mechanisms, and regenerative growth at the wound site suggests their potential as promising candidates for grafting-related studies.

In addition to the five significant (threshold score > 5) MCODE modules in the network of pattern A discussed above, the remaining 10 modules, while not meeting the threshold, also exhibit valuable insights through their node counts and functional enrichments. By exploring these aspects, we can gain a comprehensive understanding of their roles and contributions within the broader network. Below, we briefly summarize their number of nodes and key features.

The seed node of module 6 (59 nodes and 139 edges) was *Nbe.v1.1.chr12g27210* (a homolog of *LBD4*). This module is mainly enriched in the GO-BP terms, "apical protein localization", "regulation of nucleotide catabolic process", and "regulation of cutin biosynthetic process". Hub genes identified in module 6 were *Nbe.v1.1.chr06g20570* (a homolog of translocon at the outer envelope membrane of chloroplasts 159 gene, *TOC159*), *Nbe.v1.1.chr13g00825* (a homolog of *FL8*) and *Nbe.v1.1.chr05g04450* (a homolog of embryo defective 506 gene, *EMB506*) (Supplementary Dataset S5: Sheet 25).

The seed node of module 7 (78 nodes and 177 edges) was *Nbe.v1.1.chr16g09470* (a homolog gene encoding sec14p-like phosphatidylinositol transfer family protein). This module is mainly enriched in the GO-BP terms, “protein lipoylation”, “modulation of the process of another organism”, “transition metal ion transport”, and “zinc ion transmembrane transport”. Hub genes identified in module 7 were *Nbe.v1.1.chr01g25200* (a homolog of sticky generative cell gene, *SGC*), *Nbe.v1.1.chr01g38660* (a homolog gene encoding C2H2-like zinc finger protein, *ZAT9*), *Nbe.v1.1.chr13g03410* (a homolog gene encoding leucine-rich repeat family protein) and *Nbe.v1.1.chr16g24630* (a homolog of the needed for RDR2-independent DNA methylation gene, *NERD*) (Supplementary Dataset S5: Sheet 25).

The seed node of module 8 (101 nodes and 214 edges) was *Nbe.v1.1.chr02g24160* (a homolog gene encoding homeobox protein, *HAT9*). This module is mainly enriched in the GO-BP terms, “regulation of actin filament bundle assembly”, “mitochondrial protein catabolic process”, and “regulation of JUN kinase activity”. Hub genes identified in module 8 were *Nbe.v1.1.chr02g29950* (a homolog of *PEX6*), *Nbe.v1.1.chr07g23050* (a homolog of glyceraldehyde-3-phosphate dehydrogenase of plastid 1 gene, *GAPCP-1*) and *Nbe.v1.1.chr13g34440* (a homolog gene encoding KH domain-containing protein, *KH7*) (Supplementary Dataset S5: Sheet 25).

The seed node of module 9 (9 nodes and 17 edges) was *Nbe.v1.1.chr02g24160* (a homolog of root hair specific 2 gene, *RHS2*). This module had no enriched GO-BP terms, due to the small number of genes. The hub gene identified in module 9 was *Nbe.v1.1.chr14g00990* (a homolog of salt tolerance zinc-finger gene, *STZ*) (Supplementary Dataset S5: Sheet 25).

The seed node of module 10 (96 nodes and 200 edges) was *Nbe.v1.1.chr11g26950* (a homolog gene encoding RmlC-like cupins superfamily protein). This module is mainly enriched in the GO-BP terms, “telomere capping”, “xylem vessel member cell differentiation”, and “positive regulation of secondary cell wall biogenesis”. Hub genes identified in module 10 were *Nbe.v1.1.chr19g07140* (a homolog of *RIMA*), *Nbe.v1.1.chr19g07140* (a homolog of *STZ*), *Nbe.v1.1.chr07g05500* (a homolog of constitutive expresser of PR genes 1, *CPR1*), *Nbe.v1.1.chr13g18840* (a homolog gene encoding urease accessory protein F, *UREF*) and *Nbe.v1.1.chr16g25250* (a homolog of *ARI8*) (Supplementary Dataset S5: Sheet 25).

The seed node of module 11 (44 nodes and 84 edges) was *Nbe.v1.1.chr11g30130* (a homolog of PWWP-domain interactor of polycombs1 gene, *PWO1*). This module is mainly enriched in the GO-BP terms, “disruption of the cellular component of another organism”, “glycosaminoglycan catabolic process”, “peptidoglycan metabolic process”, and “cell wall disruption in another organism”. Hub genes identified in module 11 were *Nbe.v1.1.chr14g20700* (a homolog gene encoding 2-oxoglutarate (2OG) and Fe(II)-dependent oxygenase superfamily protein) and *Nbe.v1.1.chr01g17980* (a homolog gene encoding DNA-binding bromodomain-containing protein) (Supplementary Dataset S5: Sheet 25).

The seed node of module 12 (56 nodes and 99 edges) was *Nbe.v1.1.chr17g51700* (a homolog gene encoding transcription elongation factor (TFIIS) family protein). This module is mainly enriched in the GO-BP terms, “DNA-templated transcription elongation”, “putrescine biosynthetic process from arginine, using agmatinase”, “positive regulation of nitrogen utilization", and "photosynthesis, light harvesting". Hub genes identified in module 12 were *Nbe.v1.1.chr09g22760* (a homolog gene encoding transducin/WD40 repeat-like superfamily protein), *Nbe.v1.1.chr19g35640* (a homolog of lsd one like 2 gene, *LOL2*), *Nbe.v1.1.chr01g25350* (a homolog gene encoding zinc-finger C-x8-C-x5-C-x3-H type family protein, *ATC3H5*) and *Nbe.v1.1.chr05g25750* (a homolog of forms aploid and binucleate cells 1b gene, *FAB1B*) (Supplementary Dataset S5: Sheet 25).

The seed node of module 13 (6 nodes and 9 edges) was *Nbe.v1.1.chr01g21470* (a homolog gene encoding P53/DNA damage-regulated protein, *PDRG1*). This module had no enriched GO-BP terms, due to the small number of genes. The hub gene identified in module 13 was *Nbe.v1.1.chr14g00990* (a homolog gene encoding mitochondrial transcription termination factor family protein, *MTERF7*) (Supplementary Dataset S5: Sheet 25). The seed node of module 14 (with 54 nodes and 93 edges) was *Nbe.v1.1.chr11g39030* (a homolog of RECQ helicase l1 gene, *RECQl1*). This module is mainly enriched in the GO-BP terms, "cellular response to water deprivation", "cellular response to salt", and "signal transduction in response to DNA damage". Hub genes identified in module 14 were *Nbe.v1.1.chr18g41050* (a homolog gene encoding GINS complex protein), *Nbe.v1.1.chr04g35210* (a homolog of mizu-kussei 1 gene, *MIZ1*), *Nbe.v1.1.chr02g01500* (a homolog gene encoding transducin/WD40 repeat-like superfamily protein) and *Nbe.v1.1.chr15g07140* (a homolog gene encoding RING/FYVE/PHD zinc finger superfamily protein) (Supplementary Dataset S5: Sheet 25).

The seed node of module 14 (54 nodes and 93 edges) was *Nbe.v1.1.chr11g39030* (a homolog of RECQ helicase l1 gene, *RECQl1*). This module is mainly enriched in the GO-BP terms, "cellular response to water deprivation", "cellular response to salt", and "signal transduction in response to DNA damage". Hub genes identified in module 14 were *Nbe.v1.1.chr18g41050* (a homolog gene encoding GINS complex protein), *Nbe.v1.1.chr04g35210* (a homolog of mizu-kussei 1 gene, *MIZ1*), *Nbe.v1.1.chr02g01500* (a homolog gene encoding transducin/WD40 repeat-like superfamily protein) and *Nbe.v1.1.chr15g07140* (a homolog gene encoding RING/FYVE/PHD zinc finger superfamily protein) (Supplementary Dataset S5: Sheet 25).

The seed node of module 15 (22 nodes and 31 edges) was *Nbe.v1.1.chr03g22120* (a homolog gene encoding protein kinase superfamily protein). This module is mainly enriched in the GO-BP terms, "positive regulation of mitochondrial translation", "regulation of actin cytoskeleton reorganization", and "cellular response to estrogen stimulus". The hub gene identified in module 15 was *Nbe.v1.1.chr04g20410* (a homolog gene encoding F-box/RNI-like superfamily protein) (Supplementary Dataset S5: Sheet 25).

***Functional enrichments and hub genes of GLay modules in pattern A.***
GLay module 1 in pattern A is enriched in cell wall-related processes (secondary cell wall biogenesis, xylem vessel member cell differentiation, xylem development and tracheary element differentiation), cell death and defense (regulation of cell killing and positive regulation of killing of cells of another organism), water stress responses (Regulation of response to water deprivation), metabolic processes (strigolactone metabolic process, phosphorus metabolic process and lactone metabolic process) and host-symbiont interactions (modulation by symbiont of host programmed cell death, symbiont of host resistance gene-

dependent defense response and induction by symbiont of host programmed cell death). In this module, we identified hub genes involved in cambial regulation and meristematic activity-related (*Nbe.v1.1.chr02g13470*, a homolog of *LBD4*), stress tolerance and pathogen defense (*Nbe.v1.1.chr19g35510*, a homolog of *LECRK-VII.1* and *Nbe.v1.1.chr04g32570*, a homolog gene encoding heat shock protein 70, *HSP70*), sugar transport protein (*Nbe.v1.1.chr15g24130*, a homolog of *STP14*) (Supplementary Dataset S7: Sheet 37). *Nbe.v1.1.chr02g13470* (a homolog of *LBD4*) is a previously known gene that is related to meristematic activity processes during grafting [7]. This gene, following a synthesized molecular-informed model for the developmental progression of junction formation, was predicted as a hub gene at later developmental stages (3 and 5 DAG) during pepper-pepper grafting. It was identified to be involved in meristematic activity and a key regulator of two xyloglucan endotransglucosylase/hydrolase (*XTH*) genes (*XTH22* and *XTH38*) [7]. In the current GLay module 1, *LBD4* was identified to regulate AUX/IAA transcriptional regulator family protein (*IAA17* and *AXR3*) involved in auxin signaling and cell elongation [199] ethylene-responsive transcription factor (*ERF106*) involved in ethylene-activated signaling and identified to regulate *PcXTH17*, *SrXTH17*, and *SrXTH25* during *Populus cathayana* and *Salix rehderiana* grafting [200], rhamnogalacturonate lyase family protein (*RGIL6* and *RGL1*) involved in cell-cell adhesion and pectin composition [201,202], and *β-1,3-endoglucanase* involved in defense response [203,204]. For the remaining hub genes, *Nbe.v1.1.chr19g35510* (a homolog of *LECRK-VII.1*) encodes a member of the LecRK family, which contains a conserved lectin domain and a kinase domain involved in the regulation of jasmonic acid (JA) synthesis and response by affecting the activity of lipoxygenase (LOX) [50]. *LecRK-VII.1* has been identified for salt stress response and affects the expression of the $Ca^{2+}$ sensors, including annexin (ANN) [50]. Hence, *LECRK-VII.1* is probably involved in the stress response processes of grafting. *Nbe.v1.1.chr04g32570* (a homolog of *HSP70*) has been identified to be upregulated in response to biotic stressors and is a crucial component of plant immunity [205]. The gene is known to be involved in PAMP (pathogen-associated molecular pattern)-triggered immunity (PTI) and effector-triggered immunity (ETI) responses [205,206]. Therefore, *HSP70* is probably involved in immunity response processes of grafting. *Nbe.v1.1.chr15g24130* (a homolog of *STP14*) is a sugar transporter that encodes a protein involved in the active transport of galactose [207–209]. This has shown high expression levels in both source and sink tissues, and its expression is regulated by cell wall degradation-induced factors [208] such as cell wall-degrading β-galactosidases (*MUM2* and *Gal2*) involved in cell wall degradation [210,211]. *STP14* is crucial for recycling cell wall-derived galactose during developmental processes [208], and so *STP14* is probably involved in sugar transport, cell wall sugar recycling, and the cell wall degradation process of grafting.

GLay module 2 in pattern A is enriched in DNA-related processes (DNA damage response, DNA repair and DNA recombination), RNA-related processes (RNA metabolic process, RNA modification and RNA stabilization), cell cycle and division processes (cell cycle process, nuclear division, meiosis I cell cycle process and mitotic recombination), metabolic processes (nucleic acid metabolic process, macromolecule metabolic process and heterocycle metabolic process) and stress response biological processes (cellular response to stress, response to exogenous dsRNA and response to ionizing radiation). In this module, we identified hub genes involved in peroxisomal biogenesis (*Nbe.v1.s00090g29950*, a homolog of *PEX6*), cell death and defense response (*Nbe.v1.1.chr19g35640*, a homolog of *LOL2*), embryogenesis (*Nbe.v1.1.chr10g38460*, a homolog of *TPXL3*), ROS mediated ABA signaling (*Nbe.v1.1.chr16g48210*, a homolog gene encoding MAP kinase 9, *MPK9*) and RNA processing (*Nbe.v1.1.chr16g08230*, a homolog of *OTP439*) (Supplementary Dataset S7: Sheet 37). *Nbe.v1.1.chr19g35640* (a homolog of *LOL2*) encodes a protein containing a zinc finger motif, which regulates cell death and defense responses [212,213]. *LOL2* is involved in the regulation of programmed cell death and defense mechanisms against pathogens such as virulent and avirulent strains of *Pseudomonas syringae* [214,215]. Hence, *LOL2* is probably involved in defense responses and the regulation of the cell death process of grafting. *Nbe.v1.s00090g29950* (a homolog of *PEX6*) encodes a protein involved in peroxisome biogenesis and maintenance, essential organelles for plant metabolic processes [216]. *PEX6* is involved in the breakdown of fatty acids stored in vital oil bodies essential for seed germination and seedling development [217]. *PEX6* is probably involved in controlling physiological stress associated with grafting by maintaining peroxisome function and facilitating stress responses and energy supply. *Nbe.v1.1.chr10g38460* (a homolog of *TPXL3*) encodes a protein involved in microtubule organization and cell division. *TPXL3* is a primary activator of α-aurora kinases essential for cell cycle progression and binds to microtubules, with a crucial role in their organization during cell division and embryogenesis [218,219]. So, *TPXL3*'s role in microtubule organization and cell division could be crucial for the proliferation of cells at the graft junction, facilitating graft union formation. *Nbe.v1.1.chr16g48210* (a homolog of *MPK9*) is expressed in guard cells and involved in ROS-mediated ABA signaling, crucial for response to abiotic stress [220] and induces stomatal closure [221,222]. *MPK9* regulation of stomatal closure might be crucial for maintaining water balance in grafted plants, especially under stress conditions. *Nbe.v1.1.chr16g08230* (a homolog of *OTP439*) encodes a PPR repeat protein involved in the splicing of mitochondrial nad5 transcripts, essential mitochondrial function, and energy production [223]. The role of *OTP439* in maintaining mitochondrial function could support the energy demands of the grafting process. Overall, GLay module 2 integrates diverse processes, including genome maintenance and cell division, with hub genes *LOL2*, *PEX6*, *TPXL3*, *MPK9*, and *OTP439,* potentially contributing to defense activation, organelle function, and cellular regeneration during grafting.

GLay module 3 in pattern A is enriched in amino acid catabolism and metabolism (arginine catabolic and metabolic processes, methionine catabolic process, aspartate family catabolic process, proline metabolic process, and tyrosine metabolic process), protein oligomerization (protein homo-tetramerization, tetramerization and homo-oligomerization), and cellular responses, biological processes (cellular response to stimulus, cellular response to UV-A). In this module, we identified hub genes involved in protein degradation (*Nbe.v1.1.chr16g25250*, a homolog of *ARI8*), ABA signaling (*Nbe.v1.1.chr19g35510*, a homolog of *GPCR*), and cell differentiation (*Nbe.v1.1.chr19g07140*, a homolog of *RIMA*) (Supplementary Dataset S7: Sheet 37). The enriched biological processes and function of the hub genes in the GLay module 3 suggest that they may be potential key regulators for tissue regeneration and stress management for successful grafting. *Nbe.v1.1.chr16g25250* (a homolog of *ARI8*) encodes a protein involved in the ubiquitin-proteasome pathway, for the degradation of proteins tagged with ubiquitin [224]. Following the facilitation of protein degradation, *ARI8* is crucial for maintaining protein quality and regulating various cellular processes, including stress response [224]. During grafting, *ARI8* could help grafted plants manage the physiological stress associated with wound damage by regulating the degradation of stress-responsive proteins. *Nbe.v1.1.chr19g07140* (a homolog of *RIMA*) encodes a protein involved in the nuclear accumulation of IYO (MINIYO) transcriptional regulators essential for initiating auxin-irreversible cell differentiation [225]. *RIMA* is also known to be expressed in meristems and organ primordia and is required for lateral root development and callus formation [225]. Hence, *RIMA* is probably involved in auxin signaling and differentiation of cells at the graft junction, promoting the formation of new tissues and graft union formation.

GLay module 4 in pattern A is enriched in chromosome and DNA organization (chromosome organization and segregation, nuclear chromosome segregation, G2/M transition of mitotic cell cycle, and cell cycle process) and histone and protein methylation biological processes (histone lysine methylation, histone H3-K36 methylation, and histone H3-K27 methylation). In this module, we identified hub genes involved in cell division (*Nbe.v1.1.chr03g08880*, a homolog of *MDKIN1*) and DNA replication (*Nbe.v1.1.chr08g22670*, a homolog of *ORC6*) (Supplementary Dataset S7: Sheet 37). This module revealed hub genes with established roles in cell division (*MDKIN1*) and DNA replication (*ORC6*), suggesting novel regulators for cell proliferation and vascular tissue formation during grafting. *Nbe.v1.1.chr03g08880* (a homolog of *MDKIN1*) encodes a malectin domain kinesin, which is a type of motor protein associated with cell division throughout the plant, pollen, and seed development [189]. So, *MDKIN1* is probably involved in cell division and tissue regeneration and may support the formation of new vascular tissue regeneration processes of grafting. *Nbe.v1.1.chr08g22670* (a homolog of *ORC6*) encodes a key component of the origin recognition complex (ORC) protein and is involved in the initiation of DNA replication [226]. Also, *ORC6* has been reported to play a key role in chromosome segregation and cytokinesis [227]. Due to the role of *ORC6* in initiating DNA replication and cell cycle regulation, it may be involved in the proliferation of cells at the graft junction.

Following the application of MCODE and GLay to identify functional gene clusters within the Bayesian regulatory network of pattern A, several hub genes were identified in both module sets, indicating overlap and potential functional convergence. The hub genes *Nbe.v1.1.chr01g00700* (a homolog gene encoding NB-ARC domain-containing disease resistance protein), *Nbe.v1.1.chr02g05740* (a homolog of *WRKY6*), and *Nbe.v1.1.chr06g24720* (a homolog of *RAF36*) were identified in MCODE modules 3 and 5 and GLay module 1. The shared hub genes are primarily associated with plant defense, immune response, and stress adaptation, highlighting their importance in the biological processes underlying this expression pattern. Additionally, *Nbe.v1.1.chr02g03460* (a homolog of *OSCA2.3*), and *Nbe.v1.1.chr15g20950* (a homolog gene encoding prolyl oligopeptidase family protein) are present as hub genes in both MCODE module 2 and GLay module 2. These overlaps illustrate how the distinct network clustering algorithms, despite the differences in module size and network topology, consistently highlight these genes as central to the regulatory network of pattern A.

### Pattern B

#### *Gene profiles of pattern B.*

Pattern B shows genes that, compared to the starting point (intact), were upregulated from 1 to 7 DAG but were downregulated at 14 to 28 DAG after grafting. It had the third highest number of genes (228 genes) out of the six patterns. Expression Pattern B comprises diverse functional classes triggered during grafting, including transcription factors (bZIP, NAC, and WRKY), protein kinases, oxidoreductases, TPR repeat proteins, ATPases, RNA processing proteins, transporters, protein modification enzymes (RING/U-box), metabolic enzymes, and DNA metabolism proteins (ligases, repair factors). This indicates significant changes in gene regulation, cellular signaling, metabolic processes, and wound response pathways during grafting.

In the Bayesian gene regulatory network of pattern B, graft-related genes were identified with known functional roles, including cell wall modification, cell division, vascular development, lignin formation, wound and stress response. These genes related to such functional processes during grafting include genes encoding NAC domain-containing proteins (*NAC007* and *NAC060*), LOB domain-containing proteins (*LBD15*), myb domain protein 83 (*MYB83*), basic helix loop helix (bHLH) DNA-binding superfamily protein, peroxidase superfamily protein (*PRX17*), IRREGULAR XYLEM protein (*IRX15-L*), glycosyl hydrolase family protein (*BXL6*), amine oxidase 1 (*AO1*), cyclin D2;1 and cyclin A2;2 (Supplementary Dataset S8: Sheet 44). In pattern B, notable hub genes identified were *Nbe.v1.1.chr04g10170* (a homolog of *TLP-3*) and *Nbe.v1.1.chr10g32330* (a homolog of *CNGC14*). The *TLP-3* homolog is involved in the activation of plant defense pathways [48], suggesting a role in enhancing immune responses during grafting to protect plants against pathogen invasion during the critical early stages of graft union formation. The *CNGC14* homolog is known to regulate auxin-mediated growth responses and root gravitropism [49], indicating its potential involvement in promoting cell expansion, directional growth, and tissue integration during grafting. Together, these hub genes may play complementary roles during grafting, with the *TLP-3* homolog strengthening defense mechanisms while the *CNGC14* homolog facilitates auxin-driven developmental processes crucial for successful graft establishment.

In this network for pattern B, 4 significant modules (threshold score > 5) were identified using the MCODE algorithm, and 4 modules were identified using the GLay algorithm. MCODE module 1 is mainly enriched in the regulation of the mitotic cell cycle, checkpoint signaling, and phase transition-related biological processes. MCODE module 2 is mainly enriched in mitotic spindle organization and chromosome organization-related biological processes. MCODE module 3 is mainly enriched in carbohydrate metabolism-related biological processes. MCODE module 4 is mainly enriched in purine deoxyribonucleotide metabolism and biosynthesis-related biological processes. GLay module 1 is mainly enriched in RNA metabolism, transcriptional regulation, and glucose response-related biological processes. GLay module 2 is mainly enriched in the regulation of cell wall biogenesis, macromolecular transport, and cellular component organization-related biological processes. GLay module 3 is mainly enriched in the regulation of mitotic and meiotic cell cycles and chromosome organization-related biological processes. GLay module

4 is mainly enriched in the regulation of development, stress adaptation, and defense response-related biological processes.

The GO superclusters identified through REVIGO treemap analysis for pattern B included “negative regulation of mitotic cell cycle”, “nucleic acid phosphodiester bond hydrolysis”, “fucose catabolic process”, “cell cycle process”, “vesicle−mediated transport to the plasma membrane”, “fucosylation”, “sporulation”, “cell cycle” and “organic cyclic compound metabolic process”. MCODE module 1 in pattern B enriched in “negative regulation of mitotic cell cycle”, “regulation of chromosome organization”, “DNA recombination” and “deoxyribonucleoside diphosphate biosynthetic process” is associated with the “negative regulation of mitotic cell cycle”, “nucleic acid phosphodiester bond hydrolysis”, “fucose catabolic process” and “cell cycle process GO superclusters in the REVIGO treemap. MCODE module 3 revealed enriched terms including “nucleic acid phosphodiester bond hydrolysis”, “cellular nitrogen compound metabolic process”, “fucose metabolic process”, “fucosylation” associated with the GO superclusters “nucleic acid phosphodiester bond hydrolysis”, “fucose catabolic process” and “fucosylation” in the REVIGO treemap. MCODE module 4 enriched in “nucleoside monophosphate phosphorylation” is associated with the “fucose catabolic process” GO superclusters in the REVIGO treemap.

Enriched analysis of GLay module 1 in pattern B revealed biological processes including “nucleic acid phosphodiester bond hydrolysis”, “fucose metabolic process”, “protein N−linked glycosylation” corresponding to the REVIGO GO superclusters, “nucleic acid phosphodiester bond hydrolysis”, “fucose catabolic process”, and “fucosylation” in the treemap. For GLay module 2, the enriched terms “positive regulation of secondary cell wall biogenesis”, “regulation of peptide transport”, and “rRNA 2'−O−methylation” align with the REVIGO GO superclusters “negative regulation of mitotic cell cycle”, “nucleic acid phosphodiester bond hydrolysis”, and “vesicle−mediated transport to the plasma membrane”. GLay module 3 is associated with all the GO superclusters in the REVIGO treemap except for “fucosylation”. The associations between the module GO terms and the GO superclusters in pattern B reveal a coordinated interplay between cell cycle regulation, nucleic acid metabolism, and fucose-related processes, with “nucleic acid phosphodiester bond hydrolysis” being the predominant GO supercluster across the modules.

***Functional enrichments and hub genes of MCODE modules in pattern B.***

MCODE module 1 in pattern B is enriched in cell cycle-related biological processes, including mitotic cell cycle checkpoint signaling, regulation of cell cycle phase transition, negative regulation of cell cycle, mitotic DNA damage checkpoint signaling, and mitotic DNA integrity checkpoint signaling (Supplementary Dataset S5: Sheet 26). These cell cycle processes may promote graft unions by regulating cell division, maintaining genomic integrity, and supporting tissue formation and differentiation during plant grafting. For this module, we identified hub genes involved in DNA damage response (*Nbe.v1.1.chr11g05390*, a homolog of *BCP1*), vesicle transport along microtubule (*Nbe.v1.1.chr01g08340*, a homolog of *PAKRP2*), spindle assembly (*Nbe.v1.1.chr17g30120*, a homolog of *KINESIN-12E*), cell division cycle (*Nbe.v1.1.chr06g07540*, a homolog of *CDC45*) and transition of mitotic cell cycle (*Nbe.v1.1.chr16g44610*, a homolog of *CYC3B*) (Supplementary Dataset S5: Sheet 26).

The direct involvement of these genes in grafting has not been reported; however, their respective functions suggest that genome stability, precise cell division, and proliferation are required for graft union formation and tissue integration during grafting. *Nbe.v1.1.chr11g05390* is involved in DNA damage response processes and acts downstream of *SOG1* in homology-based repair. This gene is known to promote the recruitment of *RAD51* to DNA damage sites and facilitate homologous recombination and repair of double-strand breaks [228,229]. So, *BCP1* is involved in facilitating DNA damage response and repair, which can probably improve stress management and aid healing during grafting. *Nbe.v1.1.chr01g08340* has been identified to be involved in the microtubule-based movement, facilitating the transport of vesicles along phragmoplast microtubules during cell plate formation [230]. *PAKRP2,* through its ATP hydrolysis activity and its motility on single microtubules, is essential for efficient vesicle transport during cytokinesis [230], so it could help grafted plants manage the stress at the graft junction, enhancing graft success. *Nbe.v1.1.chr17g30120* (a homolog of *KINESIN-12E*) encodes a glycosyl transferase that is involved in the assembly of the mitotic spindle, a structure crucial for chromosome alignment and segregation during cell division [231]. During cytokinesis, *KINESIN-12E* localizes to the phragmoplast midzone, promoting cell plate formation and cell division [232]. The roles of *KINESIN-12E* in facilitating vesicle transport, cell plate formation, and cytokinesis suggest that it is probably involved in graft healing processes. *Nbe.v1.1.chr06g07540* (a homolog of *CDC45*) encodes the cell division cycle 45 protein known to be essential for DNA replication initiation, prior to meiosis for proper chromosome segregation and fertility, acting as a DNA polymerase α loading factor. *CDC45* is upregulated in young meiotic flower buds, indicating key roles in cell cycle progression and plant development [80]. Hence, *CDC45* is probably involved in grafted tissue growth and establishment.

MCODE module 2 in pattern B is related to spindle localization and organization (establishment of spindle/mitotic spindle localization, astral microtubule organization), chromosome movement and alignment (chromosome movement toward spindle poles, mitotic metaphase plate congression) and signal transduction pathway biological processes (Ran/Ras protein signal transduction, small GTPase-mediated signaling) (Supplementary Dataset S5: Sheet 26). These processes during grafting may ensure precise cell division and cytoskeletal reorganization at graft junctions, enabling vascular reconnection. For instance, the Ran/Ras protein signaling regulates the reassembly of the nuclear envelope, auxin transport, cell polarity, and growth [233,234], while spindle alignment maintains genomic stability during tissue regeneration [235], critical for successful graft union formation. For this module, we identified hub genes involved in stem fasciation (*Nbe.v1.1.chr09g16700*, a homolog of *FAS4*) and defense response (*Nbe.v1.1.chr02g29420*, a homolog gene encoding NB-ARC domain-containing disease resistance protein) (Supplementary Dataset S5: Sheet 26). The hub genes identified during plant grafting suggest they may be candidate genes for structural development and pathogen resilience following defense signaling. *Nbe.v1.1.chr09g16700* (a homolog of *FAS4*) encodes a FASCIATED STEM 4 protein with helicase activity involved in the development of abnormal, flattened-broader stems termed as fasciated stem [236]. This gene essentially regulates stem morphology, and its overexpression results in a fasciated stem phenotype [237]. The gene function on stem morphology and development suggests that during grafting, it may probably regulate changes in stem structure and integration, and vascular connection between grafted tissues. *Nbe.v1.1.chr01g00700* (a homolog gene encoding NB-ARC domain-containing disease resistance protein) encodes a protein that contains an NB-ARC domain, a nucleotide-binding adaptor shared by APAF-1, certain R gene products, and CED-4, and typically found in disease resistance proteins [179]. The nucleotide-binding site of the NB-ARC protein has ARC1 (Apoptotic protease activation factor 1 (APAF-1), cell death protein 4 (CED4) and ARC2 regions that activate downstream signaling targets [180,181]. This gene may play a crucial role in the immune response and disease resistance processes by recognizing and responding to pathogen attacks during grafting.

MCODE module 3 in pattern B is mainly enriched in carbohydrate metabolism processes (response to maltose, L-fucose, and fucose catabolic processes, L-fucose and fucose metabolic processes) and polyamine metabolism biological processes (spermidine biosynthetic and metabolic processes) (Supplementary Dataset S5: Sheet 26). These processes during grafting may ensure the provision of energy and enhance stress tolerance. In this module, we identified hub genes involved in root strip formation (*Nbe.v1.1.chr12g33170*, a homolog of *CASPL5C2*), phenylpropanoid and flavonoid biosynthesis (*Nbe.v1.1.chr11g20870*, a homolog gene encoding S-adenosyl-L-methionine-dependent methyltransferase superfamily protein), and membrane transport (*Nbe.v1.1.chr16g45560*, a homolog gene encoding transmembrane protein) (Supplementary Dataset S5: Sheet 26). These represent potential candidates that may influence molecular pathways such as the synthesis of secondary metabolites and cellular signaling or transport processes during grafting. *Nbe.v1.1.chr12g33170* (a homolog of *CASPL5C2*) encodes a protein that is part of the Casparian strip membrane proteins (CASP) family with known specific expression in various cell types, including trichomes, abscission zone cells, peripheral root cap cells, and xylem pole pericycle cells [84]. This gene is involved in the formation of the Casparian strip, a structure in the root endodermis for regulating water and nutrient transport between the soil and the vascular system [85]. The role of Casparian strip proteins is probably involved in regulating nutrient and water transport during grafting. *Nbe.v1.1.chr11g20870* (a homolog gene encoding S-adenosyl-L-methionine-dependent methyltransferase superfamily protein) is a key enzyme involved in the phenylpropanoid and flavonoid biosynthesis pathways. They regulate gene expression, protein function, and the secondary metabolites, which may be involved in plant growth, development, and stress responses [238].

MCODE module 4 in pattern B is mainly enriched in nucleotide metabolism (dGMP/dGDP/dATP metabolic processes, purine deoxyribonucleotide biosynthesis), glycoprotein transport, and calcium signaling biological processes (calcium-mediated signaling, ER calcium regulation) (Supplementary Dataset S5: Sheet 26). These processes, including nucleotide metabolism, may enhance DNA synthesis and cell division, glycoprotein transport may enable glycoprotein-mediated cell-cell adhesion, and coordinating calcium signaling may be involved in wound response. In this module, we identified hub genes involved in root peroxisome stress response (*Nbe.v1.1.chr06g21470*, a homolog of *PAO4*) and pathogen and hypoxia response (*Nbe.v1.1.chr17g07990*, a homolog gene encoding PADRE gene family protein) (Supplementary Dataset S5: Sheet 26), which may be important candidates for maintaining cellular homeostasis and defense response during grafting. *Nbe.v1.1.chr06g21470* (a homolog of *PAO4*) is an isoform in root peroxisomes involved in polyamine catabolism and enhances the oxidative conversion of spermine to spermidine. Its deficiency has been shown to alter the expression of drought stress response and flavonoid biosynthesis genes [239]. So, *PAO4* is probably involved in the stress response processes of grafting. *Nbe.v1.1.chr17g07990* (a homolog gene encoding the PADRE gene family protein) is known to be a prime candidate for functional mechanisms underlying plant disease resistance [240], so the PADRE gene family protein is probably involved in the defense response during grafting.

The remaining 5 MCODE modules in pattern B, while not meeting the stringent significant threshold (score > 5), also exhibit distinct, valuable network characteristics through their functional enrichments and key hub genes. To summarize, these MCODE modules encompass diverse biological processes, including “protein N-linked glycosylation” (MCODE 5, 24 nodes and 48 edges), “regulation of cellular component organization” (MCODE 6, 19 nodes and 35 edges), “positive regulation of secondary cell wall biogenesis” (MCODE 7, 28 nodes and 52 edges), “tracheary element differentiation” (MCODE 8, 22 nodes and 36 edges) and “cell cycle process” (MCODE 9, 10 nodes and 14 edges). *Nbe.v1.1.chr15g23790* (a homolog gene encoding tRNA/rRNA methyltransferase (SpoU) family protein), *Nbe.v1.1.chr03g32310* (a homolog of armadillo repeat only 4 gene, *ARO4*), *Nbe.v1.1.chr15g26880* (a homolog of phospholipase C2 gene, *PLC2*), *Nbe.v1.1.chr08g18160* (a homolog of *HSP17.6A*) and *Nbe.v1.1.chr11g35280* (a homolog of *MALE-STERILE 5*, *MS5*) were the hub genes identified in MCODE module 6 to module 9, respectively (Supplementary Dataset S5: Sheet 26). The functional enrichment profiles suggest the potential involvement of the modules in protein modification, cell wall biogenesis, xylem development and transport, and cell division.

***Functional enrichments and hub genes of GLay modules in pattern B.***

GLay module 1 in pattern B is enriched in RNA/nucleic acid metabolism (RNA modification, nucleic acid phosphodiester bond hydrolysis and RNA polymerase II/III regulation), carbohydrate metabolism (L-fucose/fucose catabolism), DNA Replication (RNA primer synthesis for mitotic/cell cycle DNA replication) and signaling biological processes (positive regulation of auxin biosynthesis and flavonoid regulation) (Supplementary Dataset S6: Sheet 32). During grafting, these biological processes, RNA modification and auxin biosynthesis, may regulate vascular reconnection and epigenetic signaling, carbohydrate metabolism may provide energy for cell wall repair, and DNA replication may enhance cell division at graft junctions. In this module, we identified hub genes involved in protein degradation (*Nbe.v1.1.chr11g36130*, a homolog gene encoding RNI-like superfamily protein), embryonic development (*Nbe.v1.1.chr11g32380*, a homolog of embryo defective 2739 gene, *EMB2739*), osmo-tolerance (*Nbe.v1.1.chr08g18160*, a homolog of *HSP17.6A*), and stress response and nuclear transcriptional activator (*Nbe.v1.1.chr10g01870*, a homolog of *C3H17*) (Supplementary Dataset S7: Sheet 38). The functions of the hub genes indicate their potential involvement in protein turnover, developmental processes, stress adaptation, and transcriptional regulation processes during grafting. *Nbe.v1.1.chr11g36130* (a homolog gene encoding RNI-like superfamily protein) is functionally related to protein degradation and mostly up-regulated in apomictic nucelli [241], so it is probably involved in regulating cell cycle and growth processes during grafting. *Nbe.v1.1.chr08g18160* (a homolog of *HSP17.6A*) encodes the cytosolic class II small heat-shock proteins (smHSP) in *A. thaliana*. The expression of the gene is known to be induced by heat and osmotic stress [242], suggesting the gene may be involved in stress regulation for water deprivation during grafting. *Nbe.v1.1.chr10g01870* (a homolog of *C3H17*) encodes a non-tandem CCCH zinc finger protein, which is ubiquitously expressed in a plant's life cycle and organ development [243]. The gene functions as a nuclear transcriptional activator and gene expression regulator and is identified to have pleiotropic effects on vegetative, flowering, seed development, and stress responses [243]. The function of *C3H17* suggests it is probably involved in RNA processing and stress management during grafting.

GLay module 2 in pattern B is enriched in cell wall biogenesis and morphogenesis (positive regulation of secondary cell wall organization/biogenesis, and regulation of cell morphogenesis), protein transport and localization (regulation of protein transport and negative regulation of lipoprotein metabolic processes), and tissue regeneration biological processes (Supplementary Dataset S6: Sheet 32). During plant grafting, these processes may influence graft success by strengthening cell walls and structural integrity through call wall biogenesis, ensuring cellular communication through protein transport, and promoting healing as a result of tissue regeneration. In this module, we identified hub genes involved in defense response (*Nbe.v1.1.chr06g26920*, a homolog gene encoding disease resistance protein (CC-NBS-LRR class) family), redox reactions (*Nbe.v1.1.chr06g33700*, a homolog gene encoding NAD(P)-binding Rossmann-fold superfamily protein), auxin and gravity induction regulation (*Nbe.v1.1.chr10g32330*, a homolog of *CNGC14*) and stress response and nuclear transcriptional activator (*Nbe.v1.1.chr03g12460*, a homolog of metacaspase 3 gene, *MC3*) (Supplementary Dataset S7: Sheet 2). These include novel hub genes that may be associated with the integration of defense, metabolic, hormonal, and stress-responsive mechanisms during plant grafting. *Nbe.v1.1.chr06g26920* encodes a disease resistance protein belonging to the CC-NBS-LRR class [244]. This protein is involved in the plant's defense mechanisms, particularly in recognizing and responding to pathogen attacks [244,245], so it is probably involved in the defense response during grafting. *Nbe.v1.1.chr06g33700* is involved in redox reactions and enzymatic processes [246]. In *Physcomitrium patens* (moss), homologous Rossmann-fold proteins are critical for chitosan-induced peroxidase activity and defense responses, as knockout lines showed reduced peroxidase activity and impaired expression of stress-responsive genes like *LOX* (lipoxygenase) [247]. Therefore, this gene probably supports graft healing through redox-mediated stress tolerance, peroxidase activation, and metabolic coordination. *Nbe.v1.1.chr10g32330* (a homolog of *CNGC14*) is a member of the cyclic nucleotide-gated channel family and is localized in the plasma membrane $Ca^{2+}$ channel. It is involved in the regulation of auxin-mediated growth responses, gravitropism in plant roots, and mediates calcium influx required for tip growth in root hairs [50,51]. The calcium signaling roles and auxin influx mediation of this gene suggest that it could be involved in cell signaling, communication, and tissue integration during grafting. *Nbe.v1.1.chr03g12460* (a homolog of *MC3*) encodes a metacaspase 3, a type of cysteine protease and phloem-specific member of the metacaspase family involved in drought tolerance, hypoxia resistance, and ABA signaling [248], unlike other metacaspases involved in programmed cell death (PCD) and pathogen defense [249,250]. This gene could therefore potentially influence the success of grafting by regulating phloem regeneration, proteolytic clearance, and stress response during grafting.

GLay module 3 in pattern B is enriched in cell cycle and division (cell cycle process, mitotic/meiotic cell cycle, nuclear division, cell cycle checkpoints and regulation chromosome organization and segregation), DNA and nucleotide metabolism processes (DNA recombination, repair, replication and nucleotide metabolism), cellular organization (organelle organization and fission, microtubule-based processes and vesicle transport) and carbohydrate and nitrogen compound metabolism biological processes (Supplementary Dataset S6: Sheet 32). During grafting, cell cycle regulation and DNA processes may be involved in cell division and differentiation of tissues. Cellular organization and vesicle transport biological processes may facilitate vascular reconnection, cell communication, and signaling pathways. In this module, we identified hub genes involved in cell division and callus proliferation (*Nbe.v1.s00110g00320*, a homolog gene encoding FKBP-like peptidyl-prolyl cis-trans isomerase family protein, *FKBP43*), cell cycle and division (*Nbe.v1.1.chr16g44610*, a homolog of *CYC3B* and *Nbe.v1.1.chr15g18390*, a homolog gene encoding transducin/WD40 repeat-like superfamily protein) (Supplementary Dataset S7: Sheet 38). Together, these genes are promising candidates for promoting effective cell division and callus formation during the grafting process. *Nbe.v1.s00110g00320* (a homolog of *FKBP43*) belongs to a group of chaperones in plants and is involved in several biochemical processes, including signal transduction [251] and protein folding [252,253]. *FKBP43* has been identified to regulate cell division, adhesion, and elongation during development and embryogenesis [254]. It is known to be required for the spatial organization of apical meristems [254], suggesting that *FKBP43* may play a key role in embryogenic callus proliferation during grafting. *Nbe.v1.1.chr16g44610* (a homolog of *CYC3B*) encodes a mitotic-like cyclin involved in regulating the cell cycle, particularly during the G2/M transition [255]. This gene is mainly expressed in tissues with active mitosis, such as roots and meristems [256,257], and is essential for controlling cell division, ensuring that cells progress through different phases of the cell cycle in a regulated manner [258]. This gene is probably involved in promoting cell division and growth during grafting.

GLay module 4 in pattern B is enriched in root and embryonic developmental processes (lateral root development, post-embryonic root development and post-embryonic plant organ development), reproductive regulation (negative regulation of flower development, photoperiodism-flowering, negative regulation of reproductive process and vegetative to reproductive phase transition of meristem), defense (response to gram-negative bacterium) and hormone response biological processes (response to brassinosteroid) (Supplementary Dataset S6: Sheet 32). During plant grafting, these biological processes may modulate defense responses, coordinate developmental transitions, mediate stress responses, and hormonal signaling. In this module, we identified a hub gene involved in the defense response to bacteria and regulation of immune response (*Nbe.v1.1.chr19g44270*, a homolog of *HOPZ-ACTIVATED RESISTANCE 1*, *ZAR1*) (Supplementary Dataset S7: Sheet 38). This hub gene has no previous knowledge regarding its specific function during plant grafting. *Nbe.v1.1.chr19g44270* (a homolog of *ZAR1*) is a canonical CC-type NLR protein conveyed by receptor-like cytoplasmic kinase sensors [259,260] involved in the immune response

[261], enabling defense mechanisms against bacterial infections, specifically *Pseudomonas syringae* [262,263]. ZAR, an immune receptor, has been identified to form a cation-selective channel resistosome, which is permeable to calcium. The activated *ZAR1* is known to form a pentamer in the plasma membrane, and its channel activity triggers immune signaling and cell death in plants [264]. *ZAR1* during grafting is likely involved in pathogen recognition, immune activation, and stress-induced cell death processes, which may facilitate wound healing through calcium-mediated signaling.

The MCODE and GLay modules in the network of pattern B showed overlapping hub genes and biological processes. *Nbe.v1.1.chr16g44610* (a homolog of *CYC3B*) and *Nbe.v1.1.chr15g18390* (a homolog gene encoding Transducin/WD40 repeat-like superfamily protein) were identified in MCODE module 1 and GLay module 3. Both were enriched in biological processes, "mitotic cell cycle checkpoint signaling" and "cell cycle process," suggesting these modules play crucial roles as cell division modulators. Additionally, *Nbe.v1.1.chr02g29420* and *Nbe.v1.1.chr15g19470* (homolog genes encoding NB-ARC domain-containing disease resistance protein) were identified in MCODE module 2 and across all GLay modules. The presence of NB-ARC domain-containing disease resistance proteins as hub genes in both module sets, along with enrichment for defense-related processes, highlights the importance of immune regulation during grafting.

**Pattern C**

***Gene profiles of pattern C.***

Pattern C shows genes that, compared to the starting point (intact), were predominantly highly expressed in the third state (14 to 28 DAG) only after grafting. It had the second-highest number of genes (1,133 genes) out of the six patterns. Genes in expression Pattern C comprise diverse functional classes including cell wall-related proteins (hydroxyproline-rich glycoprotein family protein, pectin lyase-like superfamily proteins and IRX15-like proteins), transcription factors (GATA, MYB, NAC, ethylene response factor, WRKY, bZIP), protein kinases (serine/threonine, receptor-like kinases, s-locus lectin protein kinases and cyclin-dependent kinase inhibitor), disease resistance proteins (NB-ARC domain-containing disease resistance proteins), RNA processing proteins (polynucleotidyl transferases, RNA polymerase II subunits and RNA binding family protein), transporters (ABC transporters, amino acid permeases, glucose-6-phosphate/phosphate translocator, nucleotide transporter) and signaling proteins (calcium-binding proteins, cyclic nucleotide-gated channels, regulator of G-protein signaling (RGS) proteins). This indicates grafting triggers significant changes in cell wall biogenesis, cellular signaling, and transport processes.

In the Bayesian gene regulatory network of pattern C, we identified some graft-related genes with known functional roles in plant grafting, including hormone signaling, transport and communication, stress responses, cell division, wound response, and cell wall remodeling. These genes related to such functional processes during grafting include genes encoding NAC domain-containing proteins (*NAC002*, *NAC007*, *NAC004*, *NAC029*, *NAC053* and *NAC060*), auxin efflux carrier family protein (*PIN3* and *PIN7*), basic helix loop helix (bHLH) DNA-binding superfamily protein (*bHLH121*), auxin canalization protein (*FL8*), BES1-interacting myc-like protein 2 (*BES1*), homeobox leucine zipper protein (*HAT14*), wound-responsive family protein (*WIP3*), early nodulin-like protein 9 (*ENODL9*), rhamnogalacturonate lyase family protein, mitotic-like cyclin 3B from *A. thaliana* (*CYC3B*) and cytokinin response factor 4 (*CRF4*) (Supplementary Dataset S8: Sheet 45).

The identified hub genes in pattern C are associated with a coordinated regulation of nucleic acid binding, peptide degradation, and RNA processing, depicting a framework where RNA metabolism and peptide turnover may be tightly linked. *Nbe.v1.1.chr15g14280* (a homolog gene encoding the zinc knuckle (CCHC-type) family protein), known for its role in RNA binding and post-transcriptional regulation [265,266], could influence the stability or translation of stress-responsive transcripts. *Nbe.v1.1.chr15g20950* (a homolog gene encoding the prolyl oligopeptidases (POPs) family protein, Peptidase S9), encodes a serine-type endopeptidase localized in the cytosol and is involved in the degradation of biologically important peptides such as peptide hormones [52]. This gene may modulate remodeling of the proteome at the graft interface by degrading or processing signaling peptides, peptide hormones, or damaged proteins generated during wounding and tissue regeneration, and may have a potential function under graft-induced stress. *Nbe.v1.1.chr06g34520* (a homolog of *TRZ4*), which encodes tRNase Z present in chloroplasts [53], potentially affects photosynthetic efficiency.

In this network for pattern C, 6 significant modules (threshold score > 5) were identified using the MCODE algorithm, and 4 modules were identified using the GLay algorithm. MCODE module 1 is mainly enriched in symbiont-mediated modulation of host defense response and programmed cell death-related biological processes. MCODE module 2 is mainly enriched in the regulation of synaptic signaling and chromosome organization-related biological processes. MCODE module 3 is mainly enriched in the regulation and metabolism of nucleic acids, organic compounds, and mitochondrial gene expression-related biological processes. MCODE module 4 is mainly enriched in the regulation of gravitropism, stomatal dynamics, and genome stability-related biological processes. MCODE module 5 is mainly enriched in the regulation of nucleic acid processes and telomere maintenance-related biological processes. MCODE module 6 is mainly enriched in cytoskeletal organization and stress signaling-related biological processes. GLay module 1 is mainly enriched in cytoskeletal organization, stress signaling, and interspecies interaction-related biological processes. GLay module 2 is mainly enriched in the regulation of nucleic acid metabolism, organic compounds, and DNA repair-related biological processes. GLay module 3 is mainly enriched in nucleic acid and nitrogen compound metabolism, DNA repair, damage response, and recombination-related biological processes. GLay module 4 is mainly enriched in the regulation and metabolism of sterols/steroids, and protein modification-related biological processes.

The GO superclusters identified through REVIGO treemap analysis for pattern C included “RNA modification”, “DNA damage response”, “positive regulation of chromosome organization”, “chromosome organization”, “plant organ morphogenesis”, “peroxisomal membrane transport”, “telomerase RNA stabilization”, “methylation”, “nuclear chromosome segregation”, “metabolic process”, “cell wall organization or biogenesis and “cellular process”. MCODE module 1 in pattern C enriched in “regulation of cell killing” is not associated with any of the GO superclusters in the REVIGO treemap. However, MCODE 2 is associated with the “chromosome organization” and “nuclear chromosome segregation” GO superclusters in the REVIGO treemap. MCODE module 3 enriched in “nucleic acid metabolic process” is associated with the “RNA modification” GO superclusters in the REVIGO treemap. MCODE module 4 enriched in “amyloplast organization” is not associated with any of the GO superclusters in the REVIGO treemap. MCODE module 5 enriched in “nucleic acid metabolic process” and “positive regulation of telomere maintenance” is associated with the “RNA modification”, “positive regulation of chromosome organization” and “methylation” GO superclusters in the REVIGO treemap. MCODE module 6 in pattern C enriched in “regulation of translational fidelity” and “aminoacyl−tRNA metabolism involved in translational fidelity” is associated with the “RNA modification” GO supercluster in the REVIGO treemap.

GLay modules 1 and 4 in pattern C, enriched in “entrainment of circadian clock” and “sterol biosynthetic process” biological processes, respectively, are not associated with any of the GO superclusters in the REVIGO treemap. GLay module 2 enriched in “nucleic acid metabolic process” and “positive regulation of chromosome organization” is associated with the “RNA modification”, “DNA damage response”, “positive regulation of chromosome organization”, and “telomerase RNA stabilization” GO superclusters in the REVIGO treemap. GLay module 3 is associated with all the GO superclusters in the tree map except for “plant organ morphogenesis”, “cell wall organization or biogenesis”, and “cellular process”. The associations between the module GO terms and the GO superclusters in pattern C reveal a coordinated interplay between “RNA modification”, “chromosome organization”, and “DNA damage response”.

***Functional enrichments and hub genes of MCODE modules in pattern C.***

MCODE module 1 in pattern C is enriched in cell death regulation (regulation of cell killing and modulation by symbiont of host programmed cell death), symbiont-host interaction, immune response, and stress regulation biological processes. (Supplementary Dataset S5: Sheet 27). These processes related to cell death regulation are essential for removing damaged cells at the graft interface, ensuring proper healing and vascular reconnection. Symbiont-host interaction mechanisms may influence graft compatibility and resistance to infections during the vulnerable stages of graft formation. Additionally, immune responses and stress regulation may help the plants to adapt to environmental stresses encountered during grafting. For this module, we identified hub genes involved in signal transduction (*Nbe.v1.1.chr13g10520*, a homolog gene encoding phosphatase 2C family protein, *PP2C*), transport processes (*Nbe.v1.1.chr15g31440*, a homolog gene encoding transmembrane protein), zinc homeostasis (*Nbe.v1.1.chr15g47660* and *Nbe.v1.1.chr05g05430*, homolog genes encoding major facilitator superfamily protein) (Supplementary Dataset S5: Sheet 27). These genes during plant grafting may be potential candidates for cellular signaling and homeostasis maintenance. *Nbe.v1.1.chr13g10520* (a homolog of *PP2C*) encodes a protein phosphatase 2C (*PP2C*) family protein. *PP2C* is involved in various signal transduction pathways, particularly in stress responses and ABA signaling [267,268]. They act as negative regulators of ABA signaling, which is crucial for plant responses to drought and other stress conditions [267]. *PP2C* also interacts with mitogen-activated protein kinase (MAPK) pathways, influencing growth and development [268]. So, *PP2C* regulation of ABA signaling and stress responses may be involved in environmental stress adaptation processes, improve healing at the graft interface, and ensure successful vascular reconnection. *Nbe.v1.1.chr15g31440* encodes a transmembrane protein (AT4G17250) involved in heat acclimation, and its expression increased after 37 °C heat treatment in *A. thaliana* suspension-culture cells [269]. In the context of plant grafting, the transmembrane protein may enhance adaptation to environmental conditions and improve stress resistance. *Nbe.v1.1.chr15g47660* and *Nbe.v1.1.chr05g05430* are involved in transporting a wide range of small organic molecules across cell membranes [270,271]. Specifically, this protein plays a role in zinc homeostasis, influencing zinc tolerance and accumulation (Haydon & Cobbett, 2007), so the protein could facilitate essential nutrient transport and molecular signaling processes during plant grafting.

MCODE module 2 in pattern C is enriched in synaptic transmission (synaptic vesicle priming, trans-synaptic signaling, signal release from synapse, and synaptic vesicle exocytosis), cellular, chromosome, and nuclear organization, biological processes. These processes may regulate intercellular signaling and communication and alter the expression of genes during grafting. For this module, we identified hub genes involved in the regulation of heterochromatic DNA replication and cell cycle (*Nbe.v1.1.chr13g22810*, a homolog of *ATXR6*) and zinc ion binding functioning as E3 ubiquitin ligases (*Nbe.v1.1.chr15g07140*, a homolog gene encoding RING/FYVE/PHD zinc finger superfamily protein) (Supplementary Dataset S5: Sheet 27). The regulatory function of these hub genes indicates their novel suggested roles in maintaining genomic stability, protein turnover, and facilitating abiotic stress resilience during plant grafting. *Nbe.v1.1.chr13g22810* (a homolog of *ATXR6*) encodes a SET-domain protein involved in histone H3K27 monomethylation [272,273]. This protein plays a crucial role in regulating chromatin structure and gene silencing, particularly during DNA replication and cell cycle progression [273–275]. *ATXR6* is essential for maintaining heterochromatin integrity and proper gene expression; therefore, it may be involved in the regulation of gene expression and chromatin remodeling processes. *Nbe.v1.1.chr15g07140* (a homolog gene encoding RING/FYVE/PHD zinc finger superfamily protein) is primarily involved in zinc ion binding and functions as an E3 ubiquitin ligase. They play crucial roles in regulating growth, development, and responses to abiotic stresses such as drought, salinity, and temperature [276]. This gene, through its role in ubiquitination and stress tolerance, may be involved in stress responses and the cellular homeostasis process of grafting.

MCODE module 3 in pattern C is enriched in nucleic acid and RNA metabolism biological processes, including nucleic acid phosphodiester bond hydrolysis, RNA metabolic process, negative/positive regulation of antisense RNA transcription, RNA metabolic process, RNA modification and nucleic acid metabolic process. These processes related to RNA and nucleic acid metabolism may regulate the expression of graft-related genes during grafting. For this module, we identified hub genes which may be candidates for grafting and involved in mitochondrial transcription (*Nbe.v1.1.chr19g27060*, a homolog gene encoding DNA/RNA polymerases superfamily protein), DNA repair (*Nbe.v1.1.chr17g09430*, a homolog of *POLK*), mediates protein-protein interactions (*Nbe.v1.1.chr18g27360*, a homolog gene encoding ankyrin repeat family protein), dehydration stress response (*Nbe.v1.1.chr02g03460*, a homolog of *OSCA2.3*), regulation of gene expression (*Nbe.v1.1.chr02g08930*, a homolog of *LUH*) and peroxisomal biogenesis (*Nbe.v1.1.chr02g29950*, a homolog of *PEX6*) (Supplementary Dataset S5: Sheet 27). *Nbe.v1.1.chr19g27060* (a homolog gene encoding DNA/RNA polymerases superfamily protein) is a nucleus-encoded plastid RNA polymerase involved in mitochondrial and chloroplast transcription [277–279]. This gene is involved in the synthesis of RNA from DNA templates, which is essential for gene expression and the maintenance of cellular functions [277–280]. Essential for RNA synthesis in the mitochondria and chloroplasts, this gene may probably regulate transcription linked to metabolic and energy requirements during plant grafting. *Nbe.v1.1.chr17g09430* (a homolog of *POLK*) encodes DNA polymerase kappa involved in DNA repair processes [281]. POLK plays a crucial role in translesion synthesis, allowing DNA replication beyond damaged sites and ensuring DNA polymerases high fidelity [282,283]. This gene may be involved in maintaining genomic stability during the process of plant grafting. *Nbe.v1.1.chr18g27360* (a homolog gene encoding ankyrin repeat family protein) is involved in mediating protein-protein interactions [284]. Among the 37 predicted ankyrin-repeat transmembrane proteins in *A. thaliana* [285], accelerated cell death 6 (*ACD6*) and increased tolerance to NaCl (*ITN1*) have previously been characterized as positive regulators of SA signaling in local defense responses [286] and involved in salt stress tolerance [287], respectively. The findings suggest that the gene may enhance both biotic and abiotic stress responses, potentially influencing graft success through enhanced stress resilience and signaling regulation. *Nbe.v1.1.chr02g03460* (a homolog of *OSCA2.3*) encodes a hyperosmolality-gated calcium-permeable channel family [183] involved in osmotic adjustments [183,184], with differential roles in stress response and tolerance [185,186], and so *OSCA2.3* is probably involved in osmo-sensing and stress response during grafting. *Nbe.v1.1.chr02g08930* (a homolog of *LUH*) encodes a LEUNIG-like protein similar to LEUNIG (*LUG*) [288] and functions in embryonic development, floral homeotic gene regulation, and plant biotic and abiotic stress responses [289]. The gene is also known to be involved in cell wall modifications necessary for mucilage extrusion and mediates aluminum sensitivity through modulated root cell wall pectin methyl-esterification [290,291]. Hence, *LUH* may probably be involved in modulating stress responses during grafting. *Nbe.v1.1.chr02g29950* (a homolog of *PEX6*) encodes a protein involved in peroxisome biogenesis and maintenance, essential organelles for plant metabolic processes [216]. *PEX6* is involved in the breakdown of fatty acids stored in vital oil bodies essential for seed germination and seedling development [217]. *PEX6* is probably involved in controlling physiological stresses associated with grafting by maintaining peroxisome function and facilitating stress responses and energy supply.

MCODE module 4 in pattern C is enriched in protein catabolism and ubiquitination, gravitropism, stomatal closure, amyloplast organization, histone modification, and DNA repair biological processes. These processes related to gravitropism and stomatal closure may influence water and nutrient balance during grafting. Histone modifications and DNA repair may regulate gene expression and genomic stability during grafting. For this module, we identified a hub gene involved in protein ubiquitination (*Nbe.v1.1.chr12g26540*, a homolog of *NHL8*) (Supplementary Dataset S5: Sheet 27). *Nbe.v1.1.chr12g26540* (a homolog of *NHL8*) encodes the NDR1/HIN1-like 8, an E3 ubiquitin-protein ligase containing an IBR and RWD domains [292]. This gene belongs to the subclass of RING E3 ligases and has been identified to be similar to the tobacco *HIN1* (harpin-induced) hairpin-induced gene and the *A. thaliana* NDR1 (non-race-specific disease resistance) [292,293]. *NHL8* has been identified to be involved in plant defense mechanisms against pathogens observed in *A. thaliana*, by conferring resistance to cyst nematode in soybean [292]. *NHL8* may be involved in modulating defense mechanisms and stress response during grafting.

MCODE module 5 in pattern C is enriched in nucleic acid and chromosome regulation biological processes (regulation of termination of DNA-templated transcription, nucleic acid metabolic process, regulation of DNA repair and chromosome organization), RNA metabolism and modification, protein and chemical modifications (peptidyl-lysine methylation and protein methylation/alkylation), and stress responses (response to water deprivation and salt stress). These biological processes may regulate gene expression, ensure efficient cell division, and stress responses during grafting. For this module, we identified hub genes involved in histone methylation and response to vernalization (*Nbe.v1.1.chr08g03260*, a homolog of *ASHH3*), regulation of iron homeostasis (*Nbe.v1.1.chr07g03980*, a homolog of *bHLH121*), DNA recombination, repair and replication (*Nbe.v1.1.chr07g29080*, a homolog of *RPA70B*), DNA repair and reciprocal meiotic recombination (*Nbe.v1.1.chr15g19050*, a homolog of *DMC1*), embryo development (*Nbe.v1.1.chr05g16810*, a homolog of *MEE47*) and activation of aurora kinase activity (*Nbe.v1.1.chr10g38460*, a homolog of *TPXL3*) (Supplementary Dataset S5: Sheet 27). The hub genes identified and characterized by their potential functions during grafting, including nutrient homeostasis, genome maintenance, and development, directly correlated with the GO terms of the module. *Nbe.v1.1.chr08g03260* (a homolog of *ASHH3*) encodes a histone-lysine N-methyltransferase located in the endoplasmic reticulum and involved in the methylation of histone H3 at lysine 36 (H3K36me2/3) [294], proper timing of *VERNALIZATION INSENSITIVE 3* (*VIN3*) induction for the vernalization processes [295,296]. Furthermore, a crosstalk between DNA methylation, sRNAs accumulation, and histone modifications has been established such that at the chromatin structure level, epigenetic changes induced by distant signaling are expected to occur [297,298]. Grafting has been shown to induce histone and DNA modifications, resulting in altered gene expression patterns that eventually lead to vigorous shoot phenotype in grapevine heterografts [299], flower induction in apple grafts [300], somatic embryogenesis in *Hevea brasiliensis* grafts [301], and drought stress tolerance in tomato self-grafts [302]. Hence, this histone methylation-associated gene may be involved in the regulation of epigenetic marks during grafting and may mitigate stress tolerance during the wound-healing process for improved growth and development. *Nbe.v1.1.chr07g03980* (a homolog of *bHLH121*) is involved in the iron deficiency signaling pathway [303,304] such that under iron deficiency, the phosphorylated form of *bHLH121* accumulates and forms heterodimers with iron-regulated transporter 1 (*IRT1*) and ferric reduction oxidase 2 (*FRO2*), which induces iron uptake and maintains iron homeostasis [304–307]. Recent studies have revealed that grafting creates transient water and nutrient-deficiency states due to disrupted vascular connectivity [134,308]. New grafts subjected to a low-humidity environment wilted due to severed vascular strands, but they regained the ability to transport water and maintain turgor only after xylem continuity [134]. Further, a high-stress graft model, *N. benthamiana* and *A. thaliana* interfamily heterograft, revealed that grafting creates a transient nutrient-deficient period after grafting, with recovery timing dependent on graft compatibility. In the low-stress model, *N. benthamiana* homografts, nutrient transport recovered by 7 DAG, while in interfamily heterografts, nutrient transport remained severely impaired even after 7 DAG, with the formation of autophagic structures in cells near the graft boundary over a long period, indicating prolonged nutrient stress [308]. Iron, which is a critical micronutrient for redox enzymes, the synthesis of new cellular components, and stress adaptation [309,310], is likely depleted during this vascular disruption phase. The *bHLH121* homolog gene, which responds to iron deficiency by regulating the expression of iron uptake genes, may play a pivotal role under iron-deficient conditions by regulating iron homeostasis, maintaining adequate iron levels, and improving oxidative and environmental stress adaptation during grafting. *Nbe.v1.1.chr07g29080* (a homolog of *RPA70B*) encodes the replication protein A 70 kDa subunit B, a homolog of replication protein A, involved in DNA repair, recombination, and transcription [311]. This gene, *RPA70B,* through DNA replication and repair, may be involved in genomic stability maintenance processes during plant grafting. *Nbe.v1.1.chr15g19050* (a homolog of *DMC1*), supported by *RAD51* for meiotic recombination, is involved in recombination and DNA double-strand breaks repair following homologous recombination, accurate chromosome segregation, and genetic diversity during meiosis [312–314]. *DMC1* could play a role in maintaining genomic stability and facilitating DNA repair during plant grafting. *Nbe.v1.1.chr05g16810* (a homolog of *MEE47*) encodes the maternal effect embryo arrest 47 protein involved in embryo development, leading to seed dormancy and regulation of DNA-templated transcription [315–317]. This gene may be involved in the regulation of gene expression during grafting. *Nbe.v1.1.chr10g38460* (a homolog of *TPXL3*) is a primary activator of α-Aurora kinases (*AUR1* and *AUR2*), involved in embryogenesis, cell cycle-dependent localization on microtubule arrays, spindle assembly, and mitotic progression [218,318]. This gene may be involved in microtubule organization and cell division during grafting.

MCODE module 6 in pattern C is enriched in biological processes, including cytoskeletal and cellular organization (regulation of actin filament bundle assembly, cristae formation, and tubulin deacetylation), stress response and signal transduction (positive regulation of stress-activated MAPK cascade, JNK cascade, and stress-activated MAPK cascade), and RNA and protein metabolism (aminoacyl-tRNA metabolism, regulation of translational fidelity, and cysteinyl-tRNA aminoacylation). These biological processes may regulate cytoskeletal organization, promote cellular alignment, and ensure efficient gene expression during plant grafting. For this module, we identified hub genes involved in auxin transport and vascular patterning (*Nbe.v1.1.chr13g00825*, a homolog of *FL8*), chloroplast size and localization (*Nbe.v1.1.chr01g33940*, a homolog of *REC1*), mitochondrial cristae integrity and regulation of programmed cell death (*Nbe.v1.1.chr13g06550*, a homolog of *SYCO*) and programmed cell death (*Nbe.v1.1.chr05g05310*, a homolog of *ZAT14*) (Supplementary Dataset S5: Sheet 27). The hub genes identified in this module may be crucial candidates for coordinating hormone signaling, organelle dynamics, and cell fate processes during graft formation and vascular regeneration to achieve functional tissue union. *Nbe.v1.1.chr13g00825* (a homolog of *FL8*), a member of the *FORKED-LIKE* (*FL*) genes involved in the secretory pathway and localization of the auxin efflux protein (PIN1) in pro-vascular tissue, including developing veins [98,99]. This gene may be involved in vascular differentiation and auxin transport during grafting. *Nbe.v1.1.chr01g33940* (a homolog of *REC1*) is cytosol and nucleus, accumulating extra plastidic protein involved in establishing the chloroplast size compartment and regulating cellular volume devoted to chloroplast [319,320]. The gene is crucial for proper chloroplast distribution and function and may be involved in efficient chloroplast coverage and photosynthesis during grafting.

*Nbe.v1.1.chr13g06550* (a homolog of *SYCO*) encodes the cysteinyl t-RNA synthetase localized to mitochondria [321] and has been identified to be involved in mitochondrial cristae integrity and programmed cell death [321–324]. The gene may be involved in mitochondrial functions essential for energy production and cellular metabolism during grafting. Also, the programmed cell death function may aid in pathogen defense and the removal of damaged cells during plant grafting. *Nbe.v1.1.chr05g05310* (a homolog of *ZAT14*) encodes a C2H2 zinc finger protein involved in developmental programmed cell death (dPCD) in *A. thaliana* [325]. *ZAT14* has been identified as a transcriptional repressor within the gene regulatory network of root cap dPCD, acting downstream of *ANAC033*/*SOMBRERO* (*SMB*), a key regulator of root cap dPCD [325]. The role of the gene in regulating programmed cell death is crucial for removing damaged cells, which may facilitate the integration and healing of tissues during grafting.

A total of twelve MCODE modules in the network of pattern C did not meet the threshold score > 5; however, their important characteristics, including node counts and functional enrichments, are briefly described below. These modules except for modules 15 and 17 were enriched in "zinc ion transmembrane transport" (MCODE 7, 85 nodes and 180 edges), "cell wall mannoprotein biosynthetic process" (MCODE 8, 86 nodes and 179 edges), "thylakoid membrane organization" (MCODE 9, 64 nodes and 131 edges), "entrainment of circadian clock" (MCODE 10, 65 nodes and 131 edges), "disruption of the cellular component of another organism" (MCODE 11, 40 nodes and 79 edges), "plant ovule development" (MCODE 12, 5 nodes and 8 edges), "embryonic pattern specification" (MCODE 13, 53 nodes and 98 edges), "double-strand break repair" (MCODE 14, 37 nodes and 58 edges), "RNA modification" (MCODE 16, 50 nodes and 78 edges) and "DNA repair complex assembly" (MCODE 18, 3 nodes and 3 edges) (Supplementary Dataset S5: Sheet 27).

The MCODE modules 7–16 are characterized by hub homologous genes encoding proteins involved in stress response, autophagy, metabolism, transport, signaling, and genome maintenance. They include *Nbe.v1.1.chr13g34560* (a homolog of ABA overly-sensitive 5 gene, *ABO5*), *Nbe.v1.1.chr11g26190* (a homolog of yeast autophagy 18F gene, *ATG18*), *Nbe.v1.1.chr03g30490* (a homolog of abietane diterpene oxidase 1 gene, *ADTO1*), *Nbe.v1.1.chr09g01630* (a homolog of *DTX29*), *Nbe.v1.1.chr03g08880* (a homolog of *MDKIN1*), *Nbe.v1.1.chr07g10540* (a homolog of karyopherin enabling the transport of the cytoplasmic hyl1 gene, *KETCH1*), *Nbe.v1.1.chr12g03330* (a homolog of $k^+$ efflux antiporter 4 gene, *KEA4*), *Nbe.v1.1.chr16g08080* (a homolog gene encoding NIN-like protein 8, *NLP8*), *Nbe.v1.1.chr04g06860* (a homolog gene encoding DNAJ heat shock N-terminal domain-containing protein) and *Nbe.v1.1.chr05g32170* (a homolog gene encoding sensitivity to red light reduced protein, *SRR1*) (Supplementary Dataset S5: Sheet 27). The modules based on their seed nodes, hub genes, and enriched biological processes are potentially involved in nutrient transport, cell wall biosynthesis, chloroplast structure organization, circadian rhythm coordination, inter-organismal interactions, reproduction, and genome stability and integrity.

***Functional enrichments and hub genes of GLay modules in pattern C.***

GLay module 1 in pattern C is enriched in biological processes including entrainment of the circadian clock, cell wall disruption, signal transduction and stress response (positive regulation of JNK cascade and positive regulation of stress-activated MAPK cascade), regulation of actin filament bundle assembly, DNA and RNA processes (group II intron splicing and DNA damage checkpoint signaling) (Supplementary Dataset S6: Sheet 33). In this module, we identified hub genes associated with a diverse array of stress responses and physiological processes, reflecting the complex adaptations required during graft union formation and healing. These include genes involved in xenobiotic detoxification (*Nbe.v1.1.chr09g01630*, a homolog of *DTX29*), auxin response (*Nbe.v1.1.chr03g28630*, a homolog of *ITPK1*), phosphate (Pi) homeostasis and Pi loading into root xylem (*Nbe.v1.1.chr18g29410*, a homolog of *PHO1;H3*), immune response (*Nbe.v1.1.chr16g33430*, a homolog of *RFO3*), drought and salt tolerance (*Nbe.v1.1.chr12g20590*, a homolog gene encoding calcium-dependent lipid-binding protein, *CLB*) and response to pathogens (*Nbe.v1.1.chr05g32030*, a homolog gene encoding UvrABC system protein C) (Supplementary Dataset S7: Sheet 39). *Nbe.v1.1.chr09g01630* (a homolog of *DTX29*) encodes a MATE (multidrug and toxic compound extrusion) family protein localized to the vacuolar membrane and involved in detoxification processes and functions as a xenobiotic transporter (exporting toxic compounds out of the cell) [326]. This gene may be involved in the efflux of exogenous toxic compounds and cellular homeostasis during grafting. *Nbe.v1.1.chr03g28630* (a homolog of *ITPK1*) is involved in phosphate signaling [327] and auxin responses [328]. Phosphorylation activities of the gene produce inositol pyrophosphates like 5-InsP7 and InsP81 [327] involved in maintaining phosphate homeostasis [329] and auxin-related processes such as root elongation, root hair development, leaf venation, and gravitropic responses [328]. This gene may probably regulate phosphate signaling and auxin responses during grafting. *Nbe.v1.1.chr18g29410* (a homolog of *PHO1;H3*) encodes a phosphate transporter expressed in root vascular cylinder cells [330], upregulated in response to zinc (Zn) deficiency and involved in Pi homeostasis [331] and Pi loading into xylem root [332]. This gene may probably regulate phosphate transport and homeostasis during grafting. *Nbe.v1.1.chr12g20590* (a homolog of *CLB*) encodes a calcium-dependent lipid-binding protein containing a C2 domain, which acts as a transcriptional regulator and binds specifically to the promoter of *THAS1* (thalianol synthase 1) [333]. The gene is involved in drought and salt tolerance [334], so it may be involved in stress response and tolerance during grafting.

GLay module 2 in pattern C is enriched in biological processes, including nucleic acid and RNA metabolism (RNA metabolic process and modification, DNA recombination, nucleic acid phosphodiester bond hydrolysis, and mitochondrial gene expression) and DNA repair and stress response (Supplementary Dataset S6: Sheet 33). In this module, we identified hub genes potentially implicated in essential grafting processes including secondary metabolite biosynthesis and response to environmental stresses (*Nbe.v1.1.chr14g20700*, a homolog gene encoding the 2-oxoglutarate (2OG) and Fe(II)-dependent oxygenase superfamily protein), peroxisomal biogenesis (*Nbe.v1.1.chr02g29950*, a homolog of *PEX6*), protein degradation (*Nbe.v1.1.chr18g27010*, a homolog gene encoding the F-box/RNI-like superfamily protein), protein ubiquitination (*Nbe.v1.1.chr19g02960*, a homolog of *PP2-B1*), DNA repair and reciprocal meiotic recombination (*Nbe.v1.1.chr15g19050*, a homolog of *DMC1*), chromatin remodeling and localized to the plasmodesma (*Nbe.v1.1.chr16g29610*, a homolog gene encoding set-domain containing protein lysine methyltransferase family protein, *SUVR4*), cell differentiation (*Nbe.v1.1.chr19g07140*, a homolog of *RIMA*), response to starvation (*Nbe.v1.1.chr11g26190*, a homolog gene encoding yeast autophagy 18 F-like protein, *G18F*) and defense response and signal transduction (*Nbe.v1.1.*chr07g03580, a homolog gene encoding disease resistance protein (TIR-NBS-LRR class) family) (Supplementary Dataset S7: Sheet 39). *Nbe.v1.1.chr14g20700* (a homolog gene encoding 2-oxoglutarate (2OG) and Fe(II)-dependent oxygenase superfamily protein) has been identified to be involved in the hydroxylation of various substrates crucial for secondary metabolites biosynthesis and environmental stress response [290,335]. This gene may be involved in secondary metabolite biosynthesis and aid in stress response during grafting. *Nbe.v1.1.chr19g02960* (a homolog of *PP2-B1*) encodes an F-box protein, which is a component of the SCF (SKP1-Cullin-F-box) E3 ubiquitin ligase complex involved in ubiquitination and proteasomal degradation of target proteins. This process regulates cell cycle progression, signal transduction, and stress responses [336]. The expression of PP2 is related to phloem differentiation [337], mesophyll plasmodesmata interaction for cell-to-cell trafficking [338], and this predicts the intercellular movement of PP2 within the sieve element-companion cell complex [337,339]. This gene may be involved in cellular homeostasis through protein degradation, enhancing phloem differentiation and transport during grafting. *Nbe.v1.1.chr16g29610* (a homolog of *SUVR4*) encodes a nucleolar histone methyltransferase with monomethylated histone H3K9 as the preferred substrate involved in chromatin remodeling. *SUVR4* represses rDNA in the decondensed part of the nucleolus [340,341] and regulates gene expression by binding ubiquitin and converting H3K9me1 to H3K9me3 [342]. Thus, this gene may have a role in efficient gene expression during grafting. *Nbe.v1.1.chr11g26190* (a homolog of *G18F*) encodes an autophagy-related protein located in the ER and plasma membranes [343] for cellular homeostasis maintenance and is induced by nutrient starvation. It is expressed in roots, flowers, and leaves [344] and responds to a combination of heat and drought stress [345]. Also, autophagy has been reported to be induced during grafting and wound healing to promote callus formation and contribute to tissue connectivity [308]. This gene may be involved in autophagy and stress responses during grafting. *Nbe.v1.1.chr07g03580* encodes a member of the TIR-NBS-LRR class of disease resistance proteins, which are involved in recognizing pathogen-associated molecular patterns (PAMPs), triggering defense responses, and promoting plant innate immunity [346–349]. This gene may be involved in the activation of the immune system in grafted plants, anticipating future infections and preventing pathogen attacks.

GLay module 3 in pattern C is enriched in biological processes, including RNA and nucleic acid metabolism (nucleic acid and nucleobase-containing compound metabolic processes), heterocycle and cellular nitrogen and aromatic compound metabolic processes (Supplementary Dataset S6: Sheet 33). The hub genes screened in the module are involved in nucleic acid and zinc ion binding (*Nbe.v1.1.chr15g14280*, a homolog gene encoding zinc knuckle (CCHC-type) family protein), proteolysis (*Nbe.v1.1.chr15g20950*, a homolog gene encoding prolyl oligopeptidase family protein), mitochondrial tRNA 3'-end processing (*Nbe.v1.1.chr06g34520*, a homolog of *TRZ4*), brassinosteroid biosynthesis (*Nbe.v1.1.chr03g30490*, a homolog of *ADTO1*), tRNA methylation (*Nbe.v1.1.chr17g33670*, a homolog of *TRM2B*) and translocation of cytoplasmic proteins into plastids

(*Nbe.v1.1.chr11g03340,* a homolog of secy homolog 2 gene, *SCY2*) (Supplementary Dataset S7: Sheet 39). The regulation of the hub genes highlights their probable roles in hormone biosynthesis, organelle function, and nucleic acid modification in supporting the complex physiological changes that occur during grafting. *Nbe.v1.1.chr15g14280* encodes a protein involved in nucleic acid binding and zinc ion binding and nucleic acid metabolism (transcription, RNA processing, and post-transcriptional gene silencing) [265]. This gene may be involved in gene expression and RNA processing during grafting. *Nbe.v1.1.chr15g20950* is a serine peptidase involved in proteolysis, confers abiotic stress tolerance, protein maturation, and degradation by hydrolyzing short peptides [350,351]. The role of the gene in proteolysis may be involved in cellular function and stress adaptation during grafting. *Nbe.v1.1.chr03g30490* encodes a monooxygenase enzyme that catalyzes the oxidation of abietane diterpenes, a precursor to brassinosteroid essential for regulating cell elongation, vascular differentiation, and stress response regulation [352–354]. This gene may be involved in cell division, cell elongation, and tissue differentiation during grafting. *Nbe.v1.1.chr17g33670* (a homolog of *TRM2B*) encodes a tRNA-specific methyltransferase responsible for the methylation of uridine at the 5th position (m5U) in tRNA and ensures the stability and proper functioning of tRNA essential for protein synthesis [70,355]. This gene is essential for proper protein synthesis and may probably enhance cellular growth during grafting. *Nbe.v1.1.chr11g03340* encodes a component of the sec translocase system involved in the translocation of cytoplasmic proteins into plastids, specifically the inner envelope membrane, and is essential for early plant development [356]. This may probably be involved in plastid protein translocation.

GLay module 4 in pattern C is enriched in biological processes, including sterol/steroid biosynthesis and metabolism, protein modification and regulation (protein peptidyl-prolyl isomerization, peptidyl-proline modification, protein polyubiquitination, and regulation of proteolysis) (Supplementary Dataset S6: Sheet 33). In this module, we identified hub genes that are potential candidates for key regulatory pathways during grafting, such as defense response *Nbe.v1.1.chr10g15120* and *Nbe.v1.1.chr03g44010* (homologs of *SDA1*), and gibberellic acid biosynthesis *Nbe.v1.1.chr13g28940* (a homolog of jumonji domain-containing protein 20 gene, *JMJ20*) (Supplementary Dataset S7: Sheet 39). *Nbe.v1.1.chr10g15120* and *Nbe.v1.1.chr03g44010* (homologs of *SDA1*) encodes a novel small plant-specific protein containing a highly conserved seven amino acid (S/G)WA(D/E)QWD domain at the N-terminus that is involved in modulating pathogen defense and tolerance to oxidative stress [357]. *SDA1* regulates plant immunity via the SA-mediated defense pathway and tolerance to oxidative stress. This gene may be involved in stress responses and defense mechanisms during grafting. *Nbe.v1.1.chr13g28940* is a histone arginine demethylase essential for photomorphogenesis and positively regulates seed germination following histone arginine demethylation of gibberellin 3 β-hydroxylase 1 (*GA3ox1*) and gibberellin 3 β-hydroxylase 2 (*GA3ox2*) [358]. This gene, during grafting, may probably be involved in targeted demethylation of GA biosynthesis genes in light and developmental responses.

The MCODE and GLay modules in pattern C revealed overlapped hub genes and biological processes. *Nbe.v1.1.chr15g14280* (a homolog gene encoding zinc knuckle (CCHC-type) family protein), *Nbe.v1.1.chr15g20950* (a homolog gene encoding prolyl oligopeptidase family protein), and *Nbe.v1.1.chr06g34520* (a homolog of *TRZ4*) were found in both MCODE module 3 and GLay module 3, suggesting these modules are potential modulators for RNA binding, processing, and post-transcriptional regulation. *Nbe.v1.1.chr02g29950* (a homolog of *PEX6*) was identified in both MCODE module 3 and GLay module 2. Both clustering methods revealed overlapped enriched biological processes, including nucleic acid metabolic processes and RNA metabolism, in the MCODE and GLay modules. This identifies the core modulators and regulatory processes underlying pattern C.

**Pattern D**

***Gene profiles of pattern D.***

Pattern D genes, compared to the starting point (intact), had lower expression levels in the third state (Ave 14–28 DAG) only after grafting. Genes in expression pattern D comprise diverse functional classes, including protein kinases (putative protein kinase 1, leucine-rich repeat protein kinase family protein and serine/threonine-kinase), transcription factors (NAC, transcription factor IIIB and G-box binding factor 3, E2F transcription factor 3), cell wall modification (glycosyl transferase family 35, beta galactosidase 1 and cellulose-synthase-like C6), membrane transporters ($K^+$ uptake transporter 3, phosphate transporter 3;1 and cation/H+ exchanger 18) and organelle-targeted proteins (plastid-lipid associated protein and plastid transcriptionally active 3). In the Bayesian gene regulatory network of pattern D, we identified genes repressed during grafting with known functions such as gene regulation, cell cycle regulation, and cell wall remodeling/modification, including genes encoding *NAC100*, *CYCA3;4* and *BXL6* (Supplementary Dataset S8: Sheet 46). Notably, these genes were upregulated during the early phase after grafting but repressed at the later stage, indicating a dynamic regulatory shift during grafting. This temporal expression suggests that while these genes may contribute to initial cellular responses such as cell proliferation, signaling, or cell wall remodeling immediately after grafting, their rapid repression may indicate mirror events of graft healing, including the suppression of non-essential cell cycle processes, stress responses that facilitate transition from tissue proliferation to differentiation and integration required for graft union formation.

While most grafting research focuses on upregulated processes that directly link to graft healing and graft-associated genes, our current analysis also included regulatory gene networks and modules for mirroring events of graft healing that are suppressed during grafting in patterns D, E, and F. The inclusion of these gene modules in our network analysis highlights the importance of suppressing such events to promote grafting. For example, while some NAC gene family proteins, including *VND1* and *VND7,* involved in xylem differentiation are upregulated and promote graft healing, others, like *NAC100,* are downregulated. This indicates a molecular trade-off where different NACs play opposing roles, suggesting a molecular competition between promoting cell differentiation and maintaining stem cell fate. Similarly, *BXL6,* a cell wall remodeling and degradation enzyme [359], is upregulated during the early phase after grafting but rapidly repressed at the later stage, which may be necessary to prevent excessive wall degradation, maintain cell wall rigidity, and facilitate organized vascular reconnection following cell division and callus formation.

Thus, our network analysis provides valuable insight into the importance of repressed gene activities, which may allow for the upregulation of genes that promote the processes required for effective graft integration.

In this network for pattern D, 4 significant modules (threshold score > 5) were identified using the MCODE algorithm and 3 modules using the GLay community clustering algorithm. MCODE module 1 genes are mainly enriched in meiotic cell cycle regulation, DNA replication, and membrane organization-related biological processes. The enrichment of these processes by the repressed gene sets indicates a reduction in DNA synthesis during grafting, which may promote wound healing for successful graft union formation. By toning down DNA replication, the plant may also reduce the risk of DNA damage and conserve resources for essential graft-healing processes. MCODE module 2 is mainly enriched in nucleic acid and macromolecular metabolism-related biological processes. MCODE module 3 is mainly enriched in the regulation of nutrient response, RNA, and macromolecule modification-related biological processes. Downregulation of nutrient response genes may reallocate resources from nutrient uptake to prioritize graft healing and tissue regeneration. MCODE module 4 is mainly enriched in RNA modification, vacuolar function, and triterpenoid biosynthesis-related biological processes. GLay module 1 is mainly enriched in nucleic acid and triterpenoid metabolism and macromolecule modifications related biological processes. GLay module 2 is mainly enriched in the regulation of cell morphogenesis, macromolecule modification, and nucleic acid phosphodiester bond hydrolysis-related biological processes. GLay module 3 is mainly enriched in RNA modification, nitrogen, and organic compound metabolism-related biological processes.

The GO superclusters identified through REVIGO treemap analysis for pattern D included “RNA modification”, “organic cyclic compound metabolic process”, “regulation of primary metabolic process”, “response to stimulus”, “cell recognition”, “metabolic process”, and “response to stress”. MCODE 1 in pattern D exhibits no significant functional associations between the biological processes identified within the module and the GO superclusters identified. MCODE 2 enriched in “RNA modification”, “nucleic acid phosphodiester bond hydrolysis”, “RNA metabolic process”, “nucleic acid metabolic process”, “nucleobase-containing compound metabolic process”, “macromolecule modification”, “heterocycle metabolic process”, “cellular aromatic compound metabolic process” and “organic cyclic compound metabolic process” contributes to the GO superclusters “RNA modification” and “organic cyclic compound metabolic process” in the REVIGO treemap. Enriched biological processes in MCODE module 3 including “RNA modification”, “nucleic acid phosphodiester bond hydrolysis”, “RNA metabolic process”, “nucleic acid metabolic process”, “macromolecule modification”, “organic cyclic compound metabolic process” and “organic substance metabolic process” contributes to the GO superclusters “RNA modification” and “organic cyclic compound metabolic process” in the REVIGO treemap. MCODE module 4 enriched in “RNA modification”, “RNA metabolic process”, “nucleic acid metabolic process”, “nucleobase-containing compound metabolic process”, “macromolecule modification”, “heterocycle metabolic process”, “NADP biosynthetic process”, “organic cyclic compound metabolic process” and “organic substance metabolic process” are associated with the GO superclusters “RNA modification” and “organic cyclic compound metabolic process” in the REVIGO treemap. This shows the coordinated relationship between the independent MCODE modules in the network of expression pattern D to the GO superclusters.

The REVIGO treemap indicates that GLay module 1 in pattern D enriched in “RNA modification”, “RNA metabolic process”, “nucleic acid phosphodiester bond hydrolysis”, “nucleic acid metabolic process”, “macromolecule modification” and “organic cyclic compound metabolic process” is associated with the “RNA modification” and “organic cyclic compound metabolic process” GO superclusters. GLay module 2 enriched in “nucleic acid phosphodiester bond hydrolysis” and “macromolecule modification” are associated with the GO supercluster “RNA modification” in the REVIGO treemap. GLay module 3 enriched in “RNA modification”, “nucleic acid phosphodiester bond hydrolysis”, “RNA metabolic process”, “nucleic acid metabolic process”, “macromolecule modification”, “nucleobase-containing compound metabolic process”, “heterocycle metabolic process”, “NADP biosynthetic process”, “nitrogen compound metabolic process”, “organic cyclic compound metabolic process” and “organic substance metabolic process” contributes to the GO superclusters “RNA modification” and “organic cyclic compound metabolic process” in the REVIGO treemap. Our analysis highlights the functional association between the REVIGO GO superclusters and enriched GO-BP terms in the GLay modules from pattern D.

***Functional enrichments and hub genes of MCODE modules in pattern D.***

MCODE module 1 in pattern D is enriched in cell growth and division-related biological processes including meiosis and chromosome dynamics (regulation of meiotic cell cycle, meiosis I cell cycle process, meiotic chromosome segregation and homologous recombination), DNA metabolism and cell cycle regulation (DNA replication initiation, DNA metabolic process, regulation of cell cycle and DNA repair) (Supplementary Dataset S5: Sheet 28). In this module, we identified hub genes involved in meiotic chromosome condensation *Nbe.v1.1.chr19g05150* (a homolog of *CAP-D3*), cell division *Nbe.v1.1.chr18g11830* (a homolog of *CYCA3;4*), regulation of sugar signaling *Nbe.v1.1.chr15g00270* (a homolog of *GPT2*) and cell growth and division *Nbe.v1.1.chr05g21840* (a homolog of *TOR1L2*) (Supplementary Dataset S5: Sheet 28). The GO terms enriched in the module reflect suppressed cell division, DNA replication, and sugar signaling processes, following downregulation of these hub genes during grafting. *Nbe.v1.1.chr19g05150* is localized in euchromatin loops and involved in organizing interphase chromatin in centromeric regions and 45S rDNA repeats [360]. It is essential for chromosome condensation during mitosis and meiosis, chromatid resolution, sister chromatid cohesion, and resistance to genotoxic stress [361]. *Nbe.v1.1.chr18g11830* is known to interact with cyclin-dependent kinases (CDKs) to regulate cell cycle and formative cell divisions in root meristem and stomatal development [362,363]. These genes probably involved in cell division and cell cycle processes, may be repressed during grafting, potentially to save energy and cellular resources for wound healing and tissue integration by minimizing unnecessary cell cycle activities. *Nbe.v1.1.chr15g00270* encodes a glucose 6-phosphate/phosphate translocator involved in sugar partitioning between the cytosol and plastids and thus modulates sugar signaling and the response to exogenous sugars [364]. It is also involved in the net import of glucose 6-phosphate (G6P) from the cytosol to the chloroplast and enhances starch synthesis and the oxidative pentose phosphate pathway

[365–367]. During grafting, the downregulation of *GPT2* may be a coordinated strategy to reallocate carbon resources and alter the sugar signaling pathway such that there is limited plastidial G6P import for starch biosynthesis, but rather increase cytosolic sugar availability, which may act as signals and support the energy-intensive processes like wound healing and callus formation at the graft junction. Thus, *GPT2* repression may alter sugar signaling to prioritize regeneration and successful graft union formation.

MCODE module 2 in pattern D is enriched in RNA and nucleic acid metabolism (RNA modification, RNA metabolic process, and nucleic acid phosphodiester bond hydrolysis), macromolecule and compound metabolism (macromolecule modification, cellular aromatic compound metabolic process, organic cyclic compound metabolic process, and cellular nitrogen compound metabolic process), and protein localization and transport (Supplementary Dataset S5: Sheet 28). In this module, we identified hub genes involved in protein-protein interactions *Nbe.v1.1.chr19g27950* (a homolog of TPR-like superfamily protein), chloroplast biogenesis *Nbe.v1.1.chr03g20220* (a homolog of *EMB3103*) and biogenesis of photosynthetic complexes *Nbe.v1.1.chr04g21100* (a homolog of *PDE338*) (Supplementary Dataset S5: Sheet 28). *Nbe.v1.1.chr19g27950* encodes a TPR-like protein known to contain tandem arrays of TPR motifs essential for mediating protein-protein interactions, chaperone activities, and cell cycle regulation [45,368,369]. *Nbe.v1.1.chr03g20220* (a homolog of *EMB3103*) is a member of the PPR repeat superfamily proteins, involved in chloroplast biogenesis, plastid RNA editing, and embryo development [370,371]. *Nbe.v1.1.chr04g21100* (a homolog of *PDE338*) encodes a chloroplast-localized protein that regulates the translation of *Ycf1* by binding to its mRNA. It is involved in the biogenesis of photosynthetic complexes [372,373]. These observations suggest that chloroplast function and photosystem biogenesis are suppressed during graft healing processes.

MCODE module 3 in pattern D is enriched in the regulation of responses to biological processes (regulation of response to extracellular stimulus, regulation of response to nutrient levels, and regulation of response to external stimulus), carbohydrate metabolism (fructose 2,6-bisphosphate metabolic process and fructose metabolic process), radial pattern formation, and defense response (Supplementary Dataset S5: Sheet 28). In this module, we identified a hub gene involved in transcriptional silencing, non-CG DNA methylation, and H3K9 dimethylation *Nbe.v1.1.chr03g42690* (a homolog of *AGDP1*) (Supplementary Dataset S5: Sheet 28). *Nbe.v1.1.chr03g42690* is a heterochromatin-binding protein that binds to H3K9me2 tails and long transposable elements and is involved in DNA methylation pathways and gene regulation [55,56]. In different regions of the gene sequence, DNA methylation has different regulatory effects, including CG methylation, which represses or activates gene expression under normal conditions, whereas non-CG methylation plays a major role in gene repression [374]. DNA methylation can regulate gene expression, which influences plant physiology development and induces phenotypic variation [375,376]. During grafting in tuber mustard, DNA methylation levels were changed and altered the transcription levels of coding genes related to leaf shape development, including *ARF10*, *PIN3*, *IAA20*, and *TPR2,* and small RNAs [377]. DNA methylation was suggested to potentially induce grafting phenotypic variations through the regulation of transcription changes in coding genes and microRNA genes (MIRs). Also, in *A. thaliana* and *S.*

*lycopersicum*, grafting-induced DNA methylation was found to mediate the inheritance of enhanced grafting-induced growth vigor in sexual offspring [61]. Therefore, during grafting, the repression of *AGDP1* may be involved in reducing the establishment of repressive chromatin marks and non-CG methylation to relax heterochromatin formation for greater chromatin accessibility and active expression of genes required for graft healing and integration.

MCODE module 4 in pattern D is enriched in RNA and nucleic acid metabolism biological processes (RNA modification, chloroplast RNA modification, nucleic acid metabolic process, and RNA polyadenylation), NADP biosynthetic and metabolic process (Supplementary Dataset S5: Sheet 28). In this module, we identified hub genes involved in mRNA binding (*Nbe.v1.1.chr07g11510*, a homolog gene encoding ran BP2/NZF zinc finger-like superfamily protein) and regulation of iron deficiency response and silique growth (*Nbe.v1.1.chr02g03250*, a homolog of *NAC100*) (Supplementary Dataset S5: Sheet 28). *Nbe.v1.1.chr02g03250* (a homolog of *NAC100*) modulates gene expression for iron uptake and gibberellin metabolism for silique development and elongation [378]. The grafted plants may prioritize healing and tissue regeneration over nutrient uptake and reproductive development, thereby repressing *NAC100*, which may be a strategic shift in resource allocation from nutrient stress responses and reproductive growth to successful graft union formation and regeneration.

The key features of the remaining 3 nonsignificant (threshold score > 5) MCODE modules in the network of pattern D are briefly summarized below, including the node counts and functional enrichments. These modules were enriched in "glycogen catabolic process" (MCODE 5, 17 nodes and 36 edges), "RNA modification" (MCODE 6, nodes and 52 edges), and "cell recognition" (MCODE 7, 13 nodes and 23 edges). The hub genes in MCODE modules 5–7 are involved in cell wall-degradation and remodeling *Nbe.v1.1.chr19g40800* (a homolog of *BXL6*), mitochondria, and chloroplasts RNA editing *Nbe.v1.1.chr04g22320* (homolog genes encoding PPR repeat superfamily protein) and cell-cell interaction following ubiquitin signaling *Nbe.v1.1.chr02g28600* (a homolog gene encoding Josephin protein-like protein) (Supplementary Dataset S5: Sheet 28). During grafting, the transcriptional downregulation of these genes may potentially limit excessive cell wall degradation and reduce extensive RNA processing and cell signaling or recognition events that are not directly related to the formation of the graft union.

***Functional enrichments and hub genes of GLay modules in pattern D.***

GLay module 1 in pattern D is enriched in RNA and nucleic acid metabolism biological processes (RNA modification, RNA metabolic process, chloroplast RNA modification, and nucleic acid metabolic process), pentacyclic triterpenoid biosynthetic and metabolic process (Supplementary Dataset S6: Sheet 34). In this module, we identified a hub gene involved in the phosphorylation of xylulose, a pentose sugar with a ketone functional group (*Nbe.v1.1.chr03g19170*, a homolog of *XK1*) (Supplementary Dataset S7: Sheet 40). This gene catalyzes the phosphorylation of xylulose, a five-carbon sugar, into xylulose-5-phosphate, a metabolite that enters the pentose phosphate pathway (PPP) and is involved in the biosynthesis of plastidial isoprenoids, NADPH generation for reductive biosynthesis, antioxidant defense, and producing ribose-5-phosphate for nucleotide synthesis [379,380].

The general activity of PPP is crucial for plastidial isoprenoid biosynthesis. Plastidial isoprenoids, such as carotenoids, terpenoids, and other isoprenoids, are essential for photosynthesis, defense, and hormone synthesis [381]. Thus, repression of this gene during grafting may represent a regulatory mechanism to restrict PPP and plastidial isoprenoid biosynthesis to limit the flux of pentose phosphate into the PPP, thereby prioritizing energy and resource allocation for immediate wound healing and tissue regeneration.

GLay module 2 in pattern D is enriched in the regulation of cell morphogenesis, macromolecule modification, and nucleic acid phosphodiester bond hydrolysis, biological processes (Supplementary Dataset S6: Sheet 34). In this module, we identified hub genes involved in microtubule growth (*Nbe.v1.1.chr15g00270*, a homolog of *GPT2*), trichome development (*Nbe.v1.1.chr02g06170*, a homolog gene encoding actin cytoskeleton-regulatory complex pan-like protein), cytokinesis (*Nbe.v1.1.chr02g18210*, a homolog of *SYP112*), and DNA replication (*Nbe.v1.1.chr01g35640*, a homolog of *ORC5*) (Supplementary Dataset S7: Sheet 40). *Nbe.v1.1.chr02g06170* has been identified to share conserved amino acid sequences REERVQMKL with branchless trichomes (BLT) proteins, which are involved in cytoskeletal organization for trichome branching [382,383], intracellular transport and communication [384], cell growth and division [385], and response to environmental stresses [386–389]. Trichome development and related cytoskeletal branching are energy-intensive and may not be essential for the wound healing processes; therefore, repressing this gene during grafting may suppress analogous branching, reduce cytoskeletal remodeling, and prevent premature cell differentiation. *Nbe.v1.1.chr02g18210* (a homolog of *SYP112*) is a plasma membrane-localized SNARE protein functionally equivalent to *KNOLLE* (cytokinesis-specific syntaxin protein), involved in membrane fusion during cytokinesis, and is rapidly degraded after cell division [386–389]. During grafting, repression of this gene is likely important to limit unnecessary or excessive cell division and membrane trafficking at the graft site, thereby ensuring wound healing and tissue integration-focused metabolic processes.

GLay module 3 in pattern D is enriched in RNA and nucleic acid metabolism biological processes (RNA and mRNA modification, RNA metabolic process, and RNA-templated transcription), post-translational modifications (macromolecule modification, peptidyl-lysine demethylation, and protein O-linked fucosylation), cellular response (response to nutrient levels and extracellular stimulus), and organic compound metabolism (Supplementary Dataset S6: Sheet 34). In this module, we identified hub genes involved in phosphate ion transmembrane transport and response to salt stress (*Nbe.v1.1.chr11g31030*, a homolog of *PHT3;1*), transcriptional silencing, non-CG DNA methylation, and H3K9 dimethylation (*Nbe.v1.1.chr03g42690*, a homolog of *AGDP1*) and intercellular transport and root radial pattern formation (*Nbe.v1.1.chr12g03720*, a homolog of short-root interacting embryonic lethal, *SIEL*) (Supplementary Dataset S7: Sheet 40). *Nbe.v1.1.chr11g31030* (a homolog of *PHT3;1*) encodes a mitochondrial phosphate transporter essential for importing phosphate into the mitochondrial matrix, which is essential for ATP production, cell energy metabolism, and specifically modulates responses to salt stress by regulating cellular energy status and ROS production [69,390,391]. This gene, during grafting, may be temporally upregulated for wound healing and rapidly repressed to reduce ATP synthesis and limit energy-intensive processes. Also, the repression of the gene may prevent the increased mitochondrial activity and ROS production, which may disrupt the redox balance at the graft interface. *Nbe.v1.1.chr12g03720* (a homolog of *SIEL*) encodes a SHORT-ROOT (SHR) transcription factor, which interacts with SHR and other non-cell-autonomous proteins and facilitates their movement between cells via plasmodesmata, essential for proper root patterning and development [392]. Repression of this gene during grafting may be significant in preventing extensive intercellular signaling and the activation of root-specific development that is not required for graft healing.

Several hub genes and biological processes overlapped between the MCODE and GLay modules in pattern D. For example, *Nbe.v1.1.chr15g00270* (a homolog of *GPT2*) was identified in both MCODE module 1 and GLay module 2, highlighting its potential role in regulating cell morphogenesis and sugar signaling. Similarly, *Nbe.v1.1.chr03g42690* (a homolog of *AGDP1*) was present in both MCODE module 3 and GLay module 1, suggesting its involvement in DNA methylation processes.

**Pattern E**

***Gene profiles of pattern E.***

Pattern E genes, compared to the starting point (intact), had lower expression levels in the second state (Ave 1–7 DAG) only after grafting. It had the lowest number of genes out of the six patterns (51 genes). Genes in expression Pattern E comprise diverse functional classes, including cell wall and membrane proteins (o-glycosyl hydrolase and TRICHOME BIREFRINGENCE-LIKE 13 and 14 proteins), transcription factors (*NAC75*, *NAC78* homeobox-leucine zipper, *MYB96,* and cycling DOF factor 2), and lignin biosynthesis (ferulic acid 5-hydroxylase). In the Bayesian gene regulatory network of pattern E, we identified genes with known functions in plant growth and development such as *NAC075* which is a known repressor of flowering [393], a negative regulator of leaf senescence by deterring reactive oxygen species accumulation [394] and transcriptionally associated with secondary cell wall reconstitution as a chimeric activator and modulates secondary cell wall composition including increased glucose, xylose and lower lignin content [395]. *NAC083* negatively regulates xylem vessel formation [396,397] and inhibits transcriptional activation activity of *ATAF2,* and this interaction results in a negative regulation of leaf senescence [398]. *RELATED TO AP2.7* (*RAP2.7*), a member of the AP2/ERF family transcription factors, involved in the regulation of flowering and innate immunity, and its transcription is positively regulated by the meristem-defective (*MDF*) gene in response to wounding [399] (Supplementary Dataset S8: Sheet 47). Repression of *NAC075*, *NAC083,* and *RAP2.7* may ensure the proper regulation of cell wall component synthesis, efficient vascular differentiation, and immune-related growth inhibition.

In this network for pattern E, 3 significant modules (threshold score > 5) were identified using the MCODE algorithm, and 3 modules were identified using the GLay community clustering algorithm. MCODE module 1 is mainly enriched in chloroplast-nucleus and hormone signaling and secondary metabolism-related biological processes. MCODE module 2 is mainly enriched in the regulation of histone acetylation and modifications, seed and floral development-related biological processes. MCODE module 3 is mainly enriched in the regulation of lipid, wax, and lignin biosynthesis and seed development-related biological processes. GLay module 1 is mainly enriched in hormonal signaling, lipid metabolism, and regulation of seed development and dormancy-related biological processes. GLay module 2 is mainly enriched in the regulation of histone modifications, apocarotenoid, and ABA biosynthesis-related biological processes. GLay module 3 is mainly enriched in auxin (indole-3-acetic acid) biosynthesis and metabolism-related biological processes.

The GO superclusters identified through REVIGO treemap analysis for pattern E included "auxin biosynthetic process" and "histone H3−K9 acetylation". MCODE module 1 in pattern E enriched in "auxin biosynthetic process", "auxin metabolic process" is associated with the "auxin biosynthetic process" GO supercluster in the REVIGO treemap. In the REVIGO treemap, MCODE module 2 enriched in "regulation of seed development", "histone H3-K9 acetylation", and "histone H3-K9 modification" contributed to the "auxin biosynthetic process" and "histone H3-K9 acetylation" GO superclusters. The enriched terms in MCODE module 3, including "positive regulation of wax biosynthetic process" and "regulation of wax biosynthetic process," are associated with the "auxin biosynthetic process" GO supercluster in the REVIGO treemap. This illustrates the contribution of the distinct molecular functions of the MCODE modules in pattern E to the broader biological processes in the REVIGO treemap.

GLay module 1 in pattern E enriched in "regulation of seed development", "positive regulation of wax biosynthetic process", and "regulation of wax biosynthetic process" are associated with the broader GO supercluster related to "auxin biosynthetic process". GLay module 2 enriched in "histone H3-K9 acetylation" and "histone H3-K9 modification" contributes to the GO superclusters "histone H3-K9 acetylation". GLay module 3 enriched in "auxin biosynthetic process" contributes to the GO supercluster "auxin biosynthetic process". These results indicate a potential functional link between the biological processes identified within the GLay modules and GO superclusters from pattern E.

***Functional enrichments and hub genes of MCODE modules in pattern E.***

MCODE module 1 in pattern E is enriched in the following biological processes, including chloroplast-nucleus signaling pathway, apocarotenoid biosynthetic process, tertiary alcohol biosynthetic process, abscisic acid biosynthetic process, auxin biosynthetic process, olefinic compound biosynthetic process, and sesquiterpenoid biosynthetic process (Supplementary Dataset S5: Sheet 29). In this module, we identified a hub gene involved in the stabilization of NADH dehydrogenase 1 transcript (*Nbe.v1.1.chr05g01150*, a homolog of *PPR19*) (Supplementary Dataset S5: Sheet 29). *Nbe.v1.1.chr05g01150* is involved in the stabilization of NADH dehydrogenase 1 (nad1) transcripts for proper mitochondrial function and improved development, including normal seed development, increased seed yield, improved seed germination, and growth [400]. This gene, during grafting, may be repressed to reduce mitochondrial activity and these energy-intensive developmental processes, thereby reallocating metabolic resources toward wound healing.

MCODE module 2 in pattern E is enriched in the following biological processes, including histone acetylation, regulation of seed growth, phosphate ion homeostasis, and specification of floral organ identity (Supplementary Dataset S5: Sheet 29). In this module, we identified a hub gene involved in plant immunity and senescence (*Nbe.v1.1.chr02g21910*, a homolog of *SAG101*) (Supplementary Dataset S5: Sheet 29). *Nbe.v1.1.chr02g21910* encodes a lipase-like protein involved in plant immunity and senescence. *SAG101* forms a ternary complex with *EDS1* and *PAD4,* which regulates the subcellular localization of *EDS1* for effective immune responses against pathogens like Turnip crinkle virus (TCV) and *Pseudomonas syringae* [401,402]. Repression of *SAG101* during grafting may be necessary to prevent premature activation of immune and senescence pathways that could lead to excessive cell death or defense signaling at the graft junction.

MCODE module 3 in pattern E is enriched in the following biological processes, including regulation of wax biosynthetic process, regulation of abscisic acid biosynthetic process, regulation of seed dormancy process, positive regulation of lipid biosynthetic process, and lignin biosynthetic process (Supplementary Dataset S5: Sheet 29). In this module, we identified a hub gene that may be involved in the synthesis and deposition of secondary wall cellulose (*Nbe.v1.1.chr15g07360*, a homolog of *TBL41*) (Supplementary Dataset S5: Sheet 29). *Nbe.v1.1.chr15g07360* contains two conserved domains, the TBL and DUF231, involved in the o-acetylation of cell wall polymers [403,404]. Thus, *TBL41,* a member of the other 46 TBL genes [404], may be involved in the O-acetylation of cell wall polymers and pectin esterification [405]. These modifications potentially contribute to increased rigidity and structural integrity of the secondary cell wall, which are normally beneficial for mature tissues but may hinder the dynamic processes required for successful graft union formation. During grafting, *TBL41* may be repressed to reduce the excessive cell wall stiffening and reinforcement, which may hinder cell-to-cell adhesion, tissue remodeling, and vascular reconnection at the graft interface.

***Functional enrichments and hub genes of GLay modules in pattern E.***

GLay module 1 in pattern E is enriched in the following biological processes, including positive regulation of wax biosynthetic process, regulation of wax biosynthetic process, regulation of abscisic acid biosynthetic process, regulation of seed dormancy process, and positive regulation of lipid biosynthetic process (Supplementary Dataset S6: Sheet 35). In this module, we identified hub genes involved in stabilization of NADH dehydrogenase 1 transcripts (*Nbe.v1.1.chr05g01150*, a homolog of *PPR19*), mitochondrial functional regulation (*Nbe.v1.1.chr06g38370*, a homolog of *RBL12*), and RNase BN activity (tRNA-processing ribonuclease BN) (*Nbe.v1.1.chr15g37200*, a homolog gene encoding tRNA-processing ribonuclease BN) (Supplementary Dataset S7: Sheet 41). *Nbe.v1.1.chr06g38370* (a homolog of *RBL12*) encodes a serine protease class of mitochondrion-located rhomboid-like protein involved in intramembrane proteolysis and protein proteolytic cleavage, essential for cellular functions and developmental processes critical for regulating the turnover and activity of membrane proteins that influence various developmental and stress-responsive pathways [58]. Hence, repression of *Nbe.v1.1.chr06g38370* during grafting may be necessary to modulate mitochondrial activity and excessive proteolysis that could disrupt cellular homeostasis during graft healing. *Nbe.v1.1.chr15g37200* is involved in protein synthesis following the processing of precursor transfer RNAs (tRNAs), maturation of tRNAs into their mature folds and forms, and cellular function [406,407]. Downregulation of this tRNA processing-related gene may suppress non-essential translation processes and protein synthesis, thereby reallocating resources toward graft healing processes.

GLay module 2 in pattern E is enriched in the following biological processes, including histone H3-K9 acetylation, histone H3-K9 modification, chloroplast-nucleus signaling pathway, tertiary alcohol biosynthetic process, apocarotenoid biosynthetic process, and abscisic acid biosynthetic process (Supplementary Dataset S6: Sheet 35). In this module, we identified a hub gene involved in nitrate response and signaling (*Nbe.v1.1.chr17g06980*, a homolog of *NODULE INCEPTION-LIKE PROTEIN 4, NLP4*) (Supplementary Dataset S7: Sheet 5). *Nbe.v1.1.chr17g06980* binds to the nitrate-responsive cis-element (NRE) and modulates the expression of nitrate-responsive genes involved in the nitrate signaling pathways, nitrate uptake and transport (nitrate transporter 1.1, *NRT1.1*), and assimilation (nitrate reductase 1 and 2; *NIA1* and *NIA2*) for plant growth and development [408,409], including root cell division [410]. Repression of *Nbe.v1.1.chr17g06980* may temporarily shift metabolic priorities away from nutrient assimilation toward tissue regeneration.

GLay module 3 in pattern E is enriched in the following biological processes, including the indoleacetic acid metabolic process, indoleacetic acid biosynthetic process, and auxin biosynthetic process (Supplementary Dataset S6: Sheet 35). In this module, we identified hub genes involved in protein methylation (*Nbe.v1.1.chr03g40930*, a homolog of putative methyltransferase family protein), *Nbe.v1.1.chr06g17730* (a homolog of PPR repeat superfamily protein) and *Nbe.v1.1.chr16g28330* (a homolog of hypothetical protein) (Supplementary Dataset S7: Sheet 41). *Nbe.v1.1.chr03g40930* encodes a methyltransferase enzyme that catalyzes the transfer of methyl groups to DNA, RNA, proteins, and small molecule substrates essential for regulating gene expression, protein function, and cellular signaling [411,412]. Grafting potentially triggers extensive transcriptional reprogramming and cell fate changes such that excessive methyltransferase activity could lead to inappropriate gene silencing and disruption of the expression of genes necessary for graft-related processes such as cell dedifferentiation, proliferation, vascular regeneration, and hormone signaling. Thus, repression of *Nbe.v1.1.chr03g40930* may ensure the proper regulation and maintenance of epigenetic plasticity to enhance the activation of grafting-related genes.

In pattern E, *Nbe.v1.1.chr05g01150* (a homolog of *PPR19*) was identified as an overlapping hub gene in both MCODE module 1 and GLay module 1. The methods revealed significant enrichment for biological processes related to seed development, wax, and abscisic acid biosynthesis in MCODE module 3 and GLay module 1, indicating core modulators involved in hormone signaling, seed dormancy, and stress adaptation. Additional overlap was observed in processes related to histone H3-K9 acetylation/modification and the chloroplast-nucleus signaling pathway, captured in both MCODE module 2 and GLay module 2.

**Pattern F**

***Gene profiles of pattern F.***

Pattern F genes, compared to the starting point (intact), had lower expression levels in the third state (Ave 14–28 DAG) only after grafting. Genes in expression pattern D comprise diverse functional classes, including transcription factors (transcription termination factor family protein, GATA transcription factor 11 and transcription factor IIIB), protein kinases (protein kinase superfamily protein, PB1 domain-containing protein tyrosine kinase and mitogen-activated protein kinase 3), membrane transporters (K+ uptake transporter 3, potassium channel protein and potassium transporter 1) and DNA and RNA Processing (RNA polymerase sigma subunit 2, ribonuclease P protein subunit P38-like protein and DNA glycosylase superfamily protein).

In this network for pattern F, 4 significant modules (threshold score > 5) were identified using the MCODE algorithm, and 4 modules were identified using the GLay community clustering algorithm. MCODE module 1 is mainly enriched in RNA processing, modification, and nucleic acid metabolism-related biological processes. MCODE module 2 has no gene ontology enrichment output due to the limited number of nodes in the module. MCODE module 3 is mainly enriched in cysteine catabolism, meristematic phase transition, homogalacturonan dynamics, and triterpenoid biosynthesis-related biological processes. MCODE module 4 is mainly enriched in nucleic acid and signal-dependent metabolism and genome stability-related biological processes. GLay module 1 is mainly enriched in RNA modification, mRNA, nucleic acid metabolism, and meristematic phase transitions related biological processes. GLay module 3 is mainly enriched in response to disaccharides and sucrose, regulation of DNA replication, and triterpenoid biosynthesis-related biological processes. GLay module 4 is mainly enriched in the regulation of RNA metabolism, macromolecular modification, and cell cycle progression-related biological processes.

The GO superclusters identified through REVIGO treemap analysis for pattern F revealed interconnected and conserved biological processes including “RNA modification”, “regulation of intracellular signal transduction”, “organic cyclic compound metabolic process”, “cellular response to leucine”, “ubiquitin−dependent protein catabolic process via the N−end rule pathway”, “male meiosis I” and “potassium ion transmembrane transport”. Further enrichment analysis using MCODE and GLay modules revealed significant associations between these superclusters and module-specific biological processes. MCODE module 1 in pattern F enriched in “RNA modification”, “macromolecule modification”, “nucleic acid metabolic process”, “organic cyclic compound metabolic process”, “response to sucrose”, and “potassium ion transmembrane transport” is associated with the GO superclusters in the REVIGO treemap. MCODE module 3 enriched in “regulation of timing of transition from vegetative to reproductive phase” and “regulation of development heterochronic” is associated with the GO superclusters “regulation of intracellular signal transduction” in the REVIGO treemap. MCODE module 4 enriched in “nucleic acid metabolic process”, “histone H2A ubiquitination”, “regulation of lip catabolic process” “organic cyclic compound metabolic process”, “cellular response to leucine” and “ubiquitin−dependent protein catabolic process via the N−end rule pathway” and “male meiosis I” is associated with all the GO supercluster in the REVIGO treemap except “potassium ion transmembrane transport”. This reflects the association of the distinct biological processes in the MCODE modules in the regulatory network of pattern F with the broader biological processes presented in the REVIGO treemap.

GLay module 1 in pattern F enriched in “RNA modification”, “macromolecule modification”, “mRNA modification”, “regulation of intracellular signal transduction” is associated with “RNA modification”, “organic cyclic compound metabolic process”, and “ubiquitin−dependent protein catabolic process via the N−end rule pathway” GO superclusters in the REVIGO treemap. GLay module 3 in pattern F enriched in “response to sucrose" is associated with the "cellular response to leucine" GO supercluster in the REVIGO treemap. GLay module 4 in pattern F enriched in "RNA modification", "macromolecule modification", and "macromolecule metabolic process" is associated with the "RNA modification" and "organic cyclic compound metabolic process" GO superclusters in the REVIGO treemap. The observed overlap between the biological processes in the GLay modules and the GO superclusters provides valuable insight into the integrated nature of these processes.

***Functional enrichments and hub genes of MCODE modules in pattern F.***

MCODE module 1 in pattern F is enriched in the following biological processes, including RNA and mRNA modification, nucleic acid phosphodiester bond hydrolysis, nucleic acid metabolic process, mitochondrial RNA modification and metabolic process, nucleobase-containing compound metabolic process, and macromolecule modification (Supplementary Dataset S5: Sheet 30). In this module, we identified hub genes that may be involved in the regulation of mitochondrial and chloroplast gene expression (*Nbe.v1.1.chr13g15520*, a homolog of *MTERF19*) and phospholipid biosynthetic process (*Nbe.v1.1.chr13g42810*, a homolog gene encoding alkaline-phosphatase-like family protein) (Supplementary Dataset S5: Sheet 30). *Nbe.v1.1.chr13g15520* regulates mitochondrial and chloroplast gene expression [413–415]. This family of mitochondrial genes expressed in plants is suggested to be vital for photosynthesis, cellular respiration, and stress resistance [416–418]. Since chloroplast-related processes like photosynthesis are non-essential during graft healing, repressing *Nbe.v1.1.chr13g15520* could reallocate on-demand resources toward the immediate demands of wound healing. *Nbe.v1.1.chr13g42810* is implicated in hydrolyzing organic phosphates into inorganic phosphate (Pi) under Pi-deficient conditions for improved phosphorus uptake and plant growth [419,420]. This gene may be repressed during grafting to suppress nutrient assimilation during wound healing and union formation.

MCODE module 2 in pattern F had no gene ontology enrichment output due to the limited number of nodes in the module. In this module, we identified hub genes that may be involved in chlorophyll biosynthesis and photomorphogenesis (*Nbe.v1.1.chr11g36450*, a homolog of *GATA11*) and regulating homologous recombination (*Nbe.v1.1.chr16g03590*, a homolog of *FLIP*) (Supplementary Dataset S5: Sheet 30). *Nbe.v1.1.chr11g36450* belongs to the GATA family of transcription factors involved in regulating various aspects of growth and development. Specifically, *GATA11* plays a role in the regulation of light-responsive genes and may be involved in processes such as chlorophyll biosynthesis, chloroplast development, and photomorphogenesis [421]. Downregulating *GATA11* during grafting is essential for prioritizing graft union formation processes over those related to light-mediated development and photosynthetic capacity. *Nbe.v1.1.chr16g03590* (a homolog of *FLIP*) is involved in regulating homologous recombination through a conserved complex formation with FIDGETIN-LIKE-1 and interaction with recombinases RAD51 and DMC1 to limit the number and distribution of crossover events during meiosis. This regulation ensures proper chromosome segregation and genetic diversity during sexual reproduction [422]. Homologous recombination comprises a series of interrelated pathways that function in DNA repair and provide critical support for DNA replication [423]. However, elevated activity during grafting could lead to unwanted chromosomal rearrangements and instability in rapidly dividing cells at the wound site. Repressing this gene during grafting may minimize the risk of genomic instability, which may disrupt the precision of cellular activities required for graft healing.

MCODE module 3 in pattern F is enriched in the following biological processes, including cysteine catabolic process, regulation of timing of meristematic phase transition, regulation of timing of transition from vegetative to reproductive phase, homogalacturonan metabolic and biosynthetic process, pentacyclic triterpenoid biosynthetic and metabolic process (Supplementary Dataset S5: Sheet 30). In this module, we identified a hub gene that may be involved in light signaling and photomorphogenesis (*Nbe.v1.1.chr17g39110*, a homolog of *GT2*) (Supplementary Dataset S5: Sheet 30). *Nbe.v1.1.chr17g39110* binds to GT elements in the promoters of light-regulated genes, playing a crucial role in light signaling and photomorphogenesis [424,425]. This may indicate that light response behavior is suppressed during grafting.

MCODE module 4 in pattern F is enriched in the following biological processes, including nucleic acid metabolic process, nucleobase-containing compound metabolic process, mitochondrial DNA metabolic process, regulation of transposition, ubiquitin-dependent protein catabolic process via the N-end rule pathway, negative regulation of TOR signaling, plasmid maintenance, chromosome organization, and regulation of intracellular signal transduction (Supplementary Dataset S5: Sheet 30). In this module, we identified a hub gene, *Nbe.v1.1.chr05g08230* (a homolog of *PRT6*) that may be involved in protein degradation and seed germination (Supplementary Dataset S5: Sheet 30). *Nbe.v1.1.chr05g08230* encodes a ubiquitin ligase belonging to the N-end rule pathway, which mediates the ubiquitin- and proteasome-dependent turnover of proteins with specific amino-terminal residues [426]. It specifically targets proteins with an amino-terminal arginine residue for degradation [426]. *PRT6* has also been shown to be involved in regulating seed germination, seedling establishment, responses to ABA, and sugar sensitivity [427]. The downregulation of this gene during grafting may be essential to reduce its excessive degradation of proteins that may be crucial for wound healing, since such protein turnover may disrupt the hormonal and metabolic balance required for successful graft union formation.

Additionally, the nonsignificant MCODE module (threshold score > 5) (MCODE 5, 17 nodes and 36 edges) is suggested to be functionally centered on protein quality control, as evidenced by its enrichment in “protein unfolding”, “positive regulation of translation”, and “circumnutation” with the seed node, *Nbe.v1.1.chr12g43890* (a homolog gene encoding a protein kinase superfamily protein). The hub gene identified in the module was *Nbe.v1.1.chr10g02200* (a homolog gene encoding AAA-type ATPase family protein) (Supplementary Dataset S5: Sheet 30).

***Functional enrichments and hub genes of GLay modules in pattern F.***

GLay module 1 in pattern F is enriched in the following biological processes, including positive regulation of wax biosynthetic process, regulation of wax biosynthetic process, regulation of abscisic acid biosynthetic process, regulation of seed dormancy process, and positive regulation of lipid biosynthetic process (Supplementary Dataset S6: Sheet 36). In this module, we identified hub genes involved in protein degradation and seed germination

(*Nbe.v1.1.chr05g08230*, a homolog of *PRT6*), mitochondrial RNA editing and delayed development (*Nbe.v1.1.chr05g31860*, a homolog of slow growth 1 gene, *SLG1*) and regulation of mitochondrial and chloroplast gene expression (*Nbe.v1.1.chr03g13210*, a homolog of *MTERF19*) (Supplementary Dataset S7: Sheet 42). *Nbe.v1.1.chr05g31860* is involved in the editing of mitochondrial transcript nad3, a subunit of complex I in the electron transport chain, essential for the proper function of mitochondrial genes and enhanced response to ABA and drought stress [428]. However, functional disruption of *SLG1* has been reported to result in slow growth and delayed development [428]. Downregulation of this gene during grafting may moderate mitochondrial activity and repress stress signaling by preventing the overactivation of ABA-mediated stress pathways that are not immediately beneficial for graft healing.

GLay module 3 in pattern F is enriched in the following biological processes, including response to disaccharide, response to sucrose, pre-replicative complex assembly involved in nuclear cell cycle DNA replication, priming of cellular response to stress, terpenoid biosynthetic process, and phosphorylation (Supplementary Dataset S6: Sheet 36). In this module, we identified hub genes involved in the phosphorylation of xylulose, a pentose sugar with a ketone functional group (*Nbe.v1.1.chr03g19170*, a homolog of *XK1*) and expression of plastid-encoded tRNAs (*Nbe.v1.1.chr07g12190*, a homolog of RNA-polymerase sigma subunit 2 gene, *SIG2*) (Supplementary Dataset S7: Sheet 6). *Nbe.v1.1.chr07g12190* is involved in the expression of plastid-encoded tRNAs and the repression of nucleus-encoded T7 phage-type RNA polymerase (NEP)-dependent genes, which is crucial for chloroplast development and function [429,430]. *SIG2* also impacts root elongation, and its expression has been identified to contribute to an inductive light signaling pathway by coordinating components between the nucleus and plastids [431]. This gene may be repressed during grafting to suppress light-dependent developmental processes, including energy-intensive processes, chloroplast biogenesis, and photomorphogenesis, that may compete with the regenerative demands during graft healing.

GLay module 4 in pattern F is enriched in the following biological processes, including RNA modification, macromolecule modification, nucleic acid phosphodiester bond hydrolysis, RNA metabolic process, nitrogen compound metabolic process, heterocycle metabolic process, cellular aromatic compound metabolic process, macromolecule metabolic process, and organic cyclic compound metabolic process (Supplementary Dataset S6: Sheet 36). In this module, we identified a hub gene involved in regulating homologous recombination (*Nbe.v1.1.chr16g03590*, a homolog of *FLIP*) whose repression during grafting may minimize the incidence of genomic instability (Supplementary Dataset S7: Sheet 42).

Following module detection analysis, MCODE module 2 and GLay module 4 exhibited overlap hub genes, such as *Nbe.v1.1.chr16g03590* (a homolog of *FLIP*), which regulates homologous recombination [422], and functional enrichment. The detection of *FLIP* in both modules suggests a conserved repression of this gene during grafting to ensure stable genome integrity. These complementary modules illustrate the efficiency of the distinct network clustering approaches in identifying key regulatory complexes in the network of pattern F.
