## Supplementary figures and images for "A network analysis for the identification of gene modules in the transcriptome during *Nicotiana benthamiana* interfamily grafting"

### OpokuAgyemang_et_al_Supplementary Figure S1.pdf

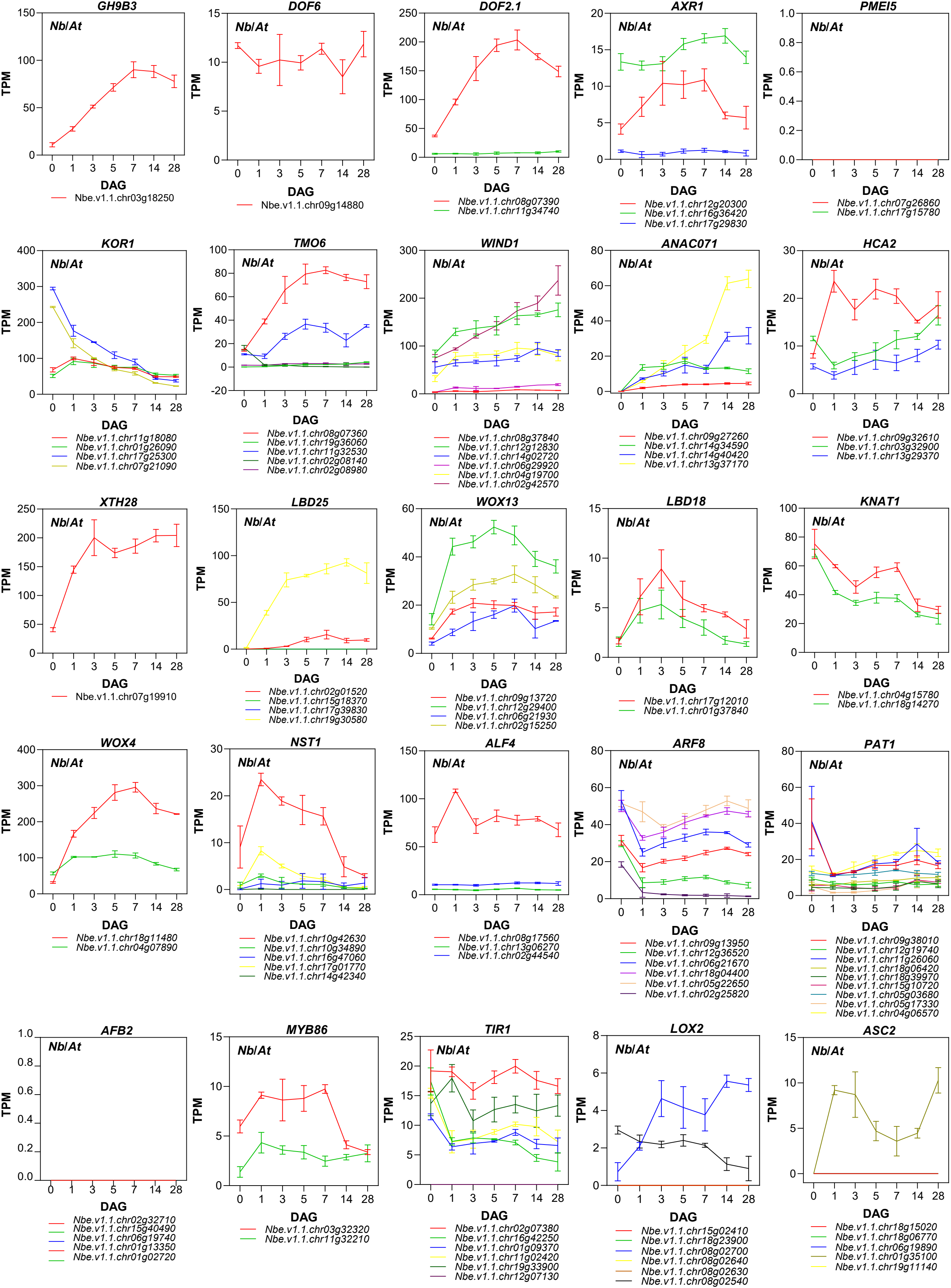
